# Gut microbe-derived short-chain fatty acids regulate joint inflammation and activation of infiltrating and resident myeloid cells after alphavirus infection

**DOI:** 10.64898/2025.12.03.691668

**Authors:** Fang R. Zhao, Maksim Kleverov, Emma S. Winkler, Russell B. Williams, Hana Janova, Lindsay Droit, Leran Wang, Ting-ting Li, Leah Heath, Ana Jung, Matthias Mack, Megan T. Baldridge, Thaddeus S. Stappenbeck, Larissa B. Thackray, Chyi-Song Hsieh, Scott A. Handley, Chun-Jun Guo, Michael A. Fischbach, Maxim N. Artyomov, Michael S. Diamond

**Author notes:** Correspondence and Lead Contact: Michael S. Diamond, M.D., Ph.D.

## Abstract

Although oral antibiotics can predispose to joint inflammation, this phenomenon remains poorly understood. Here, we leverage mouse models of alphavirus-induced arthritis to investigate the roles of gut commensals, metabolites, and host immune mechanisms in promoting musculoskeletal inflammation. Mice treated with a short course of oral antibiotics exhibited worsened arthritis after chikungunya (CHIKV) or Mayaro virus infections. This phenotype was associated with loss of short chain fatty acids (SCFA), greater intestinal permeability, and activation of gut-associated immune cells, and required TLR4 signaling, MyD88 expression, monocytes, antigen-specific and bystander CD4^+^ T cells, and pro-inflammatory cytokines. Administration of exogenous SCFA or colonization of mice with bacterial species that generate SCFA mitigated CHIKV-induced joint inflammation. Single-cell RNA sequencing revealed that gut-derived SCFA ameliorate the inflammatory phenotype of synovial CD4^+^ T cells, infiltrating monocytes, and resident osteoclast-like cells. Thus, antibiotic-triggered gut dysbiosis exacerbates alphavirus arthritis by shaping the inflammatory profile of both infiltrating and resident immune cells in joint tissues.

## INTRODUCTION

Antibiotic usage is linked to an increased risk for onset or relapse of inflammatory arthritis, including rheumatoid arthritis and juvenile idiopathic arthritis ^1–6^. Although a gut-joint linkage has been described ^7–10^, the basis for this increased risk with antibiotics use is not well understood, particularly how joint inflammation is regulated by interactions between gut microbes, mucosal barriers, and immune cells. Less is known about the effects of antibiotics and the commensal intestinal microbiota on inflammation in the context of viral arthritis or other pathogens that infect joint tissues.

Chikungunya virus (CHIKV) is a mosquito-transmitted alphavirus that causes epidemics of debilitating inflammatory arthritis, affecting millions of people globally. The resurgence of CHIKV in the Indian Ocean, Africa, and Asia during 2025 and autochthonous cases in New York has triggered global concerns about the possibility of another widespread epidemic. Three other related alphaviruses (Mayaro [MAYV], Ross River [RRV], and O’nyong-nyong [ONNV] viruses) also cause infection and musculoskeletal disease in the Americas, Oceania, and Africa, respectively^11^. CHIKV infection results in an acute febrile illness in humans, with viremia, rash, muscle pain, and severe joint inflammation, and is associated with elevated systemic levels of cytokines and chemokines including type I interferons (IFNs), IL-6, CCL2, CXCL9, CXCL10, IL-17A, and IFN-γ ^12–17^. This viral arthritis clinically mimics rheumatoid arthritis, can persist for months to years after initial infection, and can cause bone erosions, subchondral defects, and radiographic progression of joint damage ^18–21^.

In C57BL/6 mice, subcutaneous inoculation of CHIKV leads to arthritis, tenosynovitis, and myositis, like that described in humans ^22–24^. Inflammation is associated with tissue infiltration of monocytes, neutrophils, and T cells. However, there often is a discordance between viral burden and the severity of arthritis, suggesting that inflammation and joint pathology are not directly related to the amount of virus in joint-associated tissues. Indeed, CHIKV infection of monocyte-depleted animals or *Rag1^-/-^* mice lacking mature T and B cells leads to persistent local infection yet improved joint swelling compared to control-treated or wild-type mice ^22,25^. Moreover, mice lacking CD4^+^ T cells also have attenuated joint swelling after CHIKV infection without substantive effects on viral burden ^26,27^. These data highlight the differential contributions of immune cells in restricting CHIKV infection and causing musculoskeletal disease.

Here, we use mouse models of alphavirus arthritis to investigate how gut commensals, their metabolites, and host immune responses promote joint inflammation in the setting of antibiotic-mediated dysbiosis. We show that intestinal dysbiosis resulting from a short pulse of minimally absorbed oral antibiotics results in increased gastrointestinal (GI) tract permeability due to loss of specific short-chain fatty acids (SCFA) produced by the commensal intestinal microbiota. This phenotype was associated with enhanced foot swelling and joint infiltration of monocytes and CD4^+^ T cells after CHIKV infection. The increased inflammation was dependent on gut-associated immune cells and required expression of toll-like receptor (TLR)4 and MyD88, with signaling contributions from intestinal epithelial cells. The inflammatory phenotypes in antibiotic-treated mice were improved by treatment with neutralizing antibodies against IL-17A, TNF, or IFN-γ, and not observed in alphavirus-infected germ-free (GF) mice that lack commensal microbes or their metabolites. Reconstitution with single bacterial species that produce SCFA, but not congenic bacterial mutants lacking this ability, or oral administration of specific SCFA reversed the gut permeability and CHIKV-induced arthritis phenotypes seen in antibiotic-treated mice. Finally, SCFA supplementation reversed the inflammatory phenotypes of CD4^+^ T cells, monocytes, and osteoclast-like cells triggered by antibiotic treatment. Overall, our data suggest that antibiotic-induced disruption of the intestinal microbiota results in loss of microbe-derived SCFA that drive host inflammatory cascades in joint-associated tissues by shaping the phenotypes of both infiltrating and resident immune cells.

## RESULTS

### Perturbation of the intestinal microbiota exacerbates CHIKV-induced arthritis

A mouse model of CHIKV-induced musculoskeletal disease recapitulates many features of human disease, including joint swelling, tenosynovitis, and myositis ^22–24^. To begin to evaluate the effect of the microbiome on CHIKV arthritis, we pulsed mice with a short 3-day course of oral ampicillin and vancomycin treatment (**Fig 1A**). This treatment alters the bacterial community structure in the GI tract by depleting bacteria primarily from the *Bacteroidetes* phylum, leading to an overrepresentation of *Proteobacteria* and *Firmicutes* (**Fig 1B-D and S1A-C**). While treatment with either vancomycin or ampicillin alone increased foot swelling after CHIKV infection compared to control (water)-treated mice, administration of the two antibiotics in combination resulted in a larger phenotype (**Fig 1E-G**); consequently, we used the ampicillin and vancomycin (AV) combination treatment for the remainder of our microbiota perturbation studies. The AV-treatment effect on joint inflammation lasted for at least 8 weeks after the cessation of antibiotics (**Fig S1D-G**). Increased joint swelling was also observed in AV-treated mice after infection with MAYV, a related arthritogenic alphavirus (**Fig S1H**).

**Figure 1.**
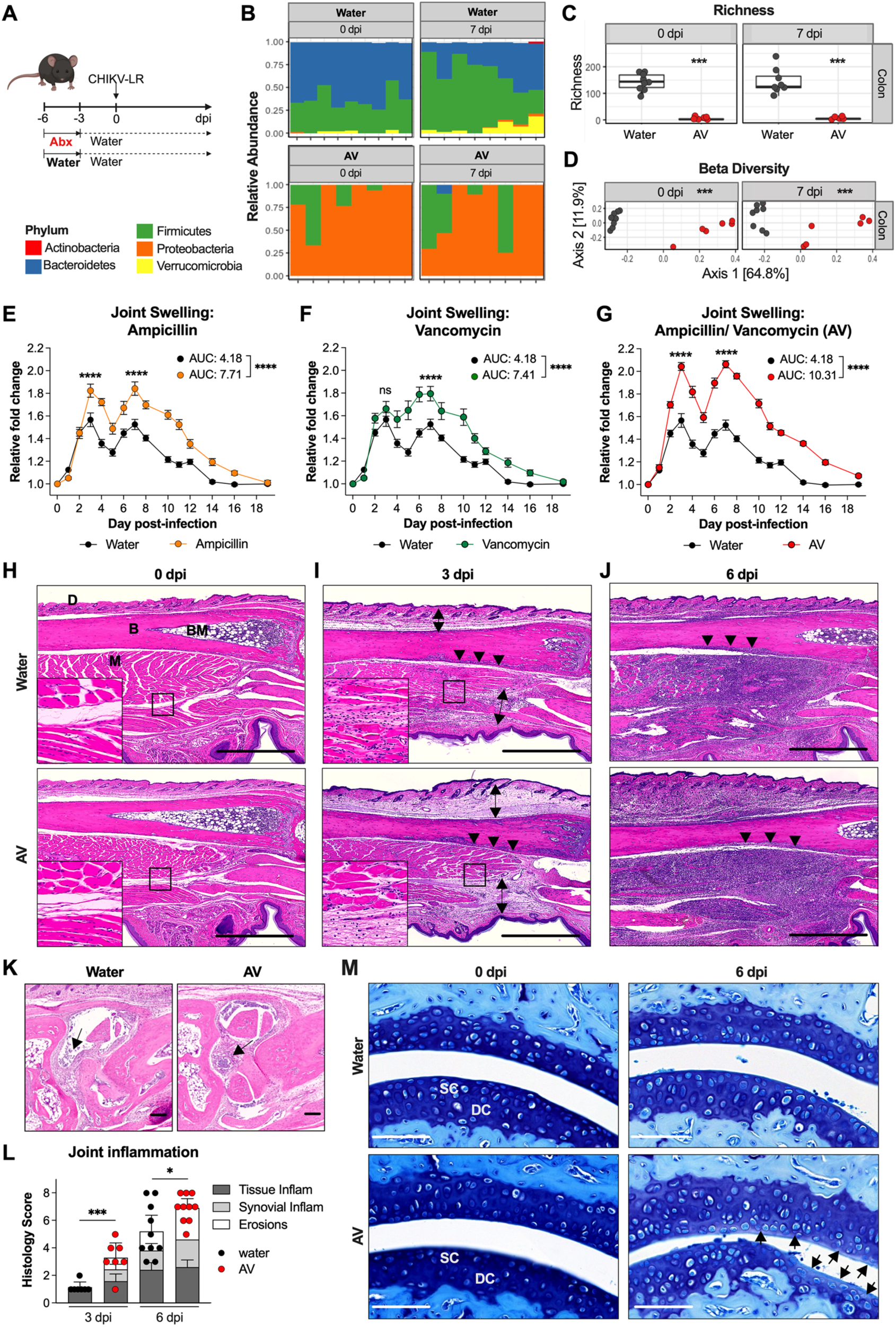
Depletion of the intestinal microbiota by oral antibiotics exacerbates musculoskeletal tissue inflammation after CHIKV infection. (A) Schematic of experimental set-up. (B-D) Colonic contents were collected from water- or AV-treated mice at 0 or 7 dpi. (B) Relative abundance of bacterial phyla detected (bacterial phyla represented in less than 1% per sample were removed), (C) number of bacterial taxa (richness), and (D) bacterial beta diversity (weighted UniFrac distance) detected in colonic contents (2 experiments, n = 8-9 per group). (E-G) Foot swelling after subcutaneous footpad injection of CHIKV in C57BL/6J mice treated with ampicillin (E, n = 10), vancomycin (F, n = 10), ampicillin/vancomycin (AV) (G, n = 15), or water (E-G, n = 15). (H-J) Hematoxylin and eosin staining of fixed foot tissues from water or AV-treated CHIKV-infected mice at (H) 0 dpi, (I) 3 dpi, and (J) 6 dpi (n = 7-10 per group). Images show 2.5x magnification; scale bars, 1 mm. D, dermis; B, bone; BM, bone marrow; M, muscle. Double-headed arrows show tissue edema. Arrow heads indicate periosteal inflammation by mononuclear cells. (K) Hematoxylin and eosin staining showing synovitis (arrows) in the joints of the mid-foot of CHIKV-infected, water or AV-treated mice at 6 dpi. Images show 5x magnification; scale bars, 100 μm. Arrows show synovial inflammation. (L) Scoring of joint inflammation and tissue damage at 3 dpi and 6 dpi. (M) Toluidine blue staining of fixed foot tissues from water or AV-treated CHIKV-infected mice at 0 and 6 dpi (representative of n = 2 for uninfected mice and n = 8 mice for infected mice, per group). Images show 40x magnification; scale bar, 100 μm. SC, superficial non-calcified cartilage; DC, deeper calcified cartilage. Arrowheads indicate areas of destaining (loss of proteoglycans) of the superficial hyaline cartilage. Statistical analysis: C, Wilcoxon tests; D, permutational multivariate analysis of variance (ADONIS); E-G, results represent the mean ± SEM from two experiments, two-way ANOVA with Šídák’s post-test or one-way ANOVA with Dunnett’s multiple comparisons test for area under the curve [AUC]; L, unpaired t-test. **** *P* < 0.0001; *** *P* < 0.001; * *P* < 0.05; ns, not significant.

Histological examination of joint tissues in the ipsilateral feet of CHIKV-infected mice at 3 days post-infection (dpi) showed greater soft tissue edema in AV-treated than in water-treated, CHIKV-infected or uninfected (0 dpi) mice (**Fig 1H-I**). Infiltration of mononuclear cells into the joint and adjacent muscle was seen in both water and AV-treated CHIKV-infected mice at this time point (**Fig 1I**). By 6 dpi, extensive soft tissue inflammation, myositis with muscle degeneration and necrosis, and periostitis was evident in tissues from all CHIKV-infected mice (**Fig 1J**). More severe synovitis and erosion of synovial membranes were apparent in the feet of CHIKV-infected, AV-treated mice (**Fig 1K-L**).

Given the association between inflammation and cartilage damage, we stained articular cartilage of water and AV-treated mice with Toluidine blue. At 6 dpi, compared to joint tissues from water-treated mice, those from AV-treated mice showed lower levels of proteoglycans in the superficial non-calcified cartilage layer (**Fig 1M**). We also performed tartrate resistant acid phosphatase (TRAP) staining to visualize osteoclasts in CHIKV-infected tissues. Normally, vascular channels connect the periosteum to the bone marrow as a means of supplying nutrients to bone marrow cells ^28^; however, these channels can enlarge or increase in number during inflammation and lead to osteoclast-induced bone erosions ^28–30^. Consistent with other etiologies of inflammatory arthritis ^30,31^, we observed transcortical vascular channels close to the epiphysis and subchondral areas of the bone in both water and AV-treated CHIKV-infected mice, with more cellularity apparent after AV-treatment (**Fig 2A**). In CHIKV-infected mice, we detected an increased density of subchondral TRAP^+^ osteoclasts in AV-treated mice at 6 dpi (**Fig 2A-C**).

**Figure 2.**
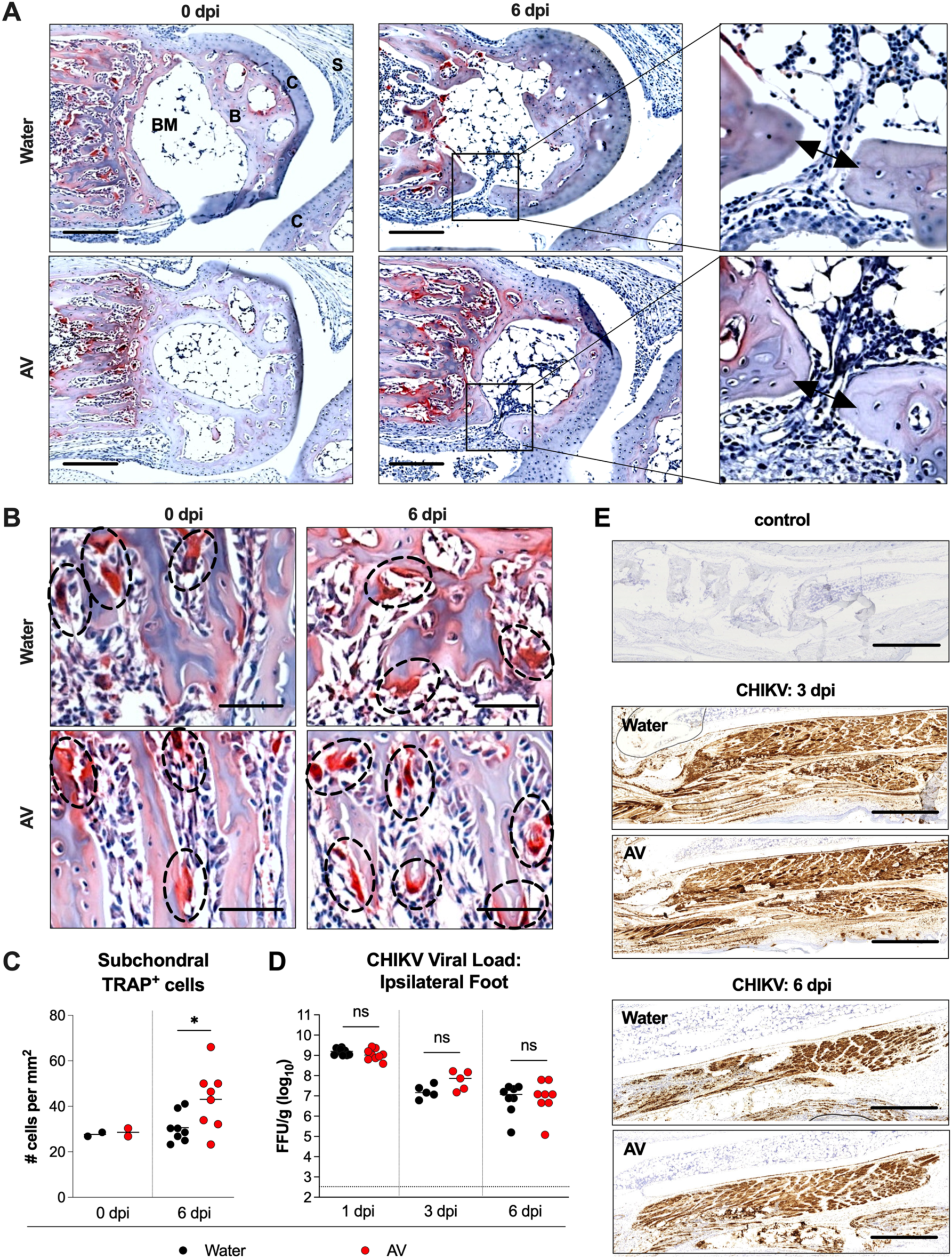
Oral antibiotics increase subchondral osteoclasts without altering viral load or tissue tropism during CHIKV infection. (**A-B**) TRAP staining of fixed foot tissues from water or AV-treated mice at 0 and 6 dpi (representative of n = 8 per group) showing red colored staining of TRAP^+^ cells. (**A**) Images showing 10x magnification; scale bar, 200 μm. B, bone; BM, bone marrow; S, synovium; C, cartilage. On higher magnification view, double-headed arrows are shown spanning the width of the vascular channels. (**B**) Dashed ovals encircle subchondral TRAP^+^ osteoclasts (red, nucleated). (**C**) Quantitation of subchondral TRAP^+^ osteoclasts in water and AV-treated mice at 0 and 6 dpi. (**D**) CHIKV loads in ipsilateral feet of water- and AV-treated mice at 1, 3, and 6 dpi were determined by focus forming assay (2 experiments, n = 5-8 per group). (**E**) CHIKV RNA *in situ* hybridization of foot tissue sections from Zika virus (ZIKV)-infected mice (negative control, top panel), and CHIKV-infected, water- or AV-treated mice at 3 dpi (middle panels) or 6 dpi (bottom panels) (representative of n = 3 per group). Images show 2.5x magnification; scale bars, 1 mm. Statistical analysis: **C-D**, Mann-Whitney test. * *P* < 0.05; ns, not significant.

To determine whether antibiotic-mediated exacerbation of CHIKV arthritis was related to increased viral burden or altered cell tropism within the affected foot, we measured levels of virus infection and visualized viral RNA within joint tissues using *in situ* hybridization. Notably, viral titers in the ipsilateral foot at 3 and 6 dpi were equivalent, and intense viral RNA staining was detected in the periosteum, articular cartilage, synovium, and muscles in both water and AV-treated infected mice (**Fig 2D-E**). These data suggest that the increased musculoskeletal tissue swelling in the affected foot of AV-treated animals is not due to greater viral infection.

### Oral antibiotic treatment increases immune cell infiltration and cytokine levels in musculoskeletal tissues after CHIKV infection

We next evaluated the effects of AV treatment on inflammation in CHIKV-infected joint-associated tissues. At 4 dpi, AV-treated mice had higher levels of IL-4, IL-5, CCL11, LIF, IL-6, IFN-γ, TNF, M-CSF, and G-CSF, as well as the IFN-induced chemokines, CCL2, CCL3, CCL4, CCL5, CXCL1, CXCL9 and CXCL10, in the ipsilateral foot (**Fig 3A and S1I-J**) compared to water-treated, infected mice. To characterize the effect of microbiota perturbation on joint-associated immune cell infiltrates, we analyzed by flow cytometry the composition of cells in the ipsilateral foot after CHIKV infection (**Fig S2A**). AV-treated mice had increased numbers of monocytes and neutrophils in the foot at 4 dpi and increased neutrophils at 7 dpi (**Fig 3B-C**). Statistically significant differences in numbers of NK, T, or B cells in joint-associated tissues were not observed (**Fig 3B-C**).

**Figure 3.**
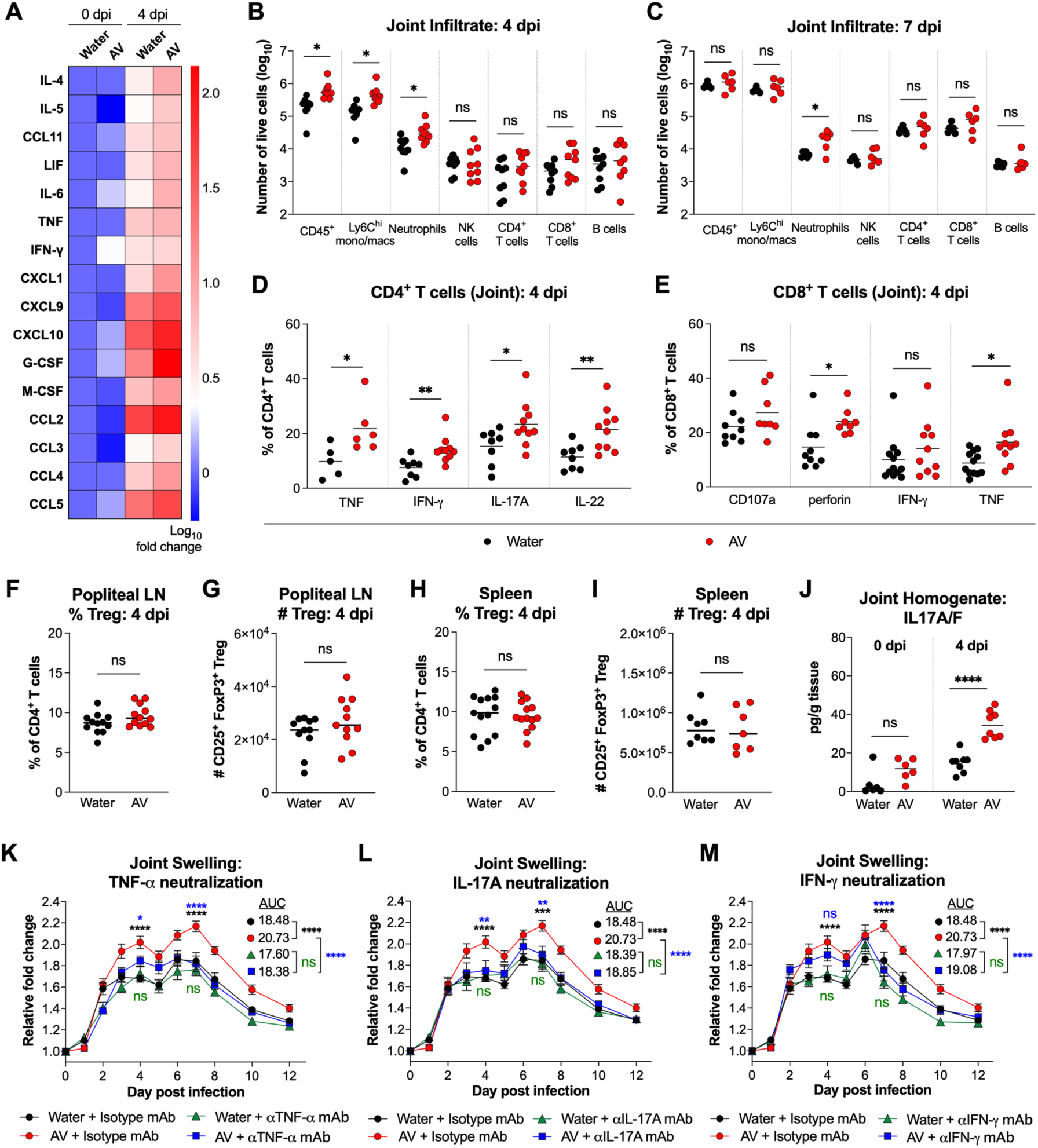
Oral antibiotics increase joint inflammation during CHIKV infection. (**A**) Heat map of cytokine and chemokine levels in foot-associated musculoskeletal tissue homogenates from water- or AV-treated mice at 0 and 4 dpi (2 experiments, n = 5-8 per group, log_10_-fold differences in protein levels are relative to water-treated animals at 0 dpi, see also **Fig S1J**). (**B-C**) Numbers of infiltrating immune cells in joint-associated tissues in the ipsilateral feet of water- or AV-treated mice were determined by flow cytometry on (**B**) 4 dpi or (**C**) 7 dpi (2 experiments, n = 6-9 per group). (**D**) Percentages of IFN-γ-, TNF-, IL-17-, and IL-22-producing CD4^+^ T cells in the ipsilateral feet of water- and AV-treated mice at 4 dpi (2-3 experiments, n = 6-11 per group). (**E**) Percentages of IFN-γ- and TNF-producing, perforin-expressing, or CD107a-expressing CD8^+^ T cells in the ipsilateral feet of control and AV-treated mice at 4 dpi (3 experiments, n = 8-17 per group). (**F-I**) Percentages and numbers CD4^+^ CD25^+^ FoxP3^+^ Tregs in the (**F-G**) draining popliteal lymph nodes and (**H-I**) spleens of water- and AV-treated mice at 4 dpi. (**J**) IL-17A/F levels in joint-associated tissue homogenates from ipsilateral feet as detected by ELISA (2 experiments, n = 5-8 per group). (**K-M**) Foot swelling of water- and AV-treated mice that were administered isotype control mAb or neutralizing mAb against (**K**) TNF, (**L**) IL-17A, or (**M**) IFN-γ (2 experiments, n = 8-9 per group). Statistical analysis: **B-J**, unpaired t-test; **K-M**, two-way ANOVA with Šídák’s post-test, or one-way ANOVA with Dunnett’s multiple comparisons test for AUC; mean values ± SEM are shown. **** *P* < 0.0001; *** *P* < 0.001; ** *P* < 0.01; * *P* < 0.05; ns, not significant.

We also assessed whether there was a qualitative difference of the T cells in joint-associated tissues in water- and AV-treated mice after CHIKV infection. AV-treated mice had greater proportions and numbers of inflammatory TNF-, IFN-γ-, IL-17A-, and IL-22-producing CD4^+^ T cells in the joint at 4 dpi (**Fig 3D and Fig S2B**). However, similar percentages of degranulation and IFN-γ production were observed in CD8^+^ T cells from the joints of water- and AV-treated mice at 4 dpi, although there was a small increase in the proportion of perforin- and TNF-expressing CD8^+^ T cells in AV-treated mice (**Fig 3E**). Significant differences in the percentages or numbers of FoxP3^+^ regulatory T cells (Tregs) were not detected between water-and AV-treated mice in the draining popliteal lymph node (**Fig 3F-G**) or spleen (**Fig 3H-I**) at 4 dpi. Using enzyme-linked immunosorbent assays, we confirmed that AV-treated mice had elevated levels of IL-17A, TNF and IFN-γ in joint-associated tissues at 4 dpi compared to water-treated mice (**Fig 3J and S1I**). To determine whether these cytokines contributed to the enhanced tissue inflammation during CHIKV infection associated with microbiota perturbation, we administered neutralizing antibodies to water- and AV-treated mice. Treatment with anti-TNF or anti-IL-17A monoclonal antibodies (mAb) improved foot-swelling in antibiotic-treated mice following CHIKV infection (**Fig 3K-L**). Neutralization of IFN-γ in AV-treated mice led to comparatively smaller reductions in inflammation at 3-4 dpi but substantially less tissue swelling at and after 7 dpi (**Fig 3M**). Thus, antibiotic-mediated dysbiosis promotes polarization of CD4^+^ T cells and infiltration of myeloid cells into the joint, resulting in higher levels of pro-inflammatory cytokines that augment foot swelling after CHIKV infection.

### Anaerobic commensal bacteria regulate musculoskeletal inflammation after CHIKV infection

We next evaluated whether we could rescue the exacerbated joint swelling phenotype after AV treatment by reconstituting the microbiota with selected bacterial species. Colonization of AV-treated mice with fecal microbiota from untreated mice normalizes the intestinal microbiota to that of untreated, conventionally housed mice ^32^. Moreover, fecal microbial transfer (FMT) from untreated mice reversed the exacerbated CHIKV-induced joint swelling phenotype seen in AV-treated mice (**Fig 4A and S2C**). To determine whether specific microbial communities modulated the severity of musculoskeletal inflammation after CHIKV infection, we colonized AV-treated mice with anaerobic or aerobic cultures derived from FMT. Reconstitution with anaerobic, but not aerobic, cultures reversed CHIKV-induced tissue swelling and immune cell infiltration to levels seen in untreated, infected mice (**Fig 4B-C**).

**Figure 4.**
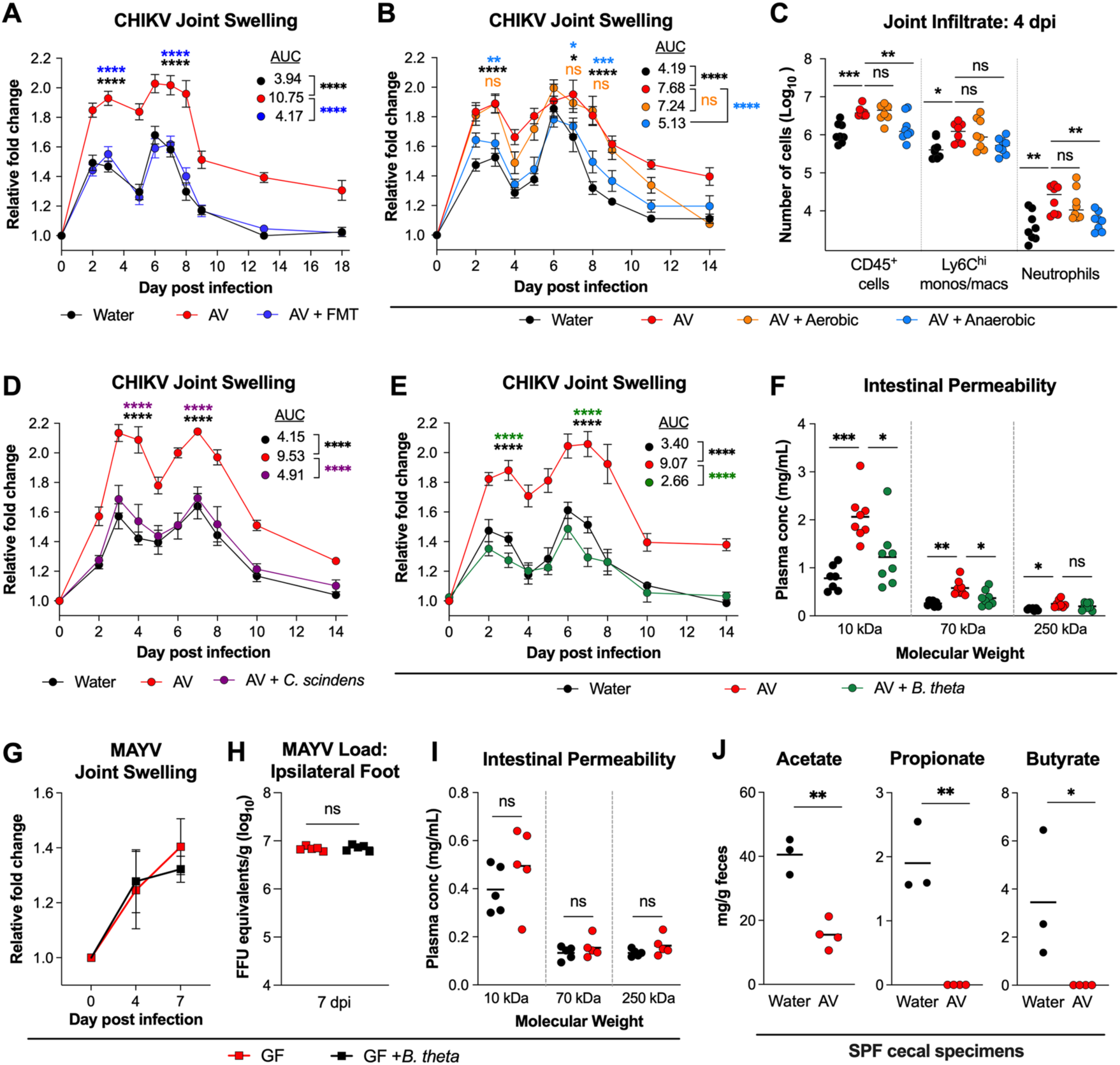
Microbial dysbiosis results in reduced SCFA production, increased intestinal permeability and enhanced joint swelling. (**A**) Foot swelling after CHIKV infection in water-treated, AV-treated, or AV-treated mice after fecal microbial transfer [FMT] (3 experiments, n = 13-14 per group). (**B-C**) Foot swelling (**B**) and immune cell infiltrates in the feet (**C**) after CHIKV infection of water-treated, AV-treated, or AV-treated mice colonized with either aerobic or anaerobic fecal cultures of FMT (2 experiments, n = 8-9 per group). (**D-E**) Foot swelling after CHIKV infection of water-treated, AV-treated, or AV-treated mice colonized with either (**D**) *C. scindens* or (**E**) *B. thetaiotaomicron* (2 experiments, n = 9 per group). (**F**) Plasma concentration of 10 kDa, 70 kDa, and 250 kDa dextrans at 1.5 h after oral gavage as a measure of paracellular intestinal permeability in mice treated as indicated (2 experiments, n = 8). (**G-I**) GF mice received PBS or colonization with *B. thetaiotaomicron* via oral gavage prior to MAYV infection. (**G**) Foot swelling after MAYV infection and (**H**) MAYV burden in tissue homogenates from the feet of GF mice or GF mice colonized with *B. thetaiotaomicron*. (**I**) Plasma concentration of 10 kDa, 70 kDa, and 250 kDa dextrans at 1.5 h after oral gavage in mice treated as indicated. (**J**) Fecal SCFA levels in water- or AV-treated mice. Statistical analysis: **A-B, D-E,** two-way ANOVA with Dunnett’s post-tests; AUC analyses were performed with one-way ANOVA with Šídák’s post-test; mean values ± SEM are shown. **C, F**, one-way ANOVA with Šídák’s post-test. **H**, Mann-Whitney test. **I-J**, unpaired t-test. **** *P* < 0.0001; *** *P* < 0.001; ** *P* < 0.01; * *P* < 0.05; ns, not significant.

We next tested whether individual anaerobic bacterial species could modulate musculoskeletal inflammation after CHIKV infection. We focused on *Bacteroides* and *Clostridium* species since these commensals are depleted by our oral AV regimen (**Fig 1B and S1A**) and have known effects on host immunity ^33–35^. Remarkably, colonization of AV-treated mice with either *Clostridium scindens* or *Bacteroides thetaiotaomicron* reversed the CHIKV-induced musculoskeletal inflammation to levels comparable to water-treated animals (**Fig 4D-E**). These data demonstrate a functional redundancy in the ability of different commensal gut bacteria to regulate virus-induced inflammation at the distal joint tissue site.

### Antibiotic treatment results in increased paracellular intestinal permeability

The gut microbiota interfaces with the intestinal epithelium to modulate permeability through transcellular (transcytosis or carrier-dependent transport) or paracellular (tight-junction-dependent) routes. Gut microbial dysbiosis can be associated with translocation of microbes, microbial products, or metabolites into the lamina propria and systemic circulation ^36,37^. To evaluate the effect of AV-treatment on intestinal permeability *in vivo*, we measured translocation of orally-gavaged fluorescently conjugated dextrans of different molecular weight from the intestinal lumen into the peripheral blood. We observed increased serum levels of 10 kilodalton (kDa), 70 kDa, and 250 kDa dextrans in AV-treated mice compared to water-treated animals at 1.5 h after oral gavage (**Fig 4F**), although the larger 250 kDa dextrans were minimally translocated in both groups. Colonization of AV-treated mice with *B. thetaiotaomicron* restored the intestinal barrier integrity (**Fig 4F**), suggesting a possible linkage between the gut microbiota, intestinal permeability, and CHIKV-induced tissue inflammation.

We next investigated the requisite role of intestinal microbes in exacerbating alphavirus-induced musculoskeletal inflammation. GF mice that were monocolonized with *B. thetaiotaomicron* or maintained as GF were inoculated with the related alphavirus, MAYV, and evaluated for foot swelling; MAYV was used, instead of CHIKV, because our gnotobiotic facility operates under A-BSL2 conditions (CHIKV require A-BSL3 containment). Non-colonized and *B. thetaiotaomicron*-colonized GF mice exhibited similar levels of foot swelling at 4 and 7 days after MAYV infection (**Fig 4G**), with similar levels of viral RNA in the foot (**Fig 4H**). Dextran translocation via paracellular intestinal permeability was also similar between GF mice and those monocolonized with *B. thetaiotaomicron* (**Fig 4I**). Together, these data suggest that perturbations of specific gut microbiota components, rather than depletion or an absence of microbiota, modulate intestinal permeability and joint inflammation after alphavirus infection.

### SCFA regulate intestinal permeability and joint inflammation after CHIKV infection

The intestinal microbiota can regulate immune responses through direct effects on immune cells or via indirect effects on the gut epithelial barrier ^36–38^. We hypothesized that the exacerbated CHIKV-induced foot swelling phenotype might be due to the loss of specific commensal bacteria and their metabolites that modulate the GI tract epithelial barrier. *Bacteroidetes*, the major phylum of bacteria depleted by our antibiotic regimen (**Fig 1B and S1A**), as well as *Clostridia*, are primary producers of SCFA and biotransformers of secondary bile acids (BAs) in the intestinal microbiota ^39,40^. Indeed, we previously found that levels of deoxycholic acid (DCA), a secondary BA, are reduced in antibiotic-treated mice ^32^. Although oral supplementation with DCA improves early systemic type I IFN responses and decreases viremia ^32^, the intestinal permeability and foot swelling after CHIKV infection worsened (**Fig S2D-F**), consistent with reports showing that secondary BAs disrupt intestinal barrier functions *in vitro* and *in vivo* ^41–44^. These results demonstrate that some microbial metabolites can have distinct roles in modulating antiviral immunity and tissue inflammation.

The three most abundantly detected SCFA (acetate, propionate, and butyrate) also were substantially reduced after AV treatment (**Fig 4J, S2G, and S3A**). In the intestines of mice and humans, these SCFA promote intestinal epithelial cell homeostasis and barrier function ^45^, and induction of regulatory immune cells ^46–48^. Because of these functions, we evaluated whether oral supplementation with individual SCFA could restore intestinal barrier integrity and modulate CHIKV-induced foot swelling in AV-treated mice (**Fig S2D**). Orally delivered butyrate or propionate attenuated CHIKV-induced foot swelling of AV-treated mice to levels comparable to those of water-treated mice (**Fig 5A-C**). Butyrate or propionate treatment also reduced the numbers of infiltrating Ly6C^hi^ monocytes and neutrophils, decreased the proportions of CD4^+^ T cells producing IFN-γ, IL-17, and IL-22, and restored intestinal barrier integrity (**Fig 5D-F**). In comparison, orally administered acetate did not change intestinal permeability, reduce myeloid cell infiltration, diminish pro-inflammatory cytokine production by CD4^+^ T cells, or protect against CHIKV-induced joint swelling after AV treatment (**Fig 5C-F**). The protective effect of butyrate or propionate supplementation was not due to altered viral infection within the foot (**Fig 5G**) and was specific to mice with perturbed microbiota due to antibiotic treatment; SCFA supplementation in conventionally housed, non-antibiotic treated mice did not reduce or exacerbate foot swelling caused by CHIKV infection (**Fig 5A-C**), in contrast to results from a prior study ^49^.

**Figure 5.**
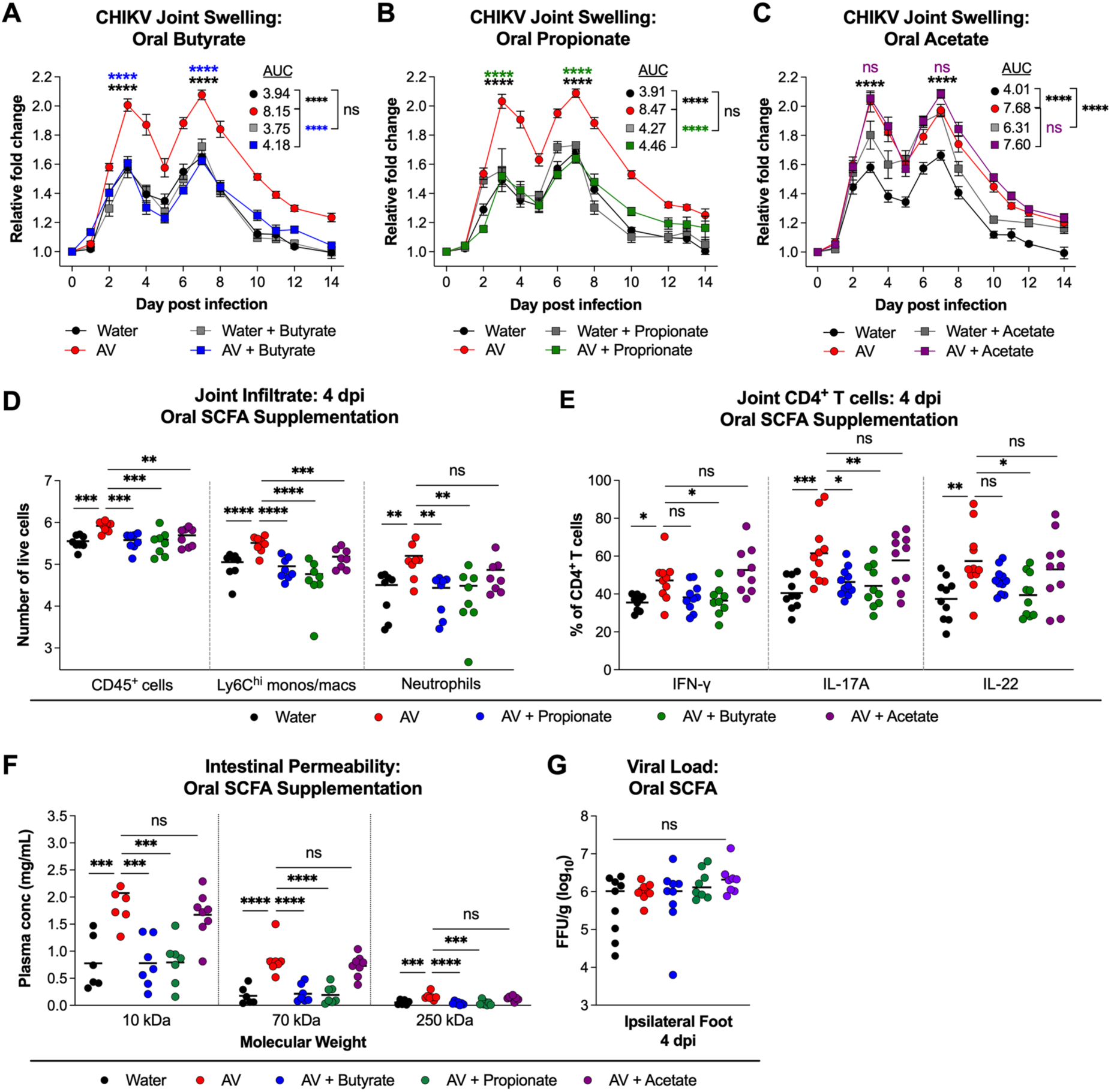
Oral supplementation with exogenous SCFA rescues antibiotic-mediated exacerbation of CHIKV-induced foot swelling. Mice were treated with water or AV, with or without SCFA supplementation (butyrate, propionate, or acetate), and subsequently inoculated with CHIKV. (**A-C**) Foot swelling after CHIKV infection in water- or AV-treated mice with or without supplementation with (**A**) butyrate, (**B**) propionate, or (**C**) acetate (2-3 experiments, n = 10-12 per group). (**D**) Numbers of joint-associated immune cells at 4 dpi (2 experiments, n = 8 per group). (E) Percentages of cytokine-producing CD4^+^ T cells at 4 dpi (3 experiments, n = 8-11 per group). (F) Intestinal permeability as measured by plasma concentrations of 10 kDa, 70 kDa, and 250 kDa dextran at 1.5 h after oral gavage (2 experiments, n = 6-7 per group). (**G**) CHIKV viral load at 4 dpi in tissue homogenates from the ipsilateral feet of mice treated as indicated (2 experiments, n = 8-9 per group). Statistical analysis: **A-C**, two-way ANOVA with Dunnett’s post-tests; AUC analyses were performed with one-way ANOVA with Šídák’s post-test; mean values ± SEM are shown. **D-G,** one-way ANOVA with Šídák’s post-test. **** *P* < 0.0001; *** *P* < 0.001; ** *P* < 0.01; * *P* < 0.05; ns, not significant.

### Rescue of enhanced joint inflammation by bacteria requires microbe-derived SCFA

To link our findings on SCFA and bacterial reconstitution, we assessed whether the capacity of *B. thetaiotaomicron* to restore the intestinal epithelial barrier and mitigate the inflammatory response to CHIKV infection in joint-associated tissues depended on its ability to produce SCFA. We used the double-crossover gene deletion method ^50,51^ to generate an isogenic clone of *B. thetaiotaomicron* that lacks the ability to generate propionate, by targeting methylmalonyl-CoA mutase (*τιBT2090-2091*). The loss of propionate production in *τιBT2090-2091 B. thetaiotaomicron* (*B. theta τιP*) was validated by gas chromatography mass spectrometry on extracts from *in vitro* cultures (**Fig 6A**).

**Figure 6.**
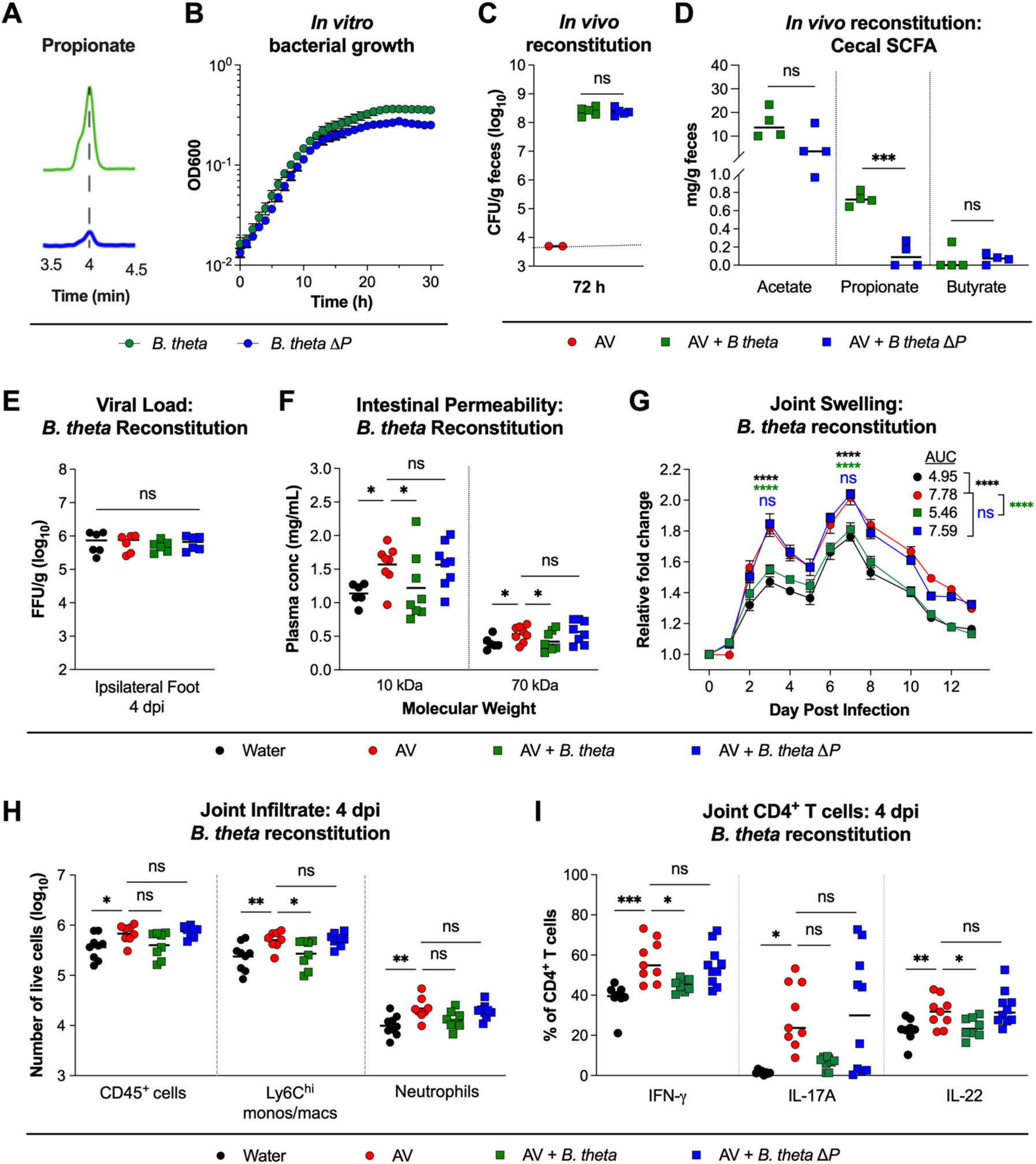
Rescue of CHIKV-induced foot swelling by bacterial colonization depends on microbe-derived SCFA. (**A**) *In vitro* production of propionate by isogenic wild-type and mutant *(ΔP) B. thetaiotaomicron*. (**B**) *In vitro* growth properties of isogenic wild-type and mutant *B. thetaiotaomicron*. (**C-D**) AV-treated mice were gavaged with sterile PBS or with wild-type or mutant *B. thetaiotaomicron*. (**C**) Fecal colony-forming unit counts on Bacteroides Bile Esculin agar. (**D**) Cecal SCFA levels were determined by gas chromatography mass spectrometry. (**E-I**) Water-treated, AV-treated, or AV-treated mice colonized with either wild-type or mutant *B. thetaiotaomicr*o*n* were inoculated with CHIKV. (**E**) CHIKV viral load at 4 dpi in homogenates of the ipsilateral feet (2 experiments, n = 6 per group). (**F**) Plasma concentration of 10 kDa and 70 kDa dextran was measured at 1.5 h after gavage of mice treated as indicated (3 experiments, n = 10-13 per group). (**G**) Foot swelling was measured after CHIKV infection (2 experiments, n = 10 per group). (**H**) Numbers of joint-associated immune cells and (**I**) percentages of cytokine-producing CD4^+^ T cells at 4 dpi (2 experiments, n = 8-10 per group). Statistical analysis: **C**, **E**, Mann-Whitney test. **D**, unpaired t-test. **F**, **H-I**, one-way ANOVA with Šídák’s post-test. **G**, two-way ANOVA with Tukey’s post-test; AUC was analyzed using one-way ANOVA with Dunnett’s post-test; mean values ± SEM are shown. **** *P* < 0.0001; *** *P* < 0.001; ** *P* < 0.01; * *P* < 0.05; ns, not significant.

Wild-type and *τιBT2090-2091 B. thetaiotaomicron* showed similar growth kinetics *in vitro* and after colonization in mice that had been pulsed with AV (**Fig 6B-C**). We confirmed that cecal samples from mice colonized with *τιBT2090-2091 B. thetaiotaomicron* had less propionate than animals colonized with wild-type *B. thetaiotaomicron* (**Fig 6D, S2G, and S3A**). In contrast to the parental strain, colonization with the *τιBT2090-2091 B. thetaiotaomicron* failed to restore intestinal permeability, diminish monocyte infiltrates or inflammatory cytokine production by CD4^+^ T cells in the joint, or reduce foot swelling at 4 days after CHIKV infection (**Fig 6E-I**).

### Enhanced joint inflammation after oral antibiotic treatment requires MyD88 signaling

We hypothesized that antibiotic-mediated perturbation of the intestinal microbiota and associated gut permeability promotes joint inflammation via sensing of microbe-derived signals by host pattern recognition receptors. MyD88 is a key adaptor protein linking the TLR and interleukin-1 (IL-1) receptor families to downstream inflammatory pathways ^52^. To assess whether MyD88 signaling contributes to the exacerbated inflammation phenotype, we treated *Myd88*^-/-^ and wild-type littermate mice with water or AV and evaluated immune cell infiltration and foot swelling after CHIKV infection. A loss of MyD88 signaling reversed the exacerbated CHIKV-induced foot swelling in AV - treated mice and reduced immune cell infiltration in the foot at 4 dpi (**Fig 7A-B**). Additionally, IFN-γ, IL-17, and IL-22 production by infiltrating CD4^+^ T cells also were reduced in AV-treated *Myd88*^-/-^ mice (**Fig 7C**). In contrast, viral titers were slightly but not significantly increased in the ipsilateral feet of *Myd88^-/-^*mice compared to wild-type littermate controls (**Fig 7D**).

**Figure 7.**
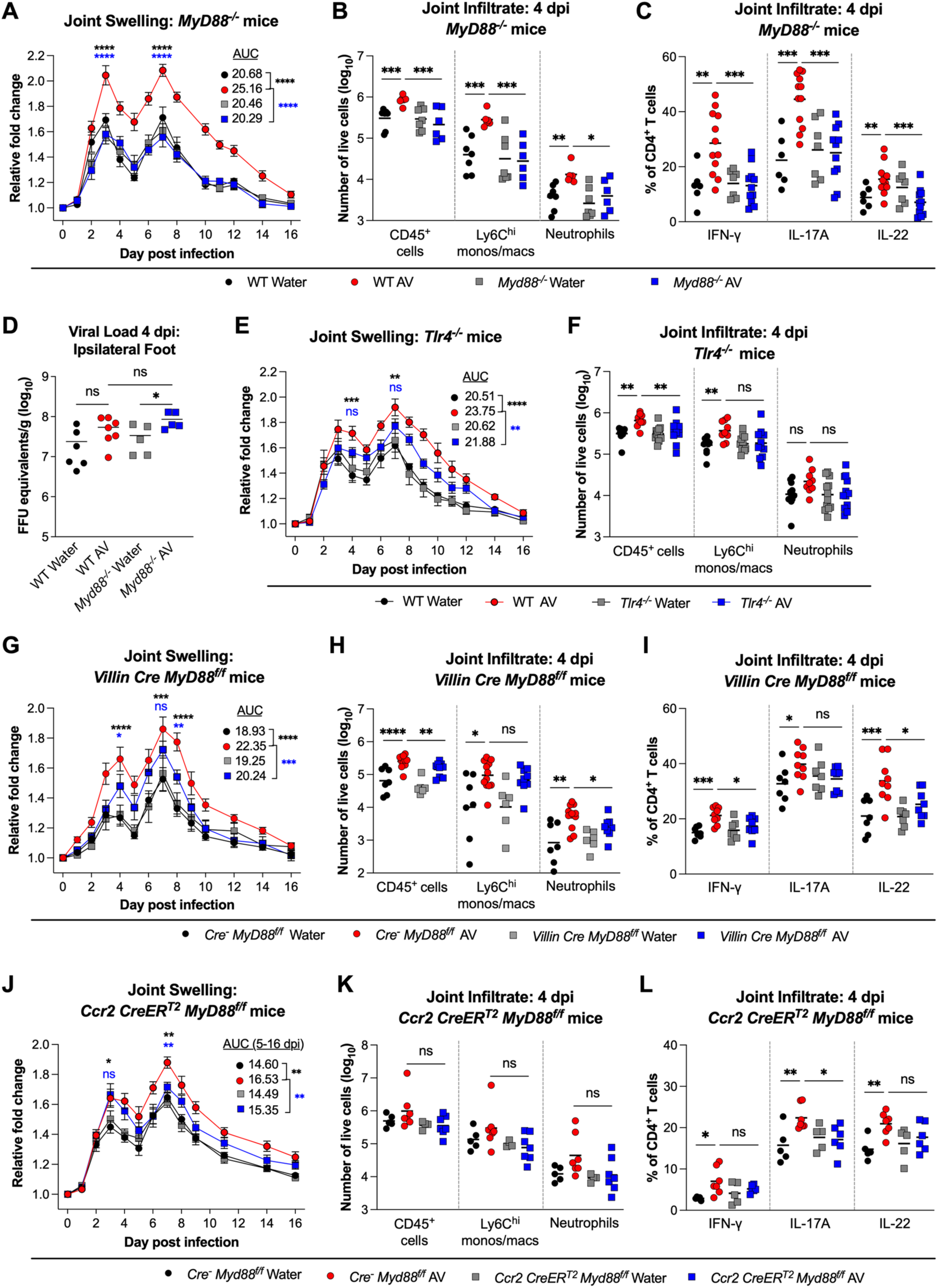
MyD88 signaling in intestinal epithelial cells promotes joint inflammation after CHIKV infection in the setting of dysbiosis. (**A-C**) Water- or AV-treated *Myd88^-/-^* and WT littermate control mice were evaluated for (**A**) foot swelling after CHIKV infection, (**B**) immune cell accumulation in the joint at 4 dpi, and (**C**) percentages of IFN-□-, IL-17-, and IL-22-producing CD4^+^ T cells in the joint at 4 dpi (2 experiments, n = 6-12 per group). (**D**) CHIKV RNA levels at 4 dpi in tissue homogenates from the feet of water or AV-treated *Myd88^-/-^* and WT littermate mice (2 experiments, n = 5-7 per group). (**E-F**) Water- or AV-treated *Tlr4^-/-^* and WT littermate control mice were assessed for (**E**) foot swelling after CHIKV infection and (**F**) immune cell accumulation at 4 dpi (2 experiments, n = 9-12 per group). (**G-I**) Water- or AV-treated *Villin*-Cre *Myd88^f/f^* and littermate control Cre^-^ *Myd88^f/f^* mice were inoculated with CHIKV. (**G**) Foot swelling after infection, (**H**) immune cell accumulation in the joint at 4 dpi, and (**I**) percentages of IFN-□-, IL-17-, and IL-22-producing CD4^+^ T cells in the joint at 4 dpi (2 to 3 experiments, n = 7-9 per group). (**J-L**) Water or AV-treated *Ccr2*-CreER^T2^ *Myd88^f/f^* mice and littermate control Cre^-^ *Myd88^f/f^* mice were pretreated with tamoxifen to induce Cre recombinase activity prior to CHIKV inoculation. (**J**) Foot swelling after infection, (**K**) immune cell accumulation in the joint at 4 dpi, and (**L**) percentages of IFN-□-, IL-17-, and IL-22-producing CD4^+^ T cells in the joint at 4 dpi (2 experiments, n = 5-7 per group). Statistical analysis: **A, E, G,** and **J**, two-way ANOVA with Tukey’s post-test, or one-way ANOVA with Dunnett’s post-test for AUC; mean values ± SEM are shown. **B-D, F, H-I, K-L,** one-way ANOVA with Šídák’s multiple comparisons test. **** *P* < 0.0001, *** *P* < 0.001, ** *P* < 0.01, * *P* < 0.05, ns, not significant.

We next investigated the potential receptors upstream of MyD88 signaling that trigger the inflammation in AV-treated CHIKV-infected animals. IL-1α and IL-1β are pro-inflammatory cytokines that signal via the IL-1 receptor, which is expressed on a range of cells including epithelial and immune cells. However, IL-1 signaling did not appear to contribute to the inflammatory phenotype, as AV-treatment of *Il1r*^-/-^ and wild-type littermate mice showed similarly worsened foot swelling after CHIKV infection (**Fig S3B**). We also considered the role of TLRs that sense pathogen-associated molecular patterns (PAMPs), including bacterial cell wall and membrane components as well as nucleic acids. In a series of studies with AV-treated congenic TLR-deficient or wild-type littermate mice, we determined that the enhanced CHIKV-induced swelling and Ly6C^hi^ monocyte infiltration in the foot were reduced at least partially in AV-treated *Tlr4^-/-^* mice (**Fig 7E-F**) but not in *Tlr2^-/-^*, *Tlr7^-/-^*, or *Tlr9^-/-^* mice (**Fig S3C-E**). These data suggest that perturbations in the intestinal microbiota result in TLR4- and MyD88-dependent signals that promote musculoskeletal inflammation in the context of alphavirus infection.

Although MyD88 signaling in intestinal epithelial cells is critical for maintaining gut homeostasis and protecting against intestinal bacterial infections ^53–57^, it also can contribute to extra-intestinal inflammation ^58,59^. We hypothesized that MyD88 signaling in enterocytes triggered by gut dysbiosis might contribute to the aggravated joint swelling in AV-treated mice after CHIKV infection. We generated *Villin*-Cre *Myd88*^flox/flox^ ^(f/f)^ mice to evaluate the role of intestinal epithelial cell MyD88 signaling in the joint inflammation associated with AV treatment and CHIKV infection. Compared to Cre-negative littermate controls, loss of intestinal epithelial cell expression of MyD88 partially decreased the AV-induced swelling phenotype after CHIKV infection (**Fig 7G**). Although reduced numbers of infiltrating monocytes were not observed in the ipsilateral feet of AV-treated *Villin*-Cre *Myd88*^f/f^ mice at 4 dpi, there was a small reduction in overall CD45^+^ immune cells and neutrophils (**Fig 7H**), and the production of IFN-γ and IL-22 by CD4^+^ T cells was lower (**Fig 7I**). These data suggest that in the context of dysbiosis, MyD88-dependent signaling in intestinal epithelial cells contributes to distal tissue inflammation by modulating immune cell recruitment and CD4^+^ T cell function. Although intestinal tuft cells can sense signals from the microbiota and link them to immune responses ^60,61^, improved foot swelling after CHIKV infection was not observed in *Pou2f3*^-/-^ mice, which lack tuft cells (**Fig S3F**).

Given the substantial joint infiltration by monocytes, we used *Ccr2*-Cre-ER^T2^ *Myd88*^f/f^ mice to evaluate whether MyD88 signaling in monocytes also contributed to joint inflammation. Following AV treatment, when compared to Cre-negative littermate control mice, *Ccr2*-Cre-ER^T2^ *Myd88*^f/f^ mice showed minimal differences in swelling at 3 to 4 dpi, decreased IL17 production by CD4^+^ T cells, and slightly reduced but non-significant differences in infiltrating immune cells in joint-associated tissues (**Fig 7J-L**). However, reduced foot swelling was observed in AV-treated *Ccr2*-Cre-ER^T2^ *Myd88*^f/f^ mice at 7 dpi during the second phase (**Fig 7J**). Thus, although monocyte accumulation in the joint correlates with enhanced inflammation in the setting of AV treatment, the effect is only partially mediated through cell-intrinsic MyD88-dependent activation.

### Monocytes and CD4^+^ T cells are required for exacerbated CHIKV-induced joint swelling in antibiotic-treated mice

We next evaluated the requirement for specific immune cell populations in mediating the increased CHIKV-induced foot swelling after AV treatment. To determine whether myeloid cells contribute, we initially used anti-Ly6C/Ly6G (Gr-1) mAb to deplete neutrophils, eosinophils, and monocytes prior to AV treatment and CHIKV infection (**Fig S3G-H**). Upon depletion of these cells, foot swelling caused by CHIKV infection was attenuated in AV-treated mice to levels like those of isotype control mAb, water-treated animals (**Fig 8A**). Whereas depletion of neutrophils alone with a mAb against Ly6G (1A8) did not affect CHIKV-induced foot swelling after AV treatment (**Fig 8B, S3G, and S3I**), inhibition of monocyte trafficking using an anti-CCR2 mAb normalized the first swelling peak (**Fig 8C, S3G, and S3J**). The lack of improvement during the second swelling peak (at 7 dpi) might be due to a compensatory influx of neutrophils and eosinophils that occurs during the later phase of CHIKV infection when monocytes and macrophages are depleted ^62^. In aggregate, these data suggest that circulating monocytes contribute to enhanced CHIKV-induced foot swelling in mice after AV treatment.

**Figure 8.**
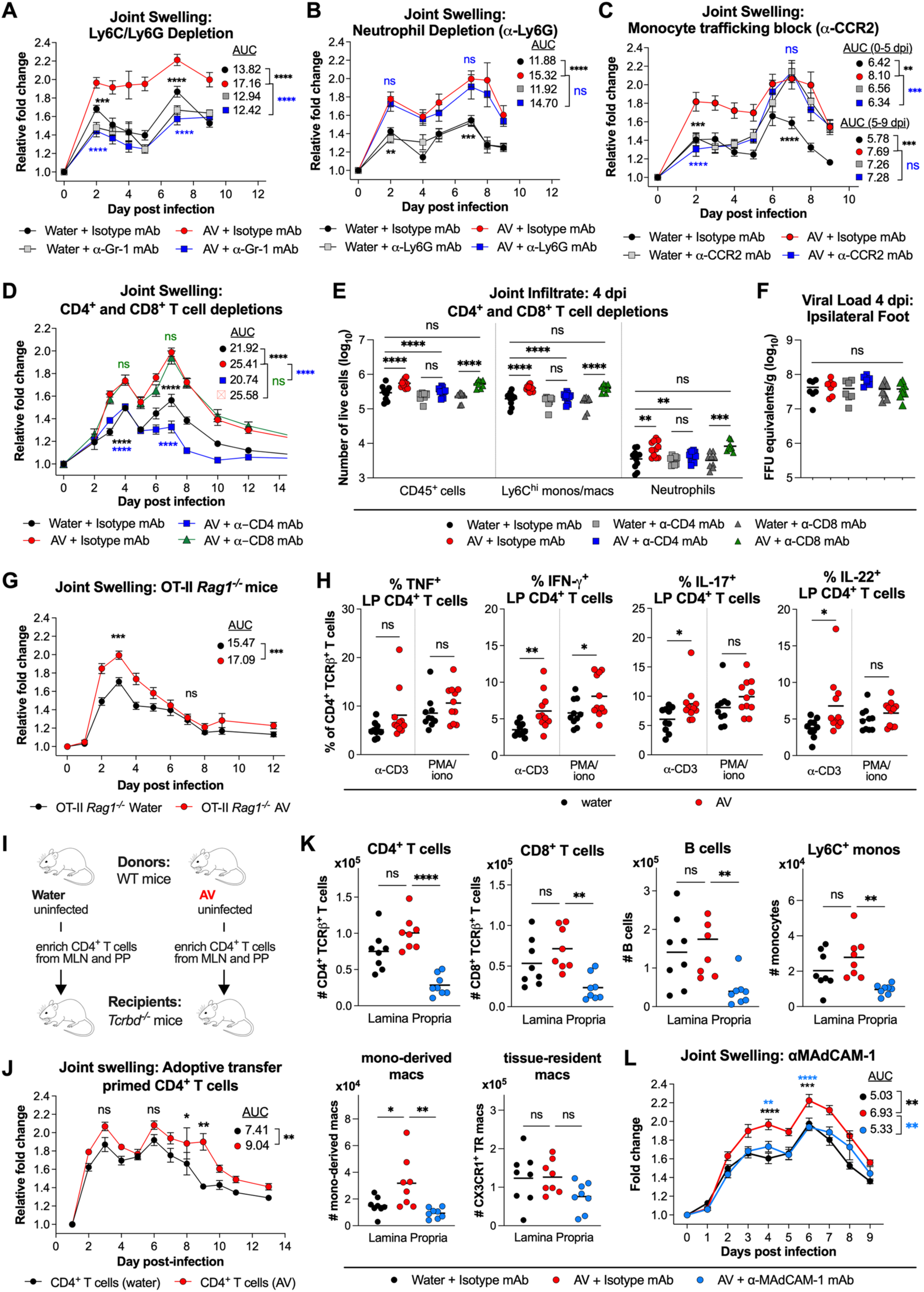
Monocytes and CD4^+^ T cells contribute to increased tissue inflammation associated with gut microbiome depletion. (A-C) Foot swelling after CHIKV infection of water or AV-treated mice that were administered isotype control mAb or mAbs against. (**A**) Gr-1 (Ly6C/Ly6G) (2 experiments, n = 9-10 per group), (**B**) Ly6G (2 experiments, n = 8 per group), or (**C**) CCR2 (2 experiments, n = 8-9 per group). (**D-F**) Mice that received either isotype control or depleting mAbs against CD4^+^ or CD8^+^ T cells were treated with water or AV, and subsequently infected with CHIKV. (**D**) Foot swelling (2 experiments, n = 9 per group), (**E**) immune cells in the joint (3 experiments, n = 9-12 per group), and (**F**) viral load in the ipsilateral foot at 4 dpi (2 experiments, n = 7-8 per group). (**G**) Foot swelling after CHIKV infection of water or AV-treated OT-II *Rag1^-/-^* mice (2 experiments, n = 6-7 per group). (**H**) Percent of CD4^+^ T cells in the colonic lamina propria producing each of the indicated cytokines in response to either a-CD3 or PMA/ionomycin stimulation. (**I-J**) CD4^+^ T cells were enriched from the MLNs and Peyer’s patches of water or AV-treated mice and adoptively transferred into *Tcrbd^-/-^*mice (**I**) at one day prior to inoculation with CHIKV. (**J**) Foot swelling after CHIKV infection of *Tcrbd^-/-^* mice that received CD4^+^ T cells from either water- or AV-treated mice. (**K-L**) Mice received isotype control or mAbs against MAdCAM-1 prior to treatment with water or AV and subsequent CHIKV infection (2 experiments, n = 8 per group). (**K**) Numbers of the indicated immune cells in the lamina propria of the colon. (**L**) Foot swelling after CHIKV infection. Statistical analysis: **A-D, G, J,** and **L,** two-way ANOVA with Dunnett’s (**A-D** and **L**) or Šídák’s (**G** and **J**) post-tests; AUC analyses were performed with one-way ANOVA with Dunnett’s post-test (**A-D** and **L**) or unpaired t-test (**G** and **J**); mean values ± SEM are shown. **E-F, K**, one-way ANOVA with Šídák’s multiple comparisons test. **H,** unpaired t-test. **** *P* < 0.0001; *** *P* < 0.001; ** *P* < 0.01; * *P* < 0.05; ns, not significant.

We also examined whether T and B cells contributed to the joint swelling phenotype seen after microbiota perturbation due to antibiotics. In *Rag1^-/-^* mice lacking mature T and B cells, AV-treatment lost its effect on CHIKV-induced foot swelling (**Fig S4A**). This phenotype was due to a requirement for T cells, as AV-treated *Tcrbd^-/-^* mice (lacking both αβ and γ8 T cells) also did not develop increased foot swelling (**Fig S4B**). In contrast, AV-treated μMT mice that have T cells, but lack B cells, developed worsened foot swelling after CHIKV infection (**Fig S4C**), like wild-type mice (**Fig 1G**).

To determine the relative contribution of CD4^+^ or CD8^+^ T cells, we depleted these populations with mAbs (**Fig S4D**). Depletion of CD8^+^ T cells had no effect on the AV-mediated swelling phenotype or joint infiltrates. However, depletion of CD4^+^ T cells suppressed foot swelling and decreased myeloid cell infiltration in AV-treated animals to levels comparable to water-treated, CHIKV-infected mice (**Fig 8D-E**). T cell depletion did not affect the viral load in the ipsilateral feet of antibiotic-treated mice at 4 dpi (**Fig 8F**).

We next assessed whether CHIKV-specific CD4^+^ T cells were required for the exacerbated joint-associated swelling by administering antibiotics to OT-II *Rag1^-/-^* mice before infection with CHIKV. In these T cell receptor (TCR) transgenic mice, virtually all CD4^+^ T cells express αβ-TCRs against chicken ovalbumin H-2^b^-restricted peptide 323-339 ^63,64^, and there are no CD8^+^ T cells (**Fig S4E**). Like what we observed in wild-type mice, AV-treated OT-II *Rag1^-/-^*mice developed enhanced foot swelling compared to untreated control mice (**Fig 8G**), and joint-associated CD4^+^ T cells were capable of producing inflammatory cytokines (**Fig S4F**), suggesting that the early antibiotic-driven inflammation in the joint is mediated at least in part by antigen-independent, bystander CD4^+^ T cells. These data also suggest that the tissue swelling after AV treatment likely is not due to autoreactive CD4^+^ T cells and molecular mimicry between microbial products and musculoskeletal tissue-associated self-antigens. In contrast, reduced swelling was seen at 7 dpi and thereafter, like that observed in AV-treated *Rag1^-/-^*, *Tcrbd^-/-^*, or CD4^+^ T cell depleted mice (**Fig 8D and S4A-B**), due to a primary role for CHIKV-specific CD4^+^ T cells in mediating the second peak of swelling.

### Gut-associated immune cells are required for increased CHIKV-induced joint swelling in antibiotic-treated mice

The gut microbiota can modulate local and peripheral Treg ^46–48^ and effector T cell ^65–68^ responses, and SCFA can induce Treg populations in certain contexts ^46–48,69^. As mentioned above, we did not detect differences in Treg percentages or numbers in the ipsilateral popliteal lymph nodes or spleens of water- and AV-treated mice at 4 dpi (**Fig 3F-I**). Furthermore, we did not observe differences in Tregs in the spleens of AV-treated mice prior to infection (**Fig S4G**). To evaluate this question further, we examined the effects of AV treatment on Treg levels and cytokine production by T cells in the mesenteric lymph nodes (MLN) that drain the intestines. AV-treated mice showed a reduced proportion of Tregs in the MLN prior to CHIKV infection but not at 4 dpi (**Fig S4H**). AV-treated mice also had increased proportions of MLN CD4^+^ T cells producing IFN-γ and IL-17 (**Fig S4I**). Similarly, AV treatment led to a modest increase in the proportion of colonic lamina propria CD4^+^ T cells producing IFN-γ, IL-17, and IL-22 (**Fig 8H and S4J-K**) without altering the proportions of iNOS- and Arg1-expressing macrophages (**Fig S4L**). These findings suggest that AV-mediated dysbiosis, marked by loss of SCFA and increased gut permeability, alters mucosal CD4^+^ T cell responses.

CD4^+^ T cells from the gut can migrate to distal sites to promote inflammation ^70^. To determine the sufficiency of AV-primed gut-associated CD4^+^ T cells in mediating enhanced joint inflammation after CHIKV infection, we adoptively transferred CD4^+^ T cells enriched from MLNs and Peyer’s patches of water or AV-treated uninfected mice into recipient *Tcrbd^-/-^* mice (**Fig 8I**). Upon subsequent CHIKV infection one day later, *Tcrbd^-/-^* mice that received CD4^+^ T cells from AV-treated mice had more joint swelling than those receiving similarly isolated T cells from water-treated animals (**Fig 8J**). This suggests that gut-associated CD4^+^ T cells primed by antibiotic-treatment can promote enhanced joint inflammation during CHIKV infection.

Finally, we investigated whether preventing immune cell migration to gut-associated tissues could reduce the antibiotic-induced exacerbation of CHIKV-related joint swelling. Mucosal addressin cell adhesion molecule-1 (MAdCAM-1), expressed on endothelial cells in the intestinal lamina propria, Peyer’s patches, and MLNs, is upregulated by pro-inflammatory cytokines ^71-74^, and acts as the primary ligand for α4β7 integrin-mediated immune cell homing to gut-associated tissues ^75–80^. Antibody blockade of MAdCAM-1 starting two days prior to initiation of oral antibiotics reduced immune cell accumulation in the colonic lamina propria (**Fig 8K**) and MLNs (**Fig S4M**), and attenuated CHIKV-induced joint swelling to levels comparable to that of isotype mAb and water-treated mice (**Fig 8L**). Together, these data show that immune cells primed in the gut-associated compartment contribute to enhanced CHIKV joint inflammation in antibiotic-treated mice.

### Oral antibiotics alter the transcriptional signature of CD4^+^ T cells, infiltrating monocytes, tissue-resident macrophages, and osteoclast-like cells in the joint

A major gap in our understanding of the gut-joint axis is how dysbiosis affects the interplay between tissue-resident and infiltrating immune cells that drive synovial inflammation and injury. To gain insight into this question, we performed single-cell RNA sequencing of sorted CD4^+^ T cells, monocytes, and macrophages from the ipsilateral joint-associated tissues of mice infected with MAYV infection that received (1) AV treatment, (2) AV treatment and oral propionate supplementation, (3) AV treatment and CD4^+^ T cell depletion, or (4) water (control) (**Fig S5A-B**). We used this alphavirus because it retains the joint swelling phenotype after AV treatment (**Fig S1H**) and enabled cell sorting at BSL2.

Within joint-associated tissues of MAYV infected mice, we identified three transcriptionally distinct populations of CD4^+^ T cells: Tregs (*Ikzf2^+^ Foxp3^+^ Ctla4^+^*); activated T cells (*Il18r1^+^ Il12rb2^+^ Themis^+^ Nkg7^+^*); and cycling T cells (*Pclaf^+^ Top2a^+^ Stmn1^+^ Mki67^+^ Cdk1^+^*) (**Fig 9A and S5C-D**). We also identified four populations of infiltrating monocytes and macrophages, including two clusters of *Ly6c^hi^ Ccr2^+^ Il1b^+^* monocytes, one cluster of *Ly6c^low^ Ccr2^+^ Il1b^+^* monocytes, and a population of *Ccr2^+^ Cd74^+^ Il1b^+^* macrophages (**Fig 9A-B and S5E**). Our analysis also revealed populations of tissue-resident synovial lining macrophages (*Cx3cr1^+^ Trim69^+^ Fap^+^*), interstitial macrophages (*Cx3cr1^-^ Trem2^+^ Folr2^+^ Aqp1^+^ Alox5^+^*), and osteoclast-like cells (*Tnfrsf11a^+^ Ctsk^+^ Atp6v0d2^+^ Mmp14^+^ Gpnmb^+^*) (**Fig 9A-B and S5E**). These cell types are implicated in mediating the bone erosion and joint destruction associated with alphavirus-induced arthritis ^18–21,81–84^.

**Figure 9.**
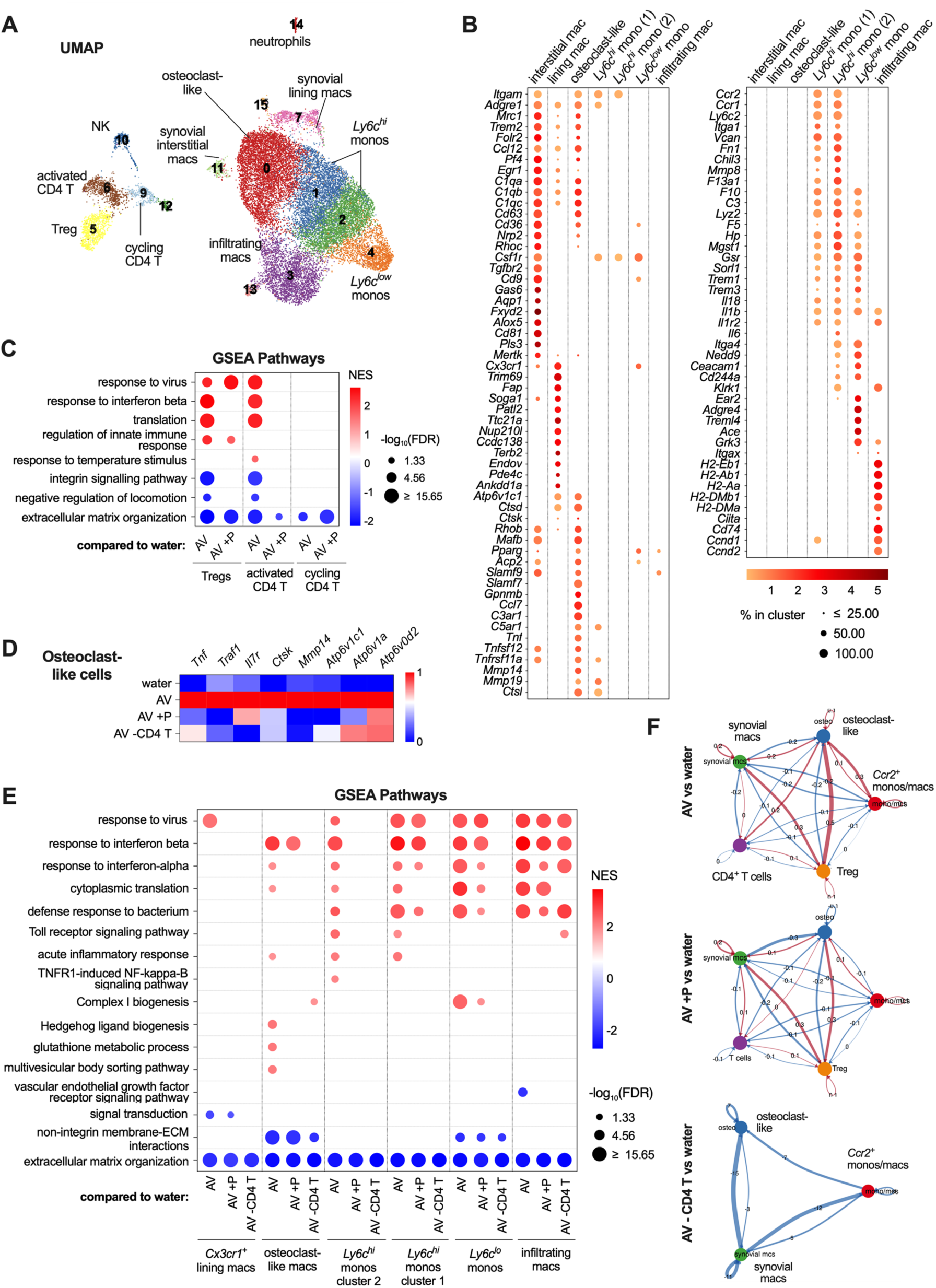
Oral antibiotics alter the transcriptional signature of tissue-resident and infiltrating immune cells in the virus-infected joint. (**A**) UMAP visualization of sort-enriched CD4^+^ T and myeloid cells from single cell-RNA sequencing data. (**B**) Bubble plot showing patterns of gene expression among synovial interstitial and lining macrophages, osteoclast-like cells, and infiltrating *Ccr2^+^* monocytes and monocyte-derived macrophages. (**C**) GSEA pathway analysis of differentially expressed genes in Tregs, activated CD4^+^ T cells, and proliferating CD4^+^ T cells from mice subjected to AV-treatment, AV-treatment with propionate supplementation, and AV-treatment with CD4^+^ T cell depletion, compared to corresponding cells from water-treated control mice. (**D**) Heatmap showing expression of osteoclast-related genes in osteoclast-like cells from mice treated with water, AV, AV with propionate supplementation, and AV with CD4^+^ T cell depletion. (**E**) GSEA pathway analysis of differentially expressed genes in tissue-resident and infiltrating monocyte and monocyte-derived macrophage populations from mice subjected to AV-treatment, AV-treatment with propionate supplementation, and AV-treatment with CD4^+^ T cell depletion, compared to water-treated control mice. (**F**) Differential interaction strength between CD4^+^ T cells, tissue-resident synovial macrophages, infiltrating monocytes and monocyte-derived macrophages, and osteoclast-like cells. The width of the line indicates strength of interaction. Red coloring indicates increased interactions between cell populations in the different AV-treated groups, whereas blue coloring indicates increased interaction in the water-treated comparator group.

Compared to the corresponding CD4^+^ T cells from water-treated animals, both Tregs and activated *Il18r1^+^ Il12rb2^+^ Nkg7^+^* T cells from joint-associated tissues of AV-treated mice displayed enhanced inflammatory signatures and reduced expression of genes related to integrin signaling and locomotion (**Fig 9C**). These transcriptional signatures were largely reversed in AV-treated mice that received oral propionate (**Fig 9C**), consistent with our observation that SCFA treatment reduces pro-inflammatory cytokine production by joint-associated CD4^+^ T cells.

Osteoclast-like cells from AV-treated mice exhibited increased expression of osteoclast-related genes including *Ctsk, Atp6v1c1, Atp6v1a, Mmp14, Tnf, and Il7r* (**Fig 9D**), as well as genes associated with inflammatory responses, Hedgehog ligand biogenesis, glutathione metabolic process, and multivesicular body sorting (**Fig 9E**), all of which are induced during osteoclast differentiation and bone homeostasis ^85–94^. *Ccr2^+^* infiltrating monocytes and monocyte-derived macrophages from AV-treated mice also showed a more prominent inflammatory response including increased expression of IFN-stimulated genes (ISGs), TNF family signaling pathways, and gene patterns induced by bacteria (**Fig 9E**). While substantial transcriptional changes were not observed in interstitial macrophages, *Cx3cr1^+^* synovial lining macrophages from AV-treated mice demonstrated an increased ISG response (**Fig 9E**). Propionate supplementation of MAYV-infected AV-treated mice largely reversed these myeloid cell changes, particularly in synovial lining macrophages, osteoclast-like cells, and some infiltrating monocytes (**Fig 9D-E**). Together, these data demonstrate that loss of bacterially derived SCFA promotes pro-inflammatory transcriptional programs in both joint-resident and infiltrating CD4^+^ T and myeloid cells, including differential upregulation of genes involved in osteoclast differentiation and activation, which can be rescued with oral SCFA supplementation.

### Increased inflammatory signature in myeloid cells in the joint is driven by CD4^+^ T cells

CD4^+^ T cells with IL-12R and IL-18R co-expression are enriched in inflamed synovial tissue from patients with rheumatoid arthritis ^95^, and activation of both receptors synergizes to induce a robust IFN-γ response ^96^. T and myeloid cell crosstalk in the joint can amplify inflammation and promote osteoclast differentiation ^97^. Given the requirement for CD4^+^ T cells in driving AV-mediated exacerbation of CHIKV arthritis, as well as our identification of activated *Il18r1^+^ Il12rb2^+^ Nkg7^+^* CD4^+^ T cells in joint-associated tissues of MAYV-infected mice, we assessed the contributions of CD4^+^ T cells to the inflammatory signatures of joint-associated macrophages. Indeed, synovial macrophages, infiltrating monocytes, and osteoclast-like cell populations did not show the increased ISG or pro-inflammatory signatures in AV-treated, MAYV-infected mice lacking CD4^+^ T cells (**Fig 9D-E**). Among *Ccr2^+^*infiltrating monocytes, depletion of CD4^+^ T cells had an even stronger effect than propionate supplementation in reversing the AV-driven transcriptional signature, suggesting that the antibiotic-mediated effects on tissue-resident and infiltrating myeloid cells in the joint is mediated principally by CD4^+^ T cells.

Finally, we assessed for potential networks of myeloid and CD4^+^ T cell interactions and crosstalk between synovial macrophages, infiltrating monocytes and macrophages, osteoclast-like cells, Tregs, and activated/cycling T cells in the joint tissues of MAYV-infected mice. Cell-to-cell interaction modeling revealed CD4^+^ T cell communication with myeloid cells in the joint via CCL5-, IFN-γ-, FasL, and adhesion molecule receptor-mediated interactions (**Fig 10A**). Activated and cycling CD4^+^ T cell interactions also promote an osteoclastogenic phenotype in infiltrating monocytes/macrophages and osteoclast-like cells through BST2, TSP-1 (encoded by *THBS1*), and MIF interactions with their respective receptor ligands (**Fig 10A**), which mediate osteoclast differentiation and function ^98–103^. CellChat-based interaction modeling predicted increased interaction strength of CD4^+^ T cells with osteoclast-like cells and of infiltrating monocytes with osteoclast-like cells in AV-treated compared to water-treated mice (**Fig 9F**). These effects were diminished in AV-treated mice supplemented with propionate or depleted of CD4^+^ T cells (**Fig 9F**). As an example, BST2-PIRA2 interactions, which provide critical costimulatory signals for RANK-induced osteoclast differentiation ^102,103^, were upregulated in AV-treated mice (**Fig 10A-B**) but reduced in AV-treated mice that received propionate (**Fig 10C**). Taken together, our transcriptional profiling data suggest the following: (1) AV-mediated dysbiosis skews CD4^+^ T cells towards an inflammatory signature that activates infiltrating myeloid cells, resident synovial macrophages, and osteoclast-like cells; and (2) this inflammatory phenotype is related to loss of certain SCFA, such as propionate, and can be reversed with oral propionate supplementation. Thus, we link specific gut microbe-derived SCFA to the integrity of the intestinal barrier, immune cell priming, modulation of resident synovial macrophages and osteoclasts in the joint, and control of virally induced tissue inflammation and arthritis.

**Figure 10.**
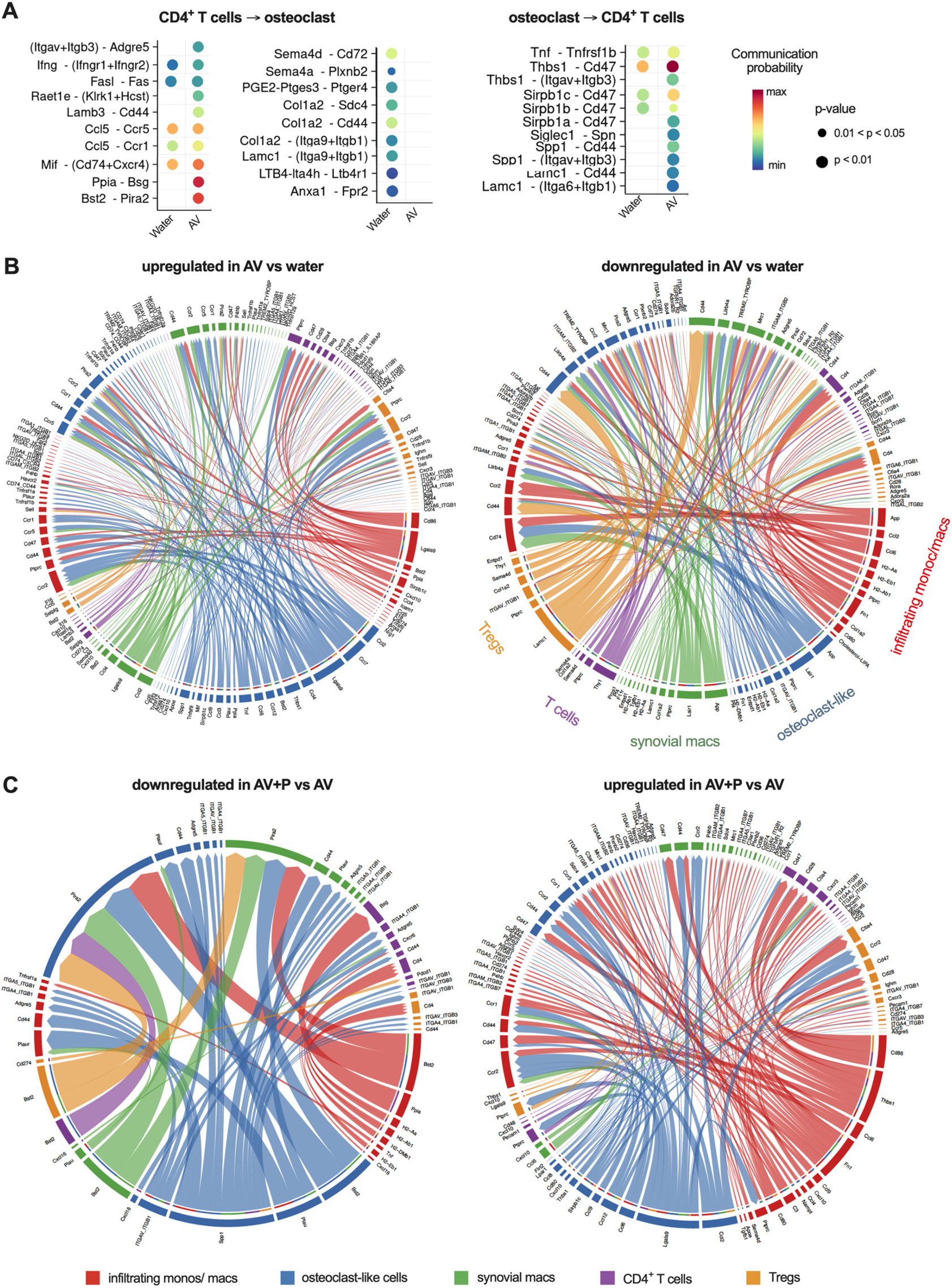
Predicted cell-cell interactions between joint-associated CD4^+^ T and myeloid cells. (**A**) Predicted T cell-osteoclast-like cell interactions based on modeling using CellChat. (**B-C**) Chord diagrams of ligand-receptor interactions among Tregs, CD4^+^ T cells, synovial macrophages, osteoclast-like cells, and infiltrating monocytes/macrophages. Shown are up- and down-regulated interactions in the joint tissues of MAYV-infected mice that received (**B**) AV compared to water, and (**C**) AV plus propionate compared AV only.

## DISCUSSION

In this study, we establish that perturbation of the intestinal microbiota due to antibiotic treatment durably exacerbates musculoskeletal inflammation during alphavirus infection and that the underlying mechanism is distinct from previously reported antiviral activities of gut bacteria (*e.g*., homeostatic induction of ISGs, bile acid-mediated priming of type I and III IFN, or effects on antiviral CD8^+^ T cell responses) ^32,65,104,105^. Using bacterial and metabolite reconstitution approaches, we link exacerbated alphavirus-induced joint inflammation to intestinal deficiencies of bacterial-derived SCFA, altered intestinal permeability, and pro-inflammatory effects of CD4^+^ T cells. Moreover, we determined that in the context of microbiota perturbation, MyD88 signaling in both intestinal epithelial cells and monocytes regulates the accumulation of inflammatory monocytes and polarized CD4^+^ T cells in the joint, which promotes differentiation of osteoclast-like cells. Our work establishes that intestinal microbes and their metabolites can affect intestinal epithelium and immune cells to regulate the severity of alphavirus-induced inflammation at a distant site, in the joint.

Despite evidence supporting a gut-joint link ^7,9,10^, the mechanisms driving these connections remain challenging to define. In humans, intestinal inflammation and infections can trigger joint-associated autoimmune inflammatory diseases including reactive and inflammatory bowel disease-associated arthritis ^106^. Although associated with HLA-B27 major histocompatibility antigen alleles, the IL-23/IL-17 axis, and tissue-specific adhesion receptor expression on immune cells, the exact mechanisms underlying joint inflammation in these conditions are still unclear ^107^. Moreover, gut dysbiosis and loss of butyrate-producing bacteria have been observed in rheumatoid arthritis ^108–110^; these changes are thought to mediate pathological effects via disruption of intestinal barrier function, dysregulation of pro-inflammatory cytokines, T follicular helper responses, and in some cases, autoantibody production. Treatment of rheumatoid arthritis with disease modifying anti-rheumatic drugs or agents that neutralize TNF activity can modulate dysbiosis ^111–113^, suggesting that systemic inflammation can drive alterations of the gut microbiota that amplify inflammation. In a similar vein, CHIKV infection of rhesus macaques alters the composition of the gut microbiota and their metabolites, which is associated with upregulation of pro-inflammatory gene expression in GI tract tissues including the inflammasome and IL-17 pathways ^114^. In contrast, in a microplastic ingestion model in mice, a mild inflammatory signature developed in the colon and MLN, without affecting the microbiome, and this was associated with increased CHIKV foot swelling ^115^. As global antibiotic consumption has continued to rise, its use has been linked to an increased risk for the onset or relapse of inflammatory rheumatoid arthritis and juvenile idiopathic arthritis ^1–6^. Nevertheless, the basis for these associations also is not understood.

Our experiments demonstrated that antibiotic-induced gut dysbiosis resulted in increased immune cell infiltration and foot swelling after CHIKV infection, which was driven by pro-inflammatory monocytes and both bystander and antigen-specific effector CD4^+^ T cells. Although monocytes and CD4^+^ T cells are known to modulate CHIKV-induced joint inflammation ^22,25–27,83,116–118^, our study revealed a distinct requirement for pro-inflammatory CD4^+^ T cells in exacerbating alphavirus-induced arthritis in the context of gut dysbiosis. This finding highlights an important role of the commensal gut microbiota and its metabolites in shaping the effector phenotypes of T cells during arthritis, affecting both infiltrating and synovial tissue-resident macrophages. In contrast to studies that used prolonged courses of broader-spectrum antibiotics, we did not observe a reduction in peripheral Tregs following a 3-day pulse of oral AV or subsequent CHIKV infection. Rather, a short-term dysbiosis led to increased frequencies and numbers of pro-inflammatory CD4^+^ T cells and infiltrating monocytes in joint-associated tissues after CHIKV infection. Treatment with neutralizing antibodies against TNF or IL-17A improved joint swelling in AV-treated mice, while IFN-γ neutralization showed a partially protective effect.

Oral antibiotics exacerbated CHIKV arthritis and myositis in a MyD88-dependent manner. This phenotype was at least partially mediated by gut epithelial-specific MyD88 signaling, indicating that enhanced joint inflammation induced by these effector T cells is linked to events at the gut mucosal interface. The effect was partially dependent on TLR4 signaling, as no significant change in joint-associated swelling was observed in antibiotic-treated mice with individual deficiencies in IL-1 receptor, TLR2, TLR7, or TLR9. Notably, *Myd88*^-/-^ mice showed a more profound reduction in swelling compared to any single TLR-deficient mouse, suggesting that multiple PAMPs induced by gut dysbiosis might contribute to inflammation, which could potentially mask the effects of individual TLRs. Alternatively, other TLRs or MyD88-dependent non-TLR signaling pathways might also play a role.

MyD88 signaling in the intestinal epithelium is essential in gut homeostasis through its function in maintaining the epithelial barrier, intestinal mucus production, and mucosa-associated antimicrobial activity ^53–56^. Our results show that, in the setting of microbial perturbation, MyD88 signaling in enterocytes promotes alphavirus-induced musculoskeletal inflammation, possibly through effects on immune cell activation like those described for ischemia/reperfusion-induced intestinal injury ^119^. In line with this, other studies using *Villin*-Cre *Myd88^f/f^*mice have shown disease amelioration in conditions such as renal injury ^59^, diet-induced obesity, and hepatic steatosis ^58^, although the specific contributions of bacteria or their products were not characterized.

Disruption of the gut barrier can lead to inflammation not only within gut-associated compartments but also in peripheral organs, through mechanisms such as the migration of activated immune cells or translocation of PAMPs into circulation. These PAMPs may be sensed by peripheral immune cells—such as monocytes—via both MyD88-dependent and independent pathways to potentially amplify inflammation. Our findings that selective MyD88 deletion in monocytes only minimally ameliorates the worsened joint inflammation associated with dysbiosis, and that CD4^+^ T cells promote monocyte infiltration and activation in the joint, suggest that monocyte-intrinsic MyD88 signaling pathway plays only a partial role in driving musculoskeletal inflammation during CHIKV infection. Further investigation is needed to define how MyD88 signaling in enterocytes transmits luminal signals from the gastrointestinal tract to shape effector T cell function and inflammation both locally and systemically.

We identified individual species of anaerobic bacteria (*e.g*., C*. scindens* and *B. thetaiotaomicron*) that can mitigate the deleterious effects of microbiota depletion on musculoskeletal inflammation during CHIKV infection. We showed that the protective effect of *B. thetaiotaomicron* is mediated through production of its primary SCFA, propionate. It is likely that multiple bacterial species across different phyla contribute to anti-inflammatory effects during systemic viral infection via the production of immunoregulatory metabolites. Although exogenous butyrate has previously been shown to regulate intestinal barrier integrity and reduce disease severity in mouse models of sterile arthritis ^120–122^, the upstream and downstream mediators were not defined. Our findings establish a distinct role for an endogenously produced SCFA in modulating virus-induced arthritis and define the epithelial and immune cell interactions—as well as both cell-intrinsic and extrinsic signaling pathways—that link the gut microbiota to joint-associated inflammation.

In the context of viral arthritis, SCFA may act locally at the intestinal interface or translocate into the circulation to influence immune cells at distal sites of inflammation. Locally, SCFA help maintain gut barrier integrity and induce intestinal Tregs ^46,47^. Circulating SCFA also can support the generation of peripheral Tregs ^48^ and modulate inflammation at extra-intestinal sites, including the lung, brain and joint, as described in models of allergic airway inflammation ^123,124^, autoinflammatory lung disease ^125^, experimental autoimmune encephalomyelitis ^126^, and sterile arthritis ^120,122^. In contrast to reports showing that oral butyrate exacerbates CHIKV arthritis in mice^49^ or exerts anti-inflammatory effects in models of allergic and inflammatory lung disease ^123,125,127^, we did not observe either aggravating or protective effects of exogenous SCFA on CHIKV-induced joint inflammation when the microbiota was intact. The difference in CHIKV arthritis outcomes may reflect dose-dependent effects of oral butyrate (200 mM in the prior study ^49^ versus 37 mM in our study) that regulate gene expression and immune function in distinct ways ^69,128^: low doses of butyrate promote colonocyte proliferation and barrier function, whereas higher doses can increase intestinal epithelium permeability and inflammation ^129–133^. Finally, our experiments showed that (1) CD4^+^ T cells from MLNs and Peyer’s patches of antibiotic-treated mice can transfer an enhanced CHIKV-induced joint swelling phenotype to untreated *Tcrbd^-/-^* recipients, and (2) blocking immune cell homing to gut-associated tissues prior to antibiotic treatment reduces the impact of antibiotics on joint inflammation. These data collectively support the conclusion that activated gut-associated immune cells— including CD4^+^ T cells—are necessary and sufficient to drive joint inflammation in the context of gut dysbiosis.

Macrophages are key regulators of tissue homeostasis, inflammation, and cartilage/bone damage ^134–138^. Notably, changes in synovial macrophage phenotypes often precede the onset of clinical disease in inflammatory arthritis ^135,136^. Bidirectional crosstalk between T cells and monocytes/macrophages within the joint microenvironment can further amplify inflammation and arthritis severity ^97^. However, to our knowledge, how the gut microbiota modulates synovial macrophage subsets and immune interactions in the joint had not been explored. We found that antibiotic-induced loss of SCFA led to increased infiltration of pro-inflammatory monocytes/macrophages into alphavirus-infected joints, an enhanced ISG signature in CX3CR1^+^ synovial lining macrophages, and an expansion and transcriptional activation of osteoclast-like cells, the latter of which are associated with erosions and bone effacement, a feature of alphavirus infection in joint-associated tissues ^18–21,81–84^. These pathological changes were ameliorated by propionate supplementation or depletion of CD4^+^ T cells, highlighting the critical role of microbial metabolites, activation of immune cells in the gut-associated compartment, and T cell-macrophage interactions in regulating joint inflammation and tissue damage.

In summary, our study provides mechanistic insight into how antibiotics disrupt the relationship between commensal gut microbes, microbial-derived metabolites that maintain epithelial integrity, and the immune pathways that regulate tissue inflammation in acute arthritis caused by globally emerging alphaviruses. Multiple bacterial species across different phyla produce similar metabolites (*e.g*., SCFA) that help maintain an equilibrium between immune tolerance and host defense, which can be disrupted by even by short-term oral antibiotics. Our work further reveals an underrecognized role for gut microbe-derived SCFA in restraining the polarization of both effector and bystander CD4^+^ T cells. This, in turn, prevents the accumulation of inflammatory monocytes, the pro-inflammatory reprogramming of synovial tissue-resident macrophages, and enhanced differentiation of osteoclast-like cells within joints during viral arthritis. Together, these results highlight the gut microbiota, their metabolites, and downstream host signaling pathways as potential therapeutic targets for virus-induced and non-infectious inflammatory joint diseases.

## ACKNOWLEDGMENTS

This study was supported by grants and contracts from NIH (R01 AI152484 to M.S.D., R01 DK132244 and R01 DK135816 to C.J.G.), an award from the Systems Immunology Initiative by Department of Pathology & Immunology of Washington University School of Medicine Department (to M.N.A); the Washington University Musculoskeletal Research Center (NIH P30 AR074992); and by a pilot award from the American Gastroenterological Association, the W.M. Keck Foundation, and the RAPP funding from the Weill Cornell Medicine (all to C.J.G.). F.R.Z was supported by a Rheumatology Research Foundation Scientist Development Award, T32 CA009547-35, and by K08 AR084597. E.S.W. was supported by F30 AI152327. We thank Michelle Elam Noll posthumously for assistance with animal husbandry. We acknowledge the Washington University Gnotobiotic Core Facility for assistance with germ-free mouse studies.

## AUTHOR CONTRIBUTIONS

Data acquisition: F.R.Z., M.K, E.S.W., H.J., R.B.W. C.J.G., T.T.L., L.W., A.J., L.D. and L.H. Analysis and interpretation of data: F.R.Z., M.K., E.S.W., C.J.G., R.W., S.H., M.F., and M.S.D. Provided critical reagents and input on experimental design: C.S.H., M.M., and M.T.B. Research supervision: M.A., T.S.S., M.F., and M.S.D. F.R.Z. and M.S.D. wrote the initial draft, with the other authors providing editorial comments.

## MATERIALS AND METHODS

### Viruses

Stocks of CHIKV La Reunion OPY1 (CHIKV-LR 2006) were generated from infectious cDNA clones using published methods ^139^. MAYV BeH407 was obtained from the World Reference Center for Emerging Viruses and Arboviruses (R. Tesh, K. Plante, and S. Weaver, University of Texas Medical Branch, Galveston, TX) ^140^. Virus was grown from P0 stocks and titrated by focus-forming assays in Vero CCL-81 cells according to published protocols. Experiments with CHIKV and MAYV were performed in certified BSL-3 and BSL2 facilities, respectively.

### Conventionally housed mice

Housing and care of laboratory animals was conducted in accordance with guidelines from the NIH Guide for the Care and Use of Laboratory Animals. Animal husbandry and experiments were performed in accordance with guidelines from the Institutional Animal Care and Use Committee at Washington University in St Louis School of Medicine (Assurance number A3381-01). Specific pathogen-free male C57BL6/J mice (000664) were obtained from Jackson Laboratory. *Rag1^-/-^* (*Rag1^tm1Mom^*, 034159)*, Tcrbd^-/-^* (*Tcrb^tm1Mom^ Tcrd^tm1Mom^*, 002122)*, µMT* (*Ighm^tm1Cgn^*, 002288)*, Myd88^-/-^* (*MyD88^tm1.1Defr^*, 009088)*, Il1r^-/-^* (*Il1r1^tm1Imx^*, 003245)*, Tlr7^-/-^* (*Tlr7^tm1Flv^*, 008380)*, Tlr9^-/-^* (*Tlr9^M7Btlr^*, 34329-Jax)*, Tlr2^-/-^*(*Tlr2^tm1Kir^*, 004650)*, Tlr4^-/-^*(*Tlr4^tm1.2Karp^*, 029015), and *Pou2f3^-/-^* (*Pou2f3^em1Cbwi^*, 037040) mice were purchased from the Jackson Laboratory and bred as heterozygotes in facilities at Washington University to generate KO and WT littermate control mice. CD45.1 OT-II *Rag1^-/-^* mice were generated and maintained using homozygote crosses. Mice lacking MyD88 expression in intestinal epithelial cells and monocytes were generated by crossing *Myd88^f/f^* mice (008888) with *Villin*-Cre (004586) or *Ccr2-*Cre-ER^T2^ mice (035229), respectively, which were obtained from the Jackson Laboratory. All mice were housed in enhanced caging that were pre-assembled prior to autoclaving, provided an autoclaved 20% protein diet and water ad libitum, and maintained on a 12 h light/dark cycle.

### Gnotobiotic Mice

C57BL/6J GF mice were bred and maintained at the Washington University Gnotobiotic Core Facility. GF status was confirmed by 16S qPCR analysis (Charles River).

### Virus infection and assessment of foot swelling

3-4-week-old mice were anesthetized with ketamine hydrochloride (80 mg/kg) and xylazine (15 mg/kg). For infections, mice were inoculated subcutaneously with 10^3^ focus-forming units (FFU) of CHIKV-LR or 10^3^ FFU of MAYV in a 10 µL volume in the left rear footpad. Digital calipers were used to measure the mediolateral aspect of the tarsus and dorsal-ventral aspect of the ipsilateral foot.

### Histology and viral RNA *in situ* hybridization (ISH)

At 3 or 6 dpi, ipsilateral feet were collected after extensive perfusion with phosphate buffered saline (PBS) and 4% paraformaldehyde (PFA). Tissue was fixed for 24 h in 4% PFA, followed by decalcification in 14% EDTA free acid (Sigma-Aldrich) at pH 7.2 for 10 days and dehydration in 70% ethanol. Paraffin-embedded tissues were sectioned; hematoxylin and eosin, TRAP, and Toluidine blue staining were performed following standard procedures. Viral *in situ* hybridization was performed using RNAscope 2.5 (Advanced Cell Diagnostics) according to manufacturer’s instructions and the CHIKV probe (479501).

### Viral burden analysis

At 0, 4, or 7 dpi, ipsilateral feet were collected from CHIKV-infected mice following extensive perfusion with PBS. Tissues were weighed and homogenized in 500 µL PBS with zirconia beads in a MagNA Lyser (Roche), followed by centrifugation of homogenates at 10,000 rpm for 5 min at 4°C. Viral burden was determined by focus forming assay (FFA) on Vero CCL-81 cells as previously described ^141^. Alternatively, for some samples, RNA was extracted using a MagMAX *mir*Vana Total RNA isolation kit (Thermo Scientific) and a Kingfisher extraction machine (Thermo Scientific). MAYV RNA levels for tissue homogenates and viral stocks were determined using previously designed primer/probe sets for MAYV ^141^ by one-step quantitative reverse transcriptase PCR (qRT-PCR) with Taqman RNA-to-Ct 1-step kit (Thermo Scientific) on an ABI 7500 Fast Instrument using standard cycling conditions. Viral titers are expressed on a log_10_ scale as FFU equivalents per gram of tissue, determined by comparing MAYV RNA equivalents on a standard curve using RNA isolated from known viral stocks.

### Antibiotic treatment, oral SCFA supplementation, oral DCA supplementation, SCFA enemas, and tamoxifen administration

3-4-week-old conventionally housed mice were provided with water or antibiotics (0.5 g/L vancomycin and/or 1 g/L ampicillin) in drinking water *ad libitum* for 3 days. For oral SCFA supplementation, sodium butyrate, sodium propionate, or sodium acetate (37 mM, Sigma-Aldrich) were added to drinking water at the time of antibiotic initiation and continued after cessation of antibiotics. For DCA supplementation, sodium deoxycholate (0.2%, Sigma-Aldrich) was added to drinking water at the time of antibiotic initiation and continued after cessation of antibiotics. For Cre induction, tamoxifen (Sigma-Aldrich) was dissolved in safflower oil and administered to 3-4-week-old *Myd88*^f/f^ *Ccr2* Cre-ER^T2^ mice via intraperitoneal injection at a dose of 75 mg/kg daily for three days prior to initiation of antibiotics.

### T cell, monocyte, and neutrophil depletion or blockade of immune cell migration

For T cell depletion, mice were treated with 25 μg of anti-CD4 (clone GK1.5; BioXCell), anti-CD8 (clone 2.43; BioXCell), or isotype control mAb (clone LTF-2; BioXCell) via intraperitoneal injection every 3 days for 1 week prior to and up to 2 weeks after infection. For combined monocyte, eosinophil, and neutrophil depletion, mice were treated with 250 μg of anti-Ly6G/Ly6C (Gr-1; clone RB6-8C5) or an isotype control (clone LTF-2; BioXCell) via intraperitoneal injection one day prior to infection and every 2 days following infection for 1 week. For neutrophil-only depletion, mice were treated with 250 μg of anti-Ly6G (clone 1A8; BioXCell) or an isotype control (clone 2A3; BioXCell) via intraperitoneal injection one day prior to infection and every 2 days following infection for 1 week. For blockade of monocyte trafficking, mice were treated with 25 μg of anti-CCR2 (clone MC-21) ^142^ or an isotype control mAb (clone LTF-2; BioXCell) via intraperitoneal injection one day prior to infection and every 2 days following infection for 1 week. Depletion or blockade of trafficking was confirmed by analysis of immune cell populations in the blood using flow cytometry. For blockade of immune cell migration to gut-associated tissues, mice were treated with 100 μg of anti-MAdCAM-1 (clone MECA-367; BioXCell) or an isotype control mAb (clone 2A3; BioXCell) via intraperitoneal injection one day prior to initiation of antibiotics and then every 3 days after. Blockade of trafficking was confirmed by analysis of immune cell populations in the colonic lamina propria and MLN using flow cytometry.

### Cytokine Neutralization

Mice were treated with neutralizing mAbs against TNF (200μg: clone MP6-XT22; BioLegend), IFN-γ (100μg: clone H22; Leinco Technologies), IL-17A (100μg: clone 17F3; BioXCell), or isotype control mAb every 3 days starting at 1 week prior to infection and continued after infection.

### *In vivo* intestinal permeability assay

Intestinal permeability was measured *in vivo* by assessing translocation of dextran from the luminal GI tract into the blood. Mice were administered 100 µL of 60 mg/mL dextran of different molecular weights, each conjugated to different fluorescent dyes (CascadeBlue 10 kDa, Thermo; tetramethylrhodamine 70 kDa, Thermo; FITC 250 kDa, Sigma-Aldrich; FITC 2000 kDa, Sigma-Aldrich). Blood was collected into EDTA tubes after 2 and 4 h and centrifuged (1500g for 10 min at 4°C). Plasma was diluted 1:1 with PBS and plated in triplicates. Fluorescence was measured with a BioTek Synergy H1 plate reader, and concentrations were calculated based on a standard curve.

### Fecal microbial transfer, bacterial culture, and bacterial reconstitution

For FMT, cecal contents were diluted in PBS with 10% glycerol, homogenized, and passed through a 100 μm nylon filter. For bulk FMT-derived aerobic and anaerobic bacteria, FMT stocks were cultured in liquid Brain Heart Infusion (BHI) Media under aerobic or anaerobic conditions, respectively. *C. scindens* and *B. thetaiotaomicron* were cultured in BHI or chopped meat media broth under anaerobic conditions. For bacterial reconstitution experiments, mice were rested for 1 day after discontinuation of antibiotics prior to oral gavage with sterile PBS or with 10^8^ colony-forming units of bacteria in a 100 µL volume.

### Adoptive transfer of CD4^+^ T cells from MLN and Peyer’s patches into *Tcrbd^-/-^* mice

MLN and Peyer’s patches were collected from mice that received either water or AV for 6 days. Single cell suspensions were generated via mechanical dissociation through a 70 μm filter, and CD4^+^ T cells were purified by negative selection using the mouse CD4^+^ T cell isolation kit (Miltenyi Biotec) following manufacturer’s instructions. Isolated T cells from MLNs and Peyer’s patches were pooled, and 10^6^ CD4^+^ T cells from either water or AV-treated mice were adoptively transferred into *Tcrbd^-/-^*mice via retro-orbital injection. Mice were subsequently inoculated with CHIKV at one day after T cell transfer.

### Generation and characterization of *B. thetaiotaomicron* mutants

#### Vector assembly

To generate the propionate deficient mutant, the conventional double-crossover gene deletion method was utilized ^50^ to knock out *mmdA* genes (BT_2090 and BT_2091) of *B. thetaiotaomicron*. First, two fragments with a length of ∼1200 bp flanking the target gene were amplified and fused via fusion PCR. This fragment was assembled with a PCR-amplified *pExchange-tdk* ^50^ backbone using Gibson Assembly and transformed into *E. coli* S17 directly to form a suicide vector that is introduced into *B. thetaiotaomicron* by conjugation described below. All primers used in the cloning are listed in **Table S2**.

#### Introducing vectors by conjugation into B. thetaiotaomicron

The suicide vectors were introduced into *B. thetaiotaomicron* based on a published method ^143^. First, a single colony of *B. thetaiotaomicron Δtdk* (a strain with the *BT_2275* gene, which encodes a deleted thymidine kinase gene) was inoculated in 3 mL of TYGB broth culture in an anaerobic chamber at 37°C under an atmosphere consisting of 10% CO_2_, 5% H_2_, 85% N_2_ for 12 h. *E. coli* S17 harboring a suicide vector were grown in 3 mL of LB broth supplemented with carbenicillin (100 µg/mL) at 37°C with shaking at 225 rpm. After ∼10 to 12 h, when the OD600 of *E. coli* S17 reached 0.8-1.0, 3 mL of *E. coli* S17 culture was centrifuged at 1500 x g for 3 min. The supernatant was discarded, and the cell pellet was washed once with 1.5 mL of PBS (pH 7.4). The washed *E. coli* S17 cell pellet was resuspended in a 3-mL overnight culture of *B. thetaiotaomicron* and gently mixed by pipetting.

The mixture was passed through a 0.2 µm filter. The liquid was discarded, and the filter membrane with the mixture of donor and recipient cells was placed onto the surface of a pre-reduced TSAB plate. The plate was incubated in a 37°C incubator aerobically. After incubating aerobically at 37°C for 24 h, the filter membrane was soaked in 2 mL of pre-reduced TYGB medium. The cells on the filter were resuspended into the medium by vortexing. The mixture was then transferred into the anaerobic chamber, and 100 µL was plated onto a pre-reduced TSAB plate + 200 µg/mL gentamycin + 25 µg/mL erythromycin. Colonies with plasmid inserted into the genome typically appeared after 36 to 48 h. Eight of these colonies were picked and re-streaked on pre-reduced TSAB plate + 200 µg/mL gentamycin + 15 µg/mL thiamphenicol to isolate single colonies. After 1 to 2 days, a single colony of each single crossover recombinant was cultured into TYGB medium, grown overnight for 16-24 h, and combined and plated on TSAB plates containing FUdR (200 μg/ml). Ten FUdR-resistant colonies were re-streaked to single colonies and subject to diagnostic PCR using the primers listed in **Table S2**.

#### LC-MS analysis of propionate in liquid culture

Single colonies of wide-type or mutant bacterial strains were inoculated in BHI+ Medium and grown anaerobically at 37°C. After 24 h, supernatant of each sample was collected for LC-MS analysis. 20 µL of supernatant of liquid culture was mixed with 200 µL of SCFA derivatization solution (freshly prepared acetonitrile solution containing 1 mM 2,2′-dipyridyl disulfide (DPDS), 1 mM triphenylphosphine (TPP) and 1 mM 2-hydrazinoquinoline (HQ)). The resulting mixture was vortexed and incubated at 60°C for 1 h, centrifuged at 21,000 g for 20 min, and its supernatant was analyzed by LC-MS. We used the following solvent system for detection of SCFA derivatives: A: H_2_O with 0.1% formic acid; B: methanol with 0.1% formic acid. 1 μL of each sample was injected with a flow rate of 0.35 ml/min and a column temperature of 40°C. The gradient for HPLC-MS analysis was: 0 - 6.0 min 99.5%A – 70.0%A, 6.0 - 9.0 min 70.0%A – 2.0%A, 9.0 - 9.4 min 2.0%A – 2.0%A, 9.4 - 9.6 min 2.0%A – 99.5%A. Peaks were assigned by comparison with authentic standards.

#### In vitro growth curve of B. theta WT and mutant strain

Three single colonies of wide-type or mutant strains were inoculated in BHI+ Medium and anaerobically cultured for overnight at 37°C to reach late-log phase. Then, 2 µL of the culture was resuspended in 200 µL of BHI+ broth, loaded into a 96-well plate, and incubated anaerobically at 37°C in a Multiskan SkyHigh Microplate Spectrophotometer (Thermo Fisher). OD600 nm readings were recorded every 30 min for 30 h.

### Quantification of fecal *Bacteroides*

Fecal samples from mice treated with antibiotics (AV) or gavaged with sterile PBS or wild-type or mutant *B. thetaiotaomicron* were collected in sterile microcentrifuge tubes, stored on ice, and transferred to an anaerobic chamber within 1 h of collection. A volume of 500 μL of sterile PBS was added to each tube, and samples were subjected to mechanical disruption via vortexing and pipetting. Ten-fold serial dilutions were cultured on *Bacteroides* bile esculin agar and counted after 48 h to determine colony-forming units.

### Gas chromatography mass spectrometry for SCFA measurements

Cecal samples were collected and frozen on dry ice prior to storage at −80°C. Samples (30 mg) were suspended in 300 µL of extraction solvent (1-pentanol and H_2_SO_4_, with an internal standard of d7-butyric acid) and homogenized by vortexing and sonication. For standards, pentyl esters of d7-Butyric acid (Cayman Chemical Company), butyric acid (Alfa Aesar), propionic acid (Sigma-Aldrich), and acetic acid (Fluka), were prepared and subjected to the same extraction procedure. All samples were incubated for 2 h at 80°C while mixing at 1,200 rpm, before cooling to room temperature. 300 µL of 0.9% NaCl and 300 μL of hexane were added to each sample. Samples were then vortexed for 20 sec, mixed for 2 min at 1,200 rpm at room temperature, and then centrifuged for 2 min at 13,200 rpm. 100 μL of the organic phase (supernatant) was transferred to a gas chromatography (GC) vial with insert for analysis using 1 μL injections in triplicate. For GC mass spectrometric analysis, an Agilent 7890A GC equipped with a 5975c MS detector (He carrier gas) was used. Inlet settings: 250°C, 17.8 psi, total flow 14.2 mL/min, and split ratio 10:1. Column: Phenomenex Zebron ZB-5MSi Guardian fused silica capillary column (30 m x 0.25 mm x 0.25 μm film thickness), 1.2 mL/min flowrate, 17.8 psi, constant flow. Oven: initial temperature 85°C hold for 4 min, ramp 1 to 110°C at 1 °C/min hold for 0 min, ramp 2 to 300 °C at 85°C/min hold for 2 min, ramp 3 to 85 °C/min hold for 1.2 min, total run time 36.2 min. Mass spectrometry (MS) conditions: transfer line 300°C, MS source 230°C, MS quad 150°C electron energy 70.3 eV. Data were visualized as EIC with *m/z* 61, 75, 89, and 96 observed for characteristic fragments from pentyl acetate, pentyl propionate, pentyl butyrate, and d7-pentyl butyrate (IS), respectively.

### Flow cytometry

Single cell suspensions were obtained from spleens and lymph nodes by mechanical dissociation, followed by filtration through 70 µm nylon mesh filters. For colonic lamina propria, mice underwent terminal anesthesia and extensive perfusion with PBS. The colon was collected and flushed with cold PBS, and surrounding mesentery and fat were removed. The colon then was cut longitudinally and washed again in Hank’s balanced salt solution (HBSS) containing 15 mM HEPES. To remove the epithelium, tissue pieces were incubated in HBSS containing 1 mM DTT and 15 mM HEPES for 20 min at 37°C with rotation, followed by incubation in HBSS containing 5 mM EDTA and 15 mM HEPES for 30 min at 37°C with rotation. To isolate lamina propria cells, the remaining tissue pieces were rinsed by sequential swishing in Petri dishes containing HBSS with 15 mM HEPES, then digested in complete RPMI (10% FBS, 2 mM L-glutamine, 1 mM sodium pyruvate, 50 µM beta-mercaptoethanol, and penicillin/streptomycin) with 100 U/mL collagenase IV for 1 h at 37°C with rotation. Digested samples were filtered over 100 µm nylon mesh filters and centrifuged at 850 x g for 10 min. Immune cells in the lamina propria were enriched by 70/40% Percoll gradient centrifugation at 850 x g for 20 min (no brake) with cells collected from the interface. For ipsilateral foot specimens, tissues were grossly dissected after perfusion, followed by digestion in RPMI with 10% FBS, 15 mM HEPES, type I collagenase (Thermo; 2.5 mg/mL) and DNase (Sigma-Aldrich; 10 µg/mL) for 1 h at 37°C with gentle agitation, followed by filtration through a 70 µm nylon filter. Cells were stained with primary antibodies in PBS with 2% FBS. For intracellular staining, cells were fixed and permeabilized using the FoxP3/Transcription Factor Staining Buffer Set (eBioscience) following manufacturer’s instructions and stained overnight at 4°C with secondary antibody in permeabilization buffer. For intracellular cytokine analysis of T cells, cells were incubated *ex vivo* at 37°C for 4 h in the presence of brefeldin A (BD Biosciences) in RPMI supplemented with 10% FBS, 2 mM L-glutamine, 1 mM sodium pyruvate, and 50 µM beta-mercaptoethanol, prior to fixation and permeabilization as detailed above. In some experiments including assessment of CD107a-cycling), cells were stimulated with anti-CD3. Conjugated antibodies against CD107a were included in the medium during the incubation period as a measure of T cell degranulation. For intracellular cytokine analysis of T cells in the spleen and lymph nodes, cells were stimulated with 1 µg/mL of plate-bound anti-CD3 mAb or PMA (50 ng/mL)/ ionomycin (1 µM) at 37°C for 4 h in the presence of brefeldin A. Samples were processed on a BD LSRFortessa X-20 flow cytometer (BD Biosciences) and analyzed using FlowJo software.

### Cytokine and chemokine analysis

Foot tissues were collected at 0 and 4 dpi. Tissues were homogenized with zirconia beads in a MagNa Lyser instrument (Roche Life Science) in 500 µL of phosphate-buffered saline and clarified by centrifugation. Cytokine and chemokine analysis was performed by Eve Technologies using a Millipore Mouse Cytokine Array/Chemokine Array platform (virus was inactivated using 1% Triton-X-100). Alternatively, cytokines (IFN-γ, TNF, and IL-17A/F) were measured from tissue homogenates prior to virus inactivation by ELISA (Biolegend) following the manufacturer’s instructions.

### Cell sort enrichment of CD4^+^ T cells, monocytes, and macrophages

Mice were treated with water, AV, AV supplemented with propionate, or AV after CD4^+^ T cell depletion (4 mice per group) prior to infection with MAYV. At 5 dpi, tissues from the ipsilateral feet were grossly dissected, and single cell suspensions were obtained by digestion with gentle agitation as described above. Cells were blocked with TruStain FcX™ PLUS (anti-mouse CD16/32, Biolegend) and then stained with TotalSeq anti-mouse hashtag antibodies (Biolegend B0302, B0304, B0306, B0308; one distinct hastag per mouse sample in each group following manufacturer’s instructions) and fluorescently conjugated antibodies against cell surface antigens. After washing, individual samples were pooled according to experimental group and resuspended in sorting buffer (PBS with 1% FBS, 1mM EDTA, and 25 mM HEPES). Fluorescence activated cell sorting was performed using a BD FACSAria II flow cytometer (BD Biosciences). Sorted cell populations were pooled according to experimental group and resuspended in PBS with 0.04% BSA prior to processing for single cell RNA sequencing.

### Single cell RNA sequencing and analysis

The Cell Ranger multi pipeline version 8.0.1, available on the 10x Genomics website, was utilized for demultiplexing pulled samples and aligning reads to the mouse reference genome mm10-2020-A. On average, 92.9% of reads per sample were successfully mapped, with over 90.5% of reads having a quality score above q30. Cells assigned one single CMO hashtag were analyzed, resulting in 27,839 cells (9,251 + 5,825 + 7,681 + 5,082 for water [W], AV only [AV], AV + propionate [AV+P], and AV + CD4^+^ T depletion [AV-T] conditions, respectively) with a median cells per sample and 2,774 genes per cell. Per sample matrices were processed using the Seurat package (version 5.1.0) ^144^. For each sample, the miQC tool ^145^ was utilized to filter cells with high mitochondrial content and quality. Samples then were merged on total expression, followed by scaling to 10,000, and subsequent log normalization. The identification of 3,000 highly variable genes, considering mean expression, involved scaling the data with simultaneous regression of unwanted variation (UMI counts and the fraction of mitochondrial reads). These scaled counts served as input for PCA. Employing the IntegrateLayers function, we conducted harmony integration on the first 50 PCA components. The first 30 dimensions of the integrated data were utilized to construct a UMAP plot and for further analysis. The FindNeighbors function on integrated data was run to create an SNN (shared nearest neighbor) graph, assisting in the identification of 21 clusters by the FindClusters function with a resolution of 0.6. Markers for cell-type identification were determined using the Wilcoxon rank-sum test from the FindAllMarkers function. Genes expressed in more than 10% of cells with at least a 0.10-fold difference were considered for each cluster. Based on obtained signatures, we deleted clusters without marker genes and repeated all steps starting from data scaling. The resulting dataset, comprising 21,129 cells (5,528 + 4,908 + 6,598 + 4,095 for water, AV, AV + propionate, and AV + CD4^+^ T depletion conditions, respectively) in 16 clusters (resolution 0.6), was passed to FindMarkers function to perform differential expression testing between conditions of specific clusters. Gene set enrichment analysis (GSEA) ^146,147^ was performed to identify pathways represented by sets of differentially expressed genes in different cell populations for each condition (AV, AV + propionate, and AV + CD4^+^ T depletion) compared to water treatment. For heatmaps, expression values were aggregated by conditions and clusters by AverageExpression function and visualized using the Phantasus app ^148^.

### Cell-to-cell interaction modeling from single cell RNA sequencing

CellChat (version 2.1.2) ^149^ was employed to analyze changes in receptor-ligand interactions among joint-associated cells across different experimental conditions. Interaction networks were constructed for each condition, encompassing five cell groups: Ccr2^+^ infiltrating monocytes/macrophages (clusters 1, 2, 3, and 4), CD4^+^ T cells (clusters 6 and 9), CD4^+^ T regulatory cells (cluster 5), synovial macrophages (clusters 7 and 11), and osteoclast-like cells (cluster 0). Initially, ligands and receptors for each group were identified using the identifyOverExpressedGenes and identifyOverExpressedInteractions functions. Interactions were then inferred using the computeCommunProb function with the “triMean” parameter, followed by pathway-level analysis with the computeCommunProbPathway function. After filtering out communications with fewer than 10 cells, and aggregating networks, the merged dataset contained 607 (W), 646 (AV), 684 (AV + propionate), and 481 (AV + CD4^+^ T depletion) communication links. Differences in network-level interactions were visualized using the netVisual_diffInteraction function. Up- and down-regulated cell-cell communications were determined through a combination of identifyOverExpressedGenes and netMappingDEG.

## Statistical analysis

Statistical significance was assigned when *P* values were < 0.05 using GraphPad Prism version 9.3. Tests, number of animals, mean values, and statistical comparison groups are indicated in the Figure legends.

## Materials availability

All requests for resources and reagents should be directed to the Lead Contact author. This includes viruses, vaccines, and primer-probe sets. All reagents will be made available on request after completion of a Materials Transfer Agreement (MTA).

## Data and code availability

All data supporting the findings of this study are available within the paper and are available from the corresponding author upon request. This paper does not include original code. 16S rRNA sequencing data generated in this study is available at the European Nucleotide Archive (Project accession: PRJEB53328). Single cell RNA sequencing data is available at Gene Expression Omnibus (Project accession: GSE288863). Any additional information required to reanalyze the data reported in this paper is available from the lead contact upon request.

## REFERENCES

1. Arvonen, M., Virta, L.J., Pokka, T., Kroger, L., and Vahasalo, P. (2015). Repeated exposure to antibiotics in infancy: a predisposing factor for juvenile idiopathic arthritis or a sign of this group’s greater susceptibility to infections? J Rheumatol 42, 521–526. 10.3899/jrheum.140348.

2. Sultan, A.A., Mallen, C., Muller, S., Hider, S., Scott, I., Helliwell, T., and Hall, L.J. (2019). Antibiotic use and the risk of rheumatoid arthritis: a population-based case-control study. BMC Med 17, 154. 10.1186/s12916-019-1394-6.

3. Raisanen, L.K., Kaariainen, S.E., Sund, R., Engberg, E., Viljakainen, H.T., and Kolho, K.L. (2023). Antibiotic exposures and the development of pediatric autoimmune diseases: a register-based case-control study. Pediatr Res 93, 1096–1104. 10.1038/s41390-022-02188-4.

4. Horton, D.B., Scott, F.I., Haynes, K., Putt, M.E., Rose, C.D., Lewis, J.D., and Strom, B.L. (2015). Antibiotic Exposure and Juvenile Idiopathic Arthritis: A Case-Control Study. Pediatrics 136, e333–343. 10.1542/peds.2015-0036.

5. Kindgren, E., and Ludvigsson, J. (2021). Infections and antibiotics during fetal life and childhood and their relationship to juvenile idiopathic arthritis: a prospective cohort study. Pediatr Rheumatol Online J 19, 145. 10.1186/s12969-021-00611-4.

6. Nagra, N.S., Robinson, D.E., Douglas, I., Delmestri, A., Dakin, S.G., Snelling, S.J.B., Carr, A.J., and Prieto-Alhambra, D. (2019). Antibiotic treatment and flares of rheumatoid arthritis: a self-controlled case series study analysis using CPRD GOLD. Sci Rep 9, 8941. 10.1038/s41598-019-45435-1.

7. Jeyaraman, M., Ram, P.R., Jeyaraman, N., and Yadav, S. (2023). The Gut-Joint Axis in Osteoarthritis. Cureus 15, e48951. 10.7759/cureus.48951.

8. Fragoulis, G.E., Liava, C., Daoussis, D., Akriviadis, E., Garyfallos, A., and Dimitroulas, T. (2019). Inflammatory bowel diseases and spondyloarthropathies: From pathogenesis to treatment. World J Gastroenterol 25, 2162–2176. 10.3748/wjg.v25.i18.2162.

9. Zaiss, M.M., Joyce Wu, H.J., Mauro, D., Schett, G., and Ciccia, F. (2021). The gut-joint axis in rheumatoid arthritis. Nat Rev Rheumatol 17, 224–237. 10.1038/s41584-021-00585-3.

10. Gracey, E., Vereecke, L., McGovern, D., Frohling, M., Schett, G., Danese, S., De Vos, M., Van den Bosch, F., and Elewaut, D. (2020). Revisiting the gut-joint axis: links between gut inflammation and spondyloarthritis. Nat Rev Rheumatol 16, 415–433. 10.1038/s41584-020-0454-9.

11. Zaid, A., Burt, F.J., Liu, X., Poo, Y.S., Zandi, K., Suhrbier, A., Weaver, S.C., Texeira, M.M., and Mahalingam, S. (2021). Arthritogenic alphaviruses: epidemiological and clinical perspective on emerging arboviruses. Lancet Infect Dis 21, e123–e133. 10.1016/S1473-3099(20)30491-6.

12. Teng, T.-S., Kam, Y.-W., Lee, B., Hapuarachchi, H.C., Wimal, A., Ng, L.-C., and Ng, L.F.J.T.J.o.i.d. (2015). A systematic meta-analysis of immune signatures in patients with acute chikungunya virus infection. 211, 1925–1935.

13. Tanabe, I.S.B., Santos, E.C., Tanabe, E.L.L., Souza, S.J.M., Santos, F.E.F., Taniele-Silva, J., Ferro, J.F.G., Lima, M.C., Moura, A.A., Anderson, L., and Bassi, E.J. (2019). Cytokines and chemokines triggered by Chikungunya virus infection in human patients during the very early acute phase. Trans R Soc Trop Med Hyg 113, 730–733. 10.1093/trstmh/trz065.

14. Ninla-Aesong, P., Mitarnun, W., and Noipha, K. (2019). Proinflammatory Cytokines and Chemokines as Biomarkers of Persistent Arthralgia and Severe Disease After Chikungunya Virus Infection: A 5-Year Follow-Up Study in Southern Thailand. Viral Immunol 32, 442–452. 10.1089/vim.2019.0064.

15. Hoarau, J.J., Jaffar Bandjee, M.C., Krejbich Trotot, P., Das, T., Li-Pat-Yuen, G., Dassa, B., Denizot, M., Guichard, E., Ribera, A., Henni, T., et al. (2010). Persistent chronic inflammation and infection by Chikungunya arthritogenic alphavirus in spite of a robust host immune response. J Immunol 184, 5914–5927. 10.4049/jimmunol.0900255.

16. Chang, A.Y., Tritsch, S., Reid, S.P., Martins, K., Encinales, L., Pacheco, N., Amdur, R.L., Porras-Ramirez, A., Rico-Mendoza, A., Li, G., et al. (2018). The Cytokine Profile in Acute Chikungunya Infection is Predictive of Chronic Arthritis 20 Months Post Infection. Diseases 6. 10.3390/diseases6040095.

17. Liu, X., Poo, Y.S., Alves, J.C., Almeida, R.P., Mostafavi, H., Tang, P.C.H., Bucala, R., Teixeira, M.M., Taylor, A., Zaid, A., and Mahalingam, S. (2022). Interleukin-17 Contributes to Chikungunya Virus-Induced Disease. mBio, e0028922. 10.1128/mbio.00289-22.

18. Schilte, C., Staikowsky, F., Couderc, T., Madec, Y., Carpentier, F., Kassab, S., Albert, M.L., Lecuit, M., and Michault, A. (2013). Chikungunya virus-associated long-term arthralgia: a 36-month prospective longitudinal study. PLoS Negl Trop Dis 7, e2137. 10.1371/journal.pntd.0002137.

19. Manimunda, S.P., Vijayachari, P., Uppoor, R., Sugunan, A.P., Singh, S.S., Rai, S.K., Sudeep, A.B., Muruganandam, N., Chaitanya, I.K., and Guruprasad, D.R. (2010). Clinical progression of chikungunya fever during acute and chronic arthritic stages and the changes in joint morphology as revealed by imaging. Trans R Soc Trop Med Hyg 104, 392–399. 10.1016/j.trstmh.2010.01.011.

20. Amaral, J.K., Taylor, P.C., and Schoen, R.T. (2024). Bone erosions and joint damage caused by chikungunya virus: a systematic review. Rev Soc Bras Med Trop 57, e00404. 10.1590/0037-8682-0433-2023.

21. Bouquillard, E., and Combe, B. (2009). A report of 21 cases of rheumatoid arthritis following Chikungunya fever. A mean follow-up of two years. Joint Bone Spine 76, 654–657. 10.1016/j.jbspin.2009.08.005.

22. Gardner, J., Anraku, I., Le, T.T., Larcher, T., Major, L., Roques, P., Schroder, W.A., Higgs, S., and Suhrbier, A. (2010). Chikungunya virus arthritis in adult wild-type mice. J Virol 84, 8021–8032. 10.1128/JVI.02603-09.

23. Morrison, T.E., Oko, L., Montgomery, S.A., Whitmore, A.C., Lotstein, A.R., Gunn, B.M., Elmore, S.A., and Heise, M.T. (2011). A mouse model of chikungunya virus-induced musculoskeletal inflammatory disease: evidence of arthritis, tenosynovitis, myositis, and persistence. Am J Pathol 178, 32–40. 10.1016/j.ajpath.2010.11.018.

24. Ziegler, S.A., Lu, L., da Rosa, A.P., Xiao, S.Y., and Tesh, R.B. (2008). An animal model for studying the pathogenesis of chikungunya virus infection. Am J Trop Med Hyg 79, 133–139.

25. Seymour, R.L., Adams, A.P., Leal, G., Alcorn, M.D., and Weaver, S.C. (2015). A Rodent Model of Chikungunya Virus Infection in RAG1 -/- Mice, with Features of Persistence, for Vaccine Safety Evaluation. PLoS Negl Trop Dis 9, e0003800. 10.1371/journal.pntd.0003800.

26. Teo, T.H., Lum, F.M., Claser, C., Lulla, V., Lulla, A., Merits, A., Renia, L., and Ng, L.F. (2013). A pathogenic role for CD4+ T cells during Chikungunya virus infection in mice. J Immunol 190, 259–269. 10.4049/jimmunol.1202177[pii].

27. Teo, T.H., Chan, Y.H., Lee, W.W., Lum, F.M., Amrun, S.N., Her, Z., Rajarethinam, R., Merits, A., Rotzschke, O., Renia, L., and Ng, L.F. (2017). Fingolimod treatment abrogates chikungunya virus-induced arthralgia. Sci Transl Med 9. 10.1126/scitranslmed.aal1333.

28. Gruneboom, A., Hawwari, I., Weidner, D., Culemann, S., Muller, S., Henneberg, S., Brenzel, A., Merz, S., Bornemann, L., Zec, K., et al. (2019). A network of trans-cortical capillaries as mainstay for blood circulation in long bones. Nat Metab 1, 236–250. 10.1038/s42255-018-0016-5.

29. Werner, D., Simon, D., Englbrecht, M., Stemmler, F., Simon, C., Berlin, A., Haschka, J., Renner, N., Buder, T., Engelke, K., et al. (2017). Early Changes of the Cortical Micro-Channel System in the Bare Area of the Joints of Patients With Rheumatoid Arthritis. Arthritis Rheumatol 69, 1580–1587. 10.1002/art.40148.

30. Binks, D.A., Gravallese, E.M., Bergin, D., Hodgson, R.J., Tan, A.L., Matzelle, M.M., McGonagle, D., and Radjenovic, A. (2015). Role of vascular channels as a novel mechanism for subchondral bone damage at cruciate ligament entheses in osteoarthritis and inflammatory arthritis. Ann Rheum Dis 74, 196–203. 10.1136/annrheumdis-2013-203972.

31. Marinova-Mutafchieva, L., Williams, R.O., Funa, K., Maini, R.N., and Zvaifler, N.J. (2002). Inflammation is preceded by tumor necrosis factor-dependent infiltration of mesenchymal cells in experimental arthritis. Arthritis Rheum 46, 507–513. 10.1002/art.10126.

32. Winkler, E.S., Shrihari, S., Hykes, B.L., Jr., Handley, S.A., Andhey, P.S., Huang, Y.S., Swain, A., Droit, L., Chebrolu, K.K., Mack, M., et al. (2020). The Intestinal Microbiome Restricts Alphavirus Infection and Dissemination through a Bile Acid-Type I IFN Signaling Axis. Cell 182, 901–918 e918. 10.1016/j.cell.2020.06.029.

33. Wexler, A.G., and Goodman, A.L. (2017). An insider’s perspective: Bacteroides as a window into the microbiome. Nat Microbiol 2, 17026. 10.1038/nmicrobiol.2017.26.

34. Jordan, C.K.I., Brown, R.L., Larkinson, M.L.Y., Sequeira, R.P., Edwards, A.M., and Clarke, T.B. (2023). Symbiotic Firmicutes establish mutualism with the host via innate tolerance and resistance to control systemic immunity. Cell Host Microbe 31, 1433–1449 e1439. 10.1016/j.chom.2023.07.008.

35. Sharon, G., Garg, N., Debelius, J., Knight, R., Dorrestein, P.C., and Mazmanian, S.K. (2014). Specialized metabolites from the microbiome in health and disease. Cell Metab 20, 719–730. 10.1016/j.cmet.2014.10.016.

36. Ghosh, S., Whitley, C.S., Haribabu, B., and Jala, V.R. (2021). Regulation of Intestinal Barrier Function by Microbial Metabolites. Cell Mol Gastroenterol Hepatol 11, 1463–1482. 10.1016/j.jcmgh.2021.02.007.

37. Gasaly, N., de Vos, P., and Hermoso, M.A. (2021). Impact of Bacterial Metabolites on Gut Barrier Function and Host Immunity: A Focus on Bacterial Metabolism and Its Relevance for Intestinal Inflammation. Front Immunol 12, 658354. 10.3389/fimmu.2021.658354.

38. Zheng, D., Liwinski, T., and Elinav, E. (2020). Interaction between microbiota and immunity in health and disease. Cell Res 30, 492–506. 10.1038/s41422-020-0332-7.

39. Ridlon, J.M., Kang, D.J., Hylemon, P.B., and Bajaj, J.S. (2014). Bile acids and the gut microbiome. Curr Opin Gastroenterol 30, 332–338. 10.1097/MOG.0000000000000057.

40. van der Hee, B., and Wells, J.M. (2021). Microbial Regulation of Host Physiology by Short-chain Fatty Acids. Trends Microbiol 29, 700–712. 10.1016/j.tim.2021.02.001.

41. Stenman, L.K., Holma, R., Eggert, A., and Korpela, R. (2013). A novel mechanism for gut barrier dysfunction by dietary fat: epithelial disruption by hydrophobic bile acids. Am J Physiol Gastrointest Liver Physiol 304, G227–234. 10.1152/ajpgi.00267.2012.

42. Raimondi, F., Santoro, P., Barone, M.V., Pappacoda, S., Barretta, M.L., Nanayakkara, M., Apicella, C., Capasso, L., and Paludetto, R. (2008). Bile acids modulate tight junction structure and barrier function of Caco-2 monolayers via EGFR activation. Am J Physiol Gastrointest Liver Physiol 294, G906–913. 10.1152/ajpgi.00043.2007.

43. Hughes, R., Kurth, M.J., McGilligan, V., McGlynn, H., and Rowland, I. (2008). Effect of colonic bacterial metabolites on Caco-2 cell paracellular permeability in vitro. Nutr Cancer 60, 259–266. 10.1080/01635580701649644.

44. Liu, L., Dong, W., Wang, S., Zhang, Y., Liu, T., Xie, R., Wang, B., and Cao, H. (2018). Deoxycholic acid disrupts the intestinal mucosal barrier and promotes intestinal tumorigenesis. Food Funct 9, 5588–5597. 10.1039/c8fo01143e.

45. Perez-Reytor, D., Puebla, C., Karahanian, E., and Garcia, K. (2021). Use of Short-Chain Fatty Acids for the Recovery of the Intestinal Epithelial Barrier Affected by Bacterial Toxins. Front Physiol 12, 650313. 10.3389/fphys.2021.650313.

46. Furusawa, Y., Obata, Y., Fukuda, S., Endo, T.A., Nakato, G., Takahashi, D., Nakanishi, Y., Uetake, C., Kato, K., Kato, T., et al. (2013). Commensal microbe-derived butyrate induces the differentiation of colonic regulatory T cells. Nature 504, 446–450. 10.1038/nature12721.

47. Smith, P.M., Howitt, M.R., Panikov, N., Michaud, M., Gallini, C.A., Bohlooly, Y.M., Glickman, J.N., and Garrett, W.S. (2013). The microbial metabolites, short-chain fatty acids, regulate colonic Treg cell homeostasis. Science 341, 569–573. 10.1126/science.1241165.

48. Arpaia, N., Campbell, C., Fan, X., Dikiy, S., van der Veeken, J., deRoos, P., Liu, H., Cross, J.R., Pfeffer, K., Coffer, P.J., and Rudensky, A.Y. (2013). Metabolites produced by commensal bacteria promote peripheral regulatory T-cell generation. Nature 504, 451–455. 10.1038/nature12726.

49. Prow, N.A., Hirata, T.D.C., Tang, B., Larcher, T., Mukhopadhyay, P., Alves, T.L., Le, T.T., Gardner, J., Poo, Y.S., Nakayama, E., et al. (2019). Exacerbation of Chikungunya Virus Rheumatic Immunopathology by a High Fiber Diet and Butyrate. Front Immunol 10, 2736. 10.3389/fimmu.2019.02736.

50. Koropatkin, N.M., Martens, E.C., Gordon, J.I., and Smith, T.J. (2008). Starch catabolism by a prominent human gut symbiont is directed by the recognition of amylose helices. Structure 16, 1105–1115. 10.1016/j.str.2008.03.017.

51. Han, S., Van Treuren, W., Fischer, C.R., Merrill, B.D., DeFelice, B.C., Sanchez, J.M., Higginbottom, S.K., Guthrie, L., Fall, L.A., Dodd, D., et al. (2021). A metabolomics pipeline for the mechanistic interrogation of the gut microbiome. Nature 595, 415–420. 10.1038/s41586-021-03707-9.

52. Balka, K.R., and De Nardo, D. (2019). Understanding early TLR signaling through the Myddosome. J Leukoc Biol 105, 339–351. 10.1002/JLB.MR0318-096R.

53. Friedrich, C., Mamareli, P., Thiemann, S., Kruse, F., Wang, Z., Holzmann, B., Strowig, T., Sparwasser, T., and Lochner, M. (2017). MyD88 signaling in dendritic cells and the intestinal epithelium controls immunity against intestinal infection with C. rodentium. PLoS Pathog 13, e1006357. 10.1371/journal.ppat.1006357.

54. Frantz, A.L., Rogier, E.W., Weber, C.R., Shen, L., Cohen, D.A., Fenton, L.A., Bruno, M.E., and Kaetzel, C.S. (2012). Targeted deletion of MyD88 in intestinal epithelial cells results in compromised antibacterial immunity associated with downregulation of polymeric immunoglobulin receptor, mucin-2, and antibacterial peptides. Mucosal Immunol 5, 501–512. 10.1038/mi.2012.23.

55. Mamareli, P., Kruse, F., Friedrich, C., Smit, N., Strowig, T., Sparwasser, T., and Lochner, M. (2019). Epithelium-specific MyD88 signaling, but not DCs or macrophages, control acute intestinal infection with Clostridium difficile. Eur J Immunol 49, 747–757. 10.1002/eji.201848022.

56. Bhinder, G., Stahl, M., Sham, H.P., Crowley, S.M., Morampudi, V., Dalwadi, U., Ma, C., Jacobson, K., and Vallance, B.A. (2014). Intestinal epithelium-specific MyD88 signaling impacts host susceptibility to infectious colitis by promoting protective goblet cell and antimicrobial responses. Infect Immun 82, 3753–3763. 10.1128/IAI.02045-14.

57. Gong, J., Xu, J., Zhu, W., Gao, X., Li, N., and Li, J. (2010). Epithelial-specific blockade of MyD88-dependent pathway causes spontaneous small intestinal inflammation. Clin Immunol 136, 245–256. 10.1016/j.clim.2010.04.001.

58. Everard, A., Geurts, L., Caesar, R., Van Hul, M., Matamoros, S., Duparc, T., Denis, R.G., Cochez, P., Pierard, F., Castel, J., et al. (2014). Intestinal epithelial MyD88 is a sensor switching host metabolism towards obesity according to nutritional status. Nat Commun 5, 5648. 10.1038/ncomms6648.

59. Watanabe, I.K.M., Andrade-Silva, M., Foresto-Neto, O., Felizardo, R.J.F., Matheus, M.A.C., Silva, R.C., Cenedeze, M.A., Honda, T.S.B., Perandini, L.A.B., Volpini, R.A., et al. (2020). Gut Microbiota and Intestinal Epithelial Myd88 Signaling Are Crucial for Renal Injury in UUO Mice. Front Immunol 11, 578623. 10.3389/fimmu.2020.578623.

60. Gerbe, F., Sidot, E., Smyth, D.J., Ohmoto, M., Matsumoto, I., Dardalhon, V., Cesses, P., Garnier, L., Pouzolles, M., Brulin, B., et al. (2016). Intestinal epithelial tuft cells initiate type 2 mucosal immunity to helminth parasites. Nature 529, 226–230. 10.1038/nature16527.

61. Yamashita, J., Ohmoto, M., Yamaguchi, T., Matsumoto, I., and Hirota, J. (2017). Skn-1a/Pou2f3 functions as a master regulator to generate Trpm5-expressing chemosensory cells in mice. PLoS One 12, e0189340. 10.1371/journal.pone.0189340.

62. Poo, Y.S., Nakaya, H., Gardner, J., Larcher, T., Schroder, W.A., Le, T.T., Major, L.D., and Suhrbier, A. (2014). CCR2 deficiency promotes exacerbated chronic erosive neutrophil-dominated chikungunya virus arthritis. J Virol 88, 6862–6872. 10.1128/JVI.03364-13.

63. Barnden, M.J., Allison, J., Heath, W.R., and Carbone, F.R. (1998). Defective TCR expression in transgenic mice constructed using cDNA-based alpha- and beta-chain genes under the control of heterologous regulatory elements. Immunol Cell Biol 76, 34–40. 10.1046/j.1440-1711.1998.00709.x.

64. Shinkai, Y., Koyasu, S., Nakayama, K., Murphy, K.M., Loh, D.Y., Reinherz, E.L., and Alt, F.W. (1993). Restoration of T cell development in RAG-2-deficient mice by functional TCR transgenes. Science 259, 822–825. 10.1126/science.8430336.

65. Thackray, L.B., Handley, S.A., Gorman, M.J., Poddar, S., Bagadia, P., Briseno, C.G., Theisen, D.J., Tan, Q., Hykes, B.L., Jr., Lin, H., et al. (2018). Oral Antibiotic Treatment of Mice Exacerbates the Disease Severity of Multiple Flavivirus Infections. Cell Rep 22, 3440–3453 e3446. 10.1016/j.celrep.2018.03.001.

66. Brown, E.M., Kenny, D.J., and Xavier, R.J. (2019). Gut Microbiota Regulation of T Cells During Inflammation and Autoimmunity. Annu Rev Immunol 37, 599–624. 10.1146/annurev-immunol-042718-041841.

67. Yu, A.I., Zhao, L., Eaton, K.A., Ho, S., Chen, J., Poe, S., Becker, J., Gonzalez, A., McKinstry, D., Hasso, M., et al. (2020). Gut Microbiota Modulate CD8 T Cell Responses to Influence Colitis-Associated Tumorigenesis. Cell Rep 31, 107471. 10.1016/j.celrep.2020.03.035.

68. Tanoue, T., Morita, S., Plichta, D.R., Skelly, A.N., Suda, W., Sugiura, Y., Narushima, S., Vlamakis, H., Motoo, I., Sugita, K., et al. (2019). A defined commensal consortium elicits CD8 T cells and anti-cancer immunity. Nature 565, 600–605. 10.1038/s41586-019-0878-z.

69. Park, J., Kim, M., Kang, S.G., Jannasch, A.H., Cooper, B., Patterson, J., and Kim, C.H. (2015). Short-chain fatty acids induce both effector and regulatory T cells by suppression of histone deacetylases and regulation of the mTOR-S6K pathway. Mucosal Immunol 8, 80–93. 10.1038/mi.2014.44.

70. Lefferts, A.R., Norman, E., Claypool, D.J., Kantheti, U., and Kuhn, K.A. (2022). Cytokine competent gut-joint migratory T Cells contribute to inflammation in the joint. Front Immunol 13, 932393. 10.3389/fimmu.2022.932393.

71. Ogawa, H., Binion, D.G., Heidemann, J., Theriot, M., Fisher, P.J., Johnson, N.A., Otterson, M.F., and Rafiee, P. (2005). Mechanisms of MAdCAM-1 gene expression in human intestinal microvascular endothelial cells. Am J Physiol Cell Physiol 288, C272–281. 10.1152/ajpcell.00406.2003.

72. Sikorski, E.E., Hallmann, R., Berg, E.L., and Butcher, E.C. (1993). The Peyer’s patch high endothelial receptor for lymphocytes, the mucosal vascular addressin, is induced on a murine endothelial cell line by tumor necrosis factor-alpha and IL-1. J Immunol 151, 5239–5250.

73. Oshima, T., Jordan, P., Grisham, M.B., Alexander, J.S., Jennings, M., Sasaki, M., and Manas, K. (2001). TNF-alpha induced endothelial MAdCAM-1 expression is regulated by exogenous, not endogenous nitric oxide. BMC Gastroenterol 1, 5. 10.1186/1471-230x-1-5.

74. Takeuchi, M., and Baichwal, V.R. (1995). Induction of the gene encoding mucosal vascular addressin cell adhesion molecule 1 by tumor necrosis factor alpha is mediated by NF-kappa B proteins. Proc Natl Acad Sci U S A 92, 3561–3565. 10.1073/pnas.92.8.3561.

75. Berlin, C., Berg, E.L., Briskin, M.J., Andrew, D.P., Kilshaw, P.J., Holzmann, B., Weissman, I.L., Hamann, A., and Butcher, E.C. (1993). Alpha 4 beta 7 integrin mediates lymphocyte binding to the mucosal vascular addressin MAdCAM-1. Cell 74, 185–195. 10.1016/0092-8674(93)90305-a.

76. Farstad, I.N., Halstensen, T.S., Kvale, D., Fausa, O., and Brandtzaeg, P. (1997). Topographic distribution of homing receptors on B and T cells in human gut-associated lymphoid tissue: relation of L-selectin and integrin alpha 4 beta 7 to naive and memory phenotypes. Am J Pathol 150, 187–199.

77. Briskin, M., Winsor-Hines, D., Shyjan, A., Cochran, N., Bloom, S., Wilson, J., McEvoy, L.M., Butcher, E.C., Kassam, N., Mackay, C.R., et al. (1997). Human mucosal addressin cell adhesion molecule-1 is preferentially expressed in intestinal tract and associated lymphoid tissue. Am J Pathol 151, 97–110.

78. Streeter, P.R., Berg, E.L., Rouse, B.T., Bargatze, R.F., and Butcher, E.C. (1988). A tissue-specific endothelial cell molecule involved in lymphocyte homing. Nature 331, 41–46. 10.1038/331041a0.

79. Nakache, M., Berg, E.L., Streeter, P.R., and Butcher, E.C. (1989). The mucosal vascular addressin is a tissue-specific endothelial cell adhesion molecule for circulating lymphocytes. Nature 337, 179–181. 10.1038/337179a0.

80. Holzmann, B., McIntyre, B.W., and Weissman, I.L. (1989). Identification of a murine Peyer’s patch--specific lymphocyte homing receptor as an integrin molecule with an alpha chain homologous to human VLA-4 alpha. Cell 56, 37–46. 10.1016/0092-8674(89)90981-1.

81. Chen, W., Foo, S.S., Rulli, N.E., Taylor, A., Sheng, K.C., Herrero, L.J., Herring, B.L., Lidbury, B.A., Li, R.W., Walsh, N.C., et al. (2014). Arthritogenic alphaviral infection perturbs osteoblast function and triggers pathologic bone loss. Proc Natl Acad Sci U S A 111, 6040–6045. 10.1073/pnas.1318859111.

82. Chen, W., Foo, S.S., Sims, N.A., Herrero, L.J., Walsh, N.C., and Mahalingam, S. (2015). Arthritogenic alphaviruses: new insights into arthritis and bone pathology. Trends Microbiol 23, 35–43. 10.1016/j.tim.2014.09.005.

83. Chen, W., Foo, S.S., Taylor, A., Lulla, A., Merits, A., Hueston, L., Forwood, M.R., Walsh, N.C., Sims, N.A., Herrero, L.J., and Mahalingam, S. (2015). Bindarit, an inhibitor of monocyte chemotactic protein synthesis, protects against bone loss induced by chikungunya virus infection. J Virol 89, 581–593. 10.1128/JVI.02034-14.

84. Wolf, S., Taylor, A., Zaid, A., Freitas, J., Herrero, L.J., Rao, S., Suhrbier, A., Forwood, M.R., Bucala, R., and Mahalingam, S. (2019). Inhibition of Interleukin-1beta Signaling by Anakinra Demonstrates a Critical Role of Bone Loss in Experimental Arthritogenic Alphavirus Infections. Arthritis Rheumatol 71, 1185–1190. 10.1002/art.40856.

85. Lu, W., Zheng, C., Zhang, H., Cheng, P., Miao, S., Wang, H., He, T., Fan, J., Hu, Y., Liu, H., et al. (2023). Hedgehog signaling regulates bone homeostasis through orchestrating osteoclast differentiation and osteoclast-osteoblast coupling. Cell Mol Life Sci 80, 171. 10.1007/s00018-023-04821-9.

86. Kohara, Y., Haraguchi, R., Kitazawa, R., Imai, Y., and Kitazawa, S. (2020). Hedgehog Inhibitors Suppress Osteoclastogenesis in In Vitro Cultures, and Deletion of Smo in Macrophage/Osteoclast Lineage Prevents Age-Related Bone Loss. Int J Mol Sci 21. 10.3390/ijms21082745.

87. Liang, M., Yin, X., Zhang, S., Ai, H., Luo, F., Xu, J., Dou, C., Dong, S., and Ma, Q. (2021). Osteoclast-derived small extracellular vesicles induce osteogenic differentiation via inhibiting ARHGAP1. Mol Ther Nucleic Acids 23, 1191–1203. 10.1016/j.omtn.2021.01.031.

88. Ikebuchi, Y., Aoki, S., Honma, M., Hayashi, M., Sugamori, Y., Khan, M., Kariya, Y., Kato, G., Tabata, Y., Penninger, J.M., et al. (2018). Coupling of bone resorption and formation by RANKL reverse signalling. Nature 561, 195–200. 10.1038/s41586-018-0482-7.

89. Fujita, H., Ochi, M., Ono, M., Aoyama, E., Ogino, T., Kondo, Y., and Ohuchi, H. (2019). Glutathione accelerates osteoclast differentiation and inflammatory bone destruction. Free Radic Res 53, 226–236. 10.1080/10715762.2018.1563782.

90. Han, B., Geng, H., Liu, L., Wu, Z., and Wang, Y. (2020). GSH attenuates RANKL-induced osteoclast formation in vitro and LPS-induced bone loss in vivo. Biomed Pharmacother 128, 110305. 10.1016/j.biopha.2020.110305.

91. Ha, H., Kwak, H.B., Lee, S.W., Jin, H.M., Kim, H.M., Kim, H.H., and Lee, Z.H. (2004). Reactive oxygen species mediate RANK signaling in osteoclasts. Exp Cell Res 301, 119–127. 10.1016/j.yexcr.2004.07.035.

92. Lean, J.M., Davies, J.T., Fuller, K., Jagger, C.J., Kirstein, B., Partington, G.A., Urry, Z.L., and Chambers, T.J. (2003). A crucial role for thiol antioxidants in estrogen-deficiency bone loss. J Clin Invest 112, 915–923. 10.1172/JCI18859.

93. Huh, Y.J., Kim, J.M., Kim, H., Song, H., So, H., Lee, S.Y., Kwon, S.B., Kim, H.J., Kim, H.H., Lee, S.H., et al. (2006). Regulation of osteoclast differentiation by the redox-dependent modulation of nuclear import of transcription factors. Cell Death Differ 13, 1138–1146. 10.1038/sj.cdd.4401793.

94. Kim, H., Kim, I.Y., Lee, S.Y., and Jeong, D. (2006). Bimodal actions of reactive oxygen species in the differentiation and bone-resorbing functions of osteoclasts. FEBS Lett 580, 5661–5665. 10.1016/j.febslet.2006.09.015.

95. Aita, T., Yamamura, M., Kawashima, M., Okamoto, A., Iwahashi, M., Yamana, J., and Makino, H. (2004). Expression of interleukin 12 receptor (IL-12R) and IL-18R on CD4+ T cells from patients with rheumatoid arthritis. J Rheumatol 31, 448–456.

96. Nakanishi, K., Yoshimoto, T., Tsutsui, H., and Okamura, H. (2001). Interleukin-18 regulates both Th1 and Th2 responses. Annu Rev Immunol 19, 423–474. 10.1146/annurev.immunol.19.1.423.

97. Tu, J., Huang, W., Zhang, W., Mei, J., and Zhu, C. (2021). A Tale of Two Immune Cells in Rheumatoid Arthritis: The Crosstalk Between Macrophages and T Cells in the Synovium. Front Immunol 12, 655477. 10.3389/fimmu.2021.655477.

98. Koduru, S.V., Sun, B.H., Walker, J.M., Zhu, M., Simpson, C., Dhodapkar, M., and Insogna, K.L. (2018). The contribution of cross-talk between the cell-surface proteins CD36 and CD47-TSP-1 in osteoclast formation and function. J Biol Chem 293, 15055–15069. 10.1074/jbc.RA117.000633.

99. Timmen, M., Hidding, H., Gotte, M., Khassawna, T.E., Kronenberg, D., and Stange, R. (2020). The heparan sulfate proteoglycan Syndecan-1 influences local bone cell communication via the RANKL/OPG axis. Sci Rep 10, 20510. 10.1038/s41598-020-77510-3.

100. Gu, R., Santos, L.L., Ngo, D., Fan, H., Singh, P.P., Fingerle-Rowson, G., Bucala, R., Xu, J., Quinn, J.M., and Morand, E.F. (2015). Macrophage migration inhibitory factor is essential for osteoclastogenic mechanisms in vitro and in vivo mouse model of arthritis. Cytokine 72, 135–145. 10.1016/j.cyto.2014.11.015.

101. Movila, A., Ishii, T., Albassam, A., Wisitrasameewong, W., Howait, M., Yamaguchi, T., Ruiz-Torruella, M., Bahammam, L., Nishimura, K., Van Dyke, T., and Kawai, T. (2016). Macrophage Migration Inhibitory Factor (MIF) Supports Homing of Osteoclast Precursors to Peripheral Osteolytic Lesions. J Bone Miner Res 31, 1688–1700. 10.1002/jbmr.2854.

102. Ochi, S., Shinohara, M., Sato, K., Gober, H.J., Koga, T., Kodama, T., Takai, T., Miyasaka, N., and Takayanagi, H. (2007). Pathological role of osteoclast costimulation in arthritis-induced bone loss. Proc Natl Acad Sci U S A 104, 11394–11399. 10.1073/pnas.0701971104.

103. Koga, T., Inui, M., Inoue, K., Kim, S., Suematsu, A., Kobayashi, E., Iwata, T., Ohnishi, H., Matozaki, T., Kodama, T., et al. (2004). Costimulatory signals mediated by the ITAM motif cooperate with RANKL for bone homeostasis. Nature 428, 758–763. 10.1038/nature02444.

104. Grau, K.R., Zhu, S., Peterson, S.T., Helm, E.W., Philip, D., Phillips, M., Hernandez, A., Turula, H., Frasse, P., Graziano, V.R., et al. (2020). The intestinal regionalization of acute norovirus infection is regulated by the microbiota via bile acid-mediated priming of type III interferon. Nat Microbiol 5, 84–92. 10.1038/s41564-019-0602-7.

105. Van Winkle, J.A., Peterson, S.T., Kennedy, E.A., Wheadon, M.J., Ingle, H., Desai, C., Rodgers, R., Constant, D.A., Wright, A.P., Li, L., et al. (2022). Homeostatic interferon-lambda response to bacterial microbiota stimulates preemptive antiviral defense within discrete pockets of intestinal epithelium. Elife 11. 10.7554/eLife.74072.

106. Jacques, P., and Elewaut, D. (2008). Joint expedition: linking gut inflammation to arthritis. Mucosal Immunol 1, 364–371. 10.1038/mi.2008.24.

107. Ashrafi, M., Kuhn, K.A., and Weisman, M.H. (2021). The arthritis connection to inflammatory bowel disease (IBD): why has it taken so long to understand it? RMD Open 7. 10.1136/rmdopen-2020-001558.

108. Lin, L., Zhang, K., Xiong, Q., Zhang, J., Cai, B., Huang, Z., Yang, B., Wei, B., Chen, J., and Niu, Q. (2023). Gut microbiota in pre-clinical rheumatoid arthritis: From pathogenesis to preventing progression. J Autoimmun 141, 103001. 10.1016/j.jaut.2023.103001.

109. Wang, Y., Wei, J., Zhang, W., Doherty, M., Zhang, Y., Xie, H., Li, W., Wang, N., Lei, G., and Zeng, C. (2022). Gut dysbiosis in rheumatic diseases: A systematic review and meta-analysis of 92 observational studies. EBioMedicine 80, 104055. 10.1016/j.ebiom.2022.104055.

110. He, J., Chu, Y., Li, J., Meng, Q., Liu, Y., Jin, J., Wang, Y., Wang, J., Huang, B., Shi, L., et al. (2022). Intestinal butyrate-metabolizing species contribute to autoantibody production and bone erosion in rheumatoid arthritis. Sci Adv 8, eabm1511. 10.1126/sciadv.abm1511.

111. Andreasson, K., Olofsson, T., Lagishetty, V., Alrawi, Z., Klaassens, E., Holster, S., Hesselstrand, R., Jacobs, J.P., Wallman, J.K., and Volkmann, E.R. (2024). Treatment for Rheumatoid Arthritis Associated With Alterations in the Gastrointestinal Microbiota. ACR Open Rheumatol 6, 421–427. 10.1002/acr2.11673.

112. Picchianti-Diamanti, A., Panebianco, C., Salemi, S., Sorgi, M.L., Di Rosa, R., Tropea, A., Sgrulletti, M., Salerno, G., Terracciano, F., D’Amelio, R., et al. (2018). Analysis of Gut Microbiota in Rheumatoid Arthritis Patients: Disease-Related Dysbiosis and Modifications Induced by Etanercept. Int J Mol Sci 19. 10.3390/ijms19102938.

113. Zhang, X., Zhang, D., Jia, H., Feng, Q., Wang, D., Liang, D., Wu, X., Li, J., Tang, L., Li, Y., et al. (2015). The oral and gut microbiomes are perturbed in rheumatoid arthritis and partly normalized after treatment. Nat Med 21, 895–905. 10.1038/nm.3914.

114. Chen, H., Shi, J., Tang, C., Xu, J., Li, B., Wang, J., Zhou, Y., Yang, Y., Yang, H., Huang, Q., et al. (2024). CHIKV infection drives shifts in the gastrointestinal microbiome and metabolites in rhesus monkeys. Microbiome 12, 161. 10.1186/s40168-024-01895-w.

115. Rawle, D.J., Dumenil, T., Tang, B., Bishop, C.R., Yan, K., Le, T.T., and Suhrbier, A. (2022). Microplastic consumption induces inflammatory signatures in the colon and prolongs a viral arthritis. Sci Total Environ 809, 152212. 10.1016/j.scitotenv.2021.152212.

116. Haist, K.C., Burrack, K.S., Davenport, B.J., and Morrison, T.E. (2017). Inflammatory monocytes mediate control of acute alphavirus infection in mice. PLoS Pathog 13, e1006748. 10.1371/journal.ppat.1006748.

117. Rulli, N.E., Rolph, M.S., Srikiatkhachorn, A., Anantapreecha, S., Guglielmotti, A., and Mahalingam, S. (2011). Protection from arthritis and myositis in a mouse model of acute chikungunya virus disease by bindarit, an inhibitor of monocyte chemotactic protein-1 synthesis. J Infect Dis 204, 1026–1030. 10.1093/infdis/jir470.

118. Lum, F.M., Chan, Y.H., Teo, T.H., Becht, E., Amrun, S.N., Teng, K.W., Hartimath, S.V., Yeo, N.K., Yee, W.X., Ang, N., et al. (2024). Crosstalk between CD64(+)MHCII(+) macrophages and CD4(+) T cells drives joint pathology during chikungunya. EMBO Mol Med 16, 641–663. 10.1038/s44321-024-00028-y.

119. Muhlbauer, M., Perez-Chanona, E., and Jobin, C. (2013). Epithelial cell-specific MyD88 signaling mediates ischemia/reperfusion-induced intestinal injury independent of microbial status. Inflamm Bowel Dis 19, 2857–2866. 10.1097/01.MIB.0000435445.96933.37.

120. Tajik, N., Frech, M., Schulz, O., Schalter, F., Lucas, S., Azizov, V., Durholz, K., Steffen, F., Omata, Y., Rings, A., et al. (2020). Targeting zonulin and intestinal epithelial barrier function to prevent onset of arthritis. Nat Commun 11, 1995. 10.1038/s41467-020-15831-7.

121. Matei, D.E., Menon, M., Alber, D.G., Smith, A.M., Nedjat-Shokouhi, B., Fasano, A., Magill, L., Duhlin, A., Bitoun, S., Gleizes, A., et al. (2021). Intestinal barrier dysfunction plays an integral role in arthritis pathology and can be targeted to ameliorate disease. Med (N Y) 2, 864–883 e869. 10.1016/j.medj.2021.04.013.

122. Lucas, S., Omata, Y., Hofmann, J., Bottcher, M., Iljazovic, A., Sarter, K., Albrecht, O., Schulz, O., Krishnacoumar, B., Kronke, G., et al. (2018). Short-chain fatty acids regulate systemic bone mass and protect from pathological bone loss. Nat Commun 9, 55. 10.1038/s41467-017-02490-4.

123. Trompette, A., Gollwitzer, E.S., Yadava, K., Sichelstiel, A.K., Sprenger, N., Ngom-Bru, C., Blanchard, C., Junt, T., Nicod, L.P., Harris, N.L., and Marsland, B.J. (2014). Gut microbiota metabolism of dietary fiber influences allergic airway disease and hematopoiesis. Nat Med 20, 159–166. 10.1038/nm.3444.

124. Zaiss, M.M., Rapin, A., Lebon, L., Dubey, L.K., Mosconi, I., Sarter, K., Piersigilli, A., Menin, L., Walker, A.W., Rougemont, J., et al. (2015). The Intestinal Microbiota Contributes to the Ability of Helminths to Modulate Allergic Inflammation. Immunity 43, 998–1010. 10.1016/j.immuni.2015.09.012.

125. Platt, D.J., Lawrence, D., Rodgers, R., Schriefer, L., Qian, W., Miner, C.A., Menos, A.M., Kennedy, E.A., Peterson, S.T., Stinson, W.A., et al. (2021). Transferrable protection by gut microbes against STING-associated lung disease. Cell Rep 35, 109113. 10.1016/j.celrep.2021.109113.

126. Haghikia, A., Jorg, S., Duscha, A., Berg, J., Manzel, A., Waschbisch, A., Hammer, A., Lee, D.H., May, C., Wilck, N., et al. (2015). Dietary Fatty Acids Directly Impact Central Nervous System Autoimmunity via the Small Intestine. Immunity 43, 817–829. 10.1016/j.immuni.2015.09.007.

127. Thorburn, A.N., McKenzie, C.I., Shen, S., Stanley, D., Macia, L., Mason, L.J., Roberts, L.K., Wong, C.H., Shim, R., Robert, R., et al. (2015). Evidence that asthma is a developmental origin disease influenced by maternal diet and bacterial metabolites. Nat Commun 6, 7320. 10.1038/ncomms8320.

128. Gasaly, N., Hermoso, M.A., and Gotteland, M. (2021). Butyrate and the Fine-Tuning of Colonic Homeostasis: Implication for Inflammatory Bowel Diseases. Int J Mol Sci 22. 10.3390/ijms22063061.

129. Donohoe, D.R., Collins, L.B., Wali, A., Bigler, R., Sun, W., and Bultman, S.J. (2012). The Warburg effect dictates the mechanism of butyrate-mediated histone acetylation and cell proliferation. Mol Cell 48, 612–626. 10.1016/j.molcel.2012.08.033.

130. Christl, S.U., Eisner, H.D., Dusel, G., Kasper, H., and Scheppach, W. (1996). Antagonistic effects of sulfide and butyrate on proliferation of colonic mucosa: a potential role for these agents in the pathogenesis of ulcerative colitis. Dig Dis Sci 41, 2477–2481. 10.1007/BF02100146.

131. Liu, J., Zhu, H., Li, B., Lee, C., Alganabi, M., Zheng, S., and Pierro, A. (2020). Beneficial effects of butyrate in intestinal injury. J Pediatr Surg 55, 1088–1093. 10.1016/j.jpedsurg.2020.02.036.

132. Peng, L., He, Z., Chen, W., Holzman, I.R., and Lin, J. (2007). Effects of butyrate on intestinal barrier function in a Caco-2 cell monolayer model of intestinal barrier. Pediatr Res 61, 37–41. 10.1203/01.pdr.0000250014.92242.f3.

133. Peng, L., Li, Z.R., Green, R.S., Holzman, I.R., and Lin, J. (2009). Butyrate enhances the intestinal barrier by facilitating tight junction assembly via activation of AMP-activated protein kinase in Caco-2 cell monolayers. J Nutr 139, 1619–1625. 10.3945/jn.109.104638.

134. Kurowska-Stolarska, M., and Alivernini, S. (2022). Synovial tissue macrophages in joint homeostasis, rheumatoid arthritis and disease remission. Nat Rev Rheumatol 18, 384–397. 10.1038/s41584-022-00790-8.

135. Hanlon, M.M., Smith, C.M., Canavan, M., Neto, N.G.B., Song, Q., Lewis, M.J., O’Rourke, A.M., Tynan, O., Barker, B.E., Gallagher, P., et al. (2024). Loss of synovial tissue macrophage homeostasis precedes rheumatoid arthritis clinical onset. Sci Adv 10, eadj1252. 10.1126/sciadv.adj1252.

136. Huang, Q.Q., Doyle, R., Chen, S.Y., Sheng, Q., Misharin, A.V., Mao, Q., Winter, D.R., and Pope, R.M. (2021). Critical role of synovial tissue-resident macrophage niche in joint homeostasis and suppression of chronic inflammation. Sci Adv 7. 10.1126/sciadv.abd0515.

137. Alivernini, S., MacDonald, L., Elmesmari, A., Finlay, S., Tolusso, B., Gigante, M.R., Petricca, L., Di Mario, C., Bui, L., Perniola, S., et al. (2020). Distinct synovial tissue macrophage subsets regulate inflammation and remission in rheumatoid arthritis. Nat Med 26, 1295–1306. 10.1038/s41591-020-0939-8.

138. Culemann, S., Gruneboom, A., Nicolas-Avila, J.A., Weidner, D., Lammle, K.F., Rothe, T., Quintana, J.A., Kirchner, P., Krljanac, B., Eberhardt, M., et al. (2019). Locally renewing resident synovial macrophages provide a protective barrier for the joint. Nature 572, 670–675. 10.1038/s41586-019-1471-1.

139. Tsetsarkin, K., Higgs, S., McGee, C.E., De Lamballerie, X., Charrel, R.N., and Vanlandingham, D.L. (2006). Infectious clones of Chikungunya virus (La Reunion isolate) for vector competence studies. Vector Borne Zoonotic Dis 6, 325–337. 10.1089/vbz.2006.6.325.

140. Earnest, J.T., Basore, K., Roy, V., Bailey, A.L., Wang, D., Alter, G., Fremont, D.H., and Diamond, M.S. (2019). Neutralizing antibodies against Mayaro virus require Fc effector functions for protective activity. J Exp Med 216, 2282–2301. 10.1084/jem.20190736.

141. Fox, J.M., Long, F., Edeling, M.A., Lin, H., van Duijl-Richter, M.K.S., Fong, R.H., Kahle, K.M., Smit, J.M., Jin, J., Simmons, G., et al. (2015). Broadly Neutralizing Alphavirus Antibodies Bind an Epitope on E2 and Inhibit Entry and Egress. Cell 163, 1095–1107. 10.1016/j.cell.2015.10.050.

142. Mack, M., Cihak, J., Simonis, C., Luckow, B., Proudfoot, A.E., Plachy, J., Bruhl, H., Frink, M., Anders, H.J., Vielhauer, V., et al. (2001). Expression and characterization of the chemokine receptors CCR2 and CCR5 in mice. J Immunol 166, 4697–4704. 10.4049/jimmunol.166.7.4697.

143. Salyers, A.A., Shoemaker, N., Cooper, A., D’Elia, J., and Shipman, J.A. (1999). Genetic methods for Bacteroides species. Method Microbiol 29, 229–249.

144. Hao, Y., Stuart, T., Kowalski, M.H., Choudhary, S., Hoffman, P., Hartman, A., Srivastava, A., Molla, G., Madad, S., Fernandez-Granda, C., and Satija, R. (2024). Dictionary learning for integrative, multimodal and scalable single-cell analysis. Nat Biotechnol 42, 293–304. 10.1038/s41587-023-01767-y.

145. Hippen, A.A., Falco, M.M., Weber, L.M., Erkan, E.P., Zhang, K., Doherty, J.A., Vaharautio, A., Greene, C.S., and Hicks, S.C. (2021). miQC: An adaptive probabilistic framework for quality control of single-cell RNA-sequencing data. PLoS Comput Biol 17, e1009290. 10.1371/journal.pcbi.1009290.

146. Subramanian, A., Tamayo, P., Mootha, V.K., Mukherjee, S., Ebert, B.L., Gillette, M.A., Paulovich, A., Pomeroy, S.L., Golub, T.R., Lander, E.S., and Mesirov, J.P. (2005). Gene set enrichment analysis: a knowledge-based approach for interpreting genome-wide expression profiles. Proc Natl Acad Sci U S A 102, 15545–15550. 10.1073/pnas.0506580102.

147. Mootha, V.K., Lindgren, C.M., Eriksson, K.F., Subramanian, A., Sihag, S., Lehar, J., Puigserver, P., Carlsson, E., Ridderstrale, M., Laurila, E., et al. (2003). PGC-1alpha-responsive genes involved in oxidative phosphorylation are coordinately downregulated in human diabetes. Nat Genet 34, 267–273. 10.1038/ng1180.

148. Kleverov, M., Zenkova, D., Kamenev, V., Sablina, M., Artyomov, M.N., and Sergushichev, A.A. (2024). Phantasus, a web application for visual and interactive gene expression analysis. Elife 13. 10.7554/eLife.85722.

149. Jin, S., Plikus, M.V., and Nie, Q. (2025). CellChat for systematic analysis of cell-cell communication from single-cell transcriptomics. Nat Protoc 20, 180–219. 10.1038/s41596-024-01045-4.

