## Supplemental Figures for "Gut microbe-derived short-chain fatty acids regulate joint inflammation and activation of infiltrating and resident myeloid cells after alphavirus infection"

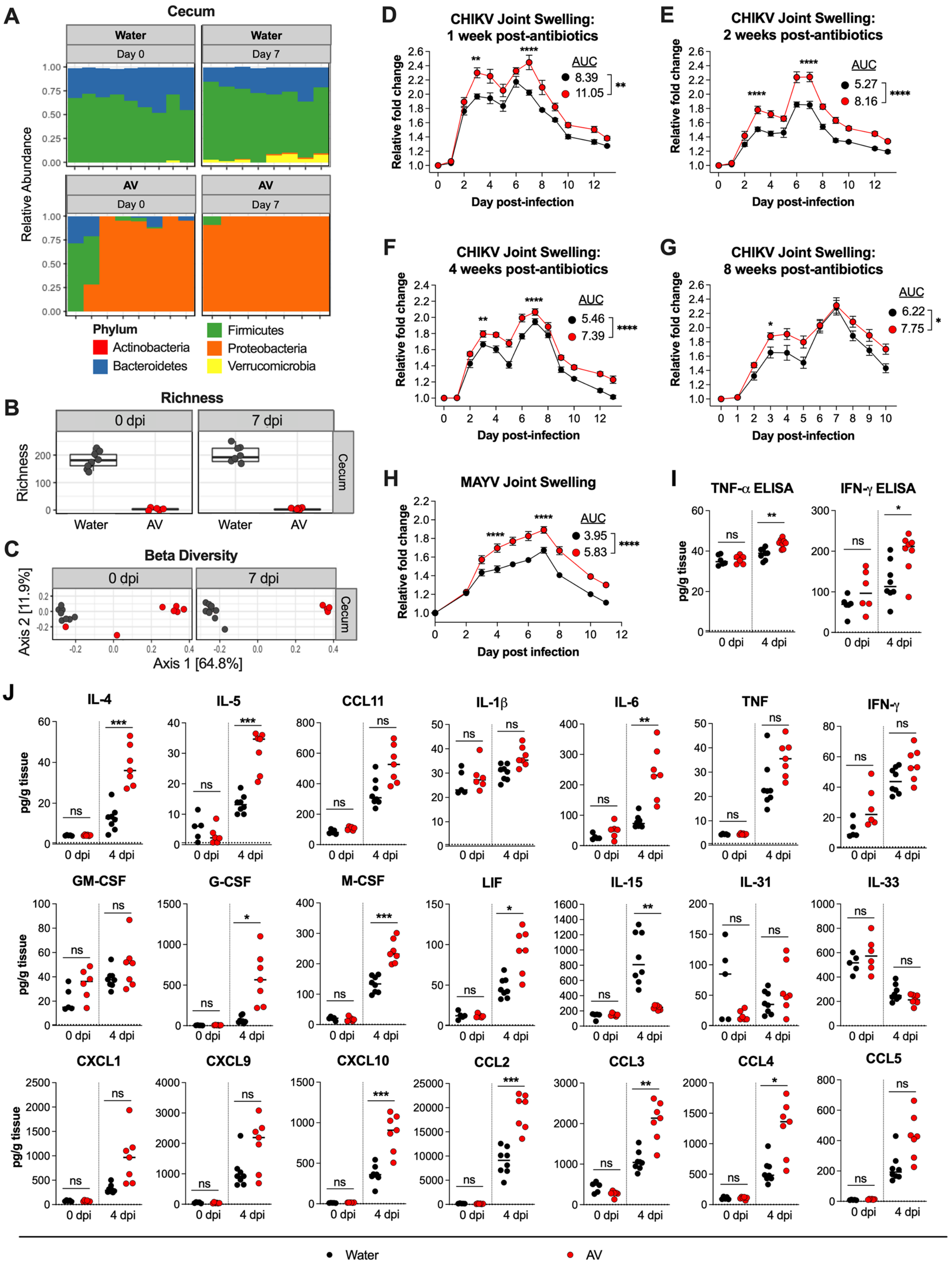
A
Cecum
Water
Day 0
Water
Day 7
AV
Day 0
AV
Day 7
Relative Abundance
1.00
0.75
0.50
0.25
0.00
Phylum
Actinobacteria
Bacteroidetes
Firmicutes
Proteobacteria
Verrucomicrobia
B
Richness
0 dpi
7 dpi
Cecum
Richness
200
100
0
Water
AV
Water
AV
C
Beta Diversity
0 dpi
7 dpi
Cecum
Axis 2 [11.9%]
Axis 1 [64.8%]
D
CHIKV Joint Swelling:
1 week post-antibiotics
Relative fold change
Day post-infection
AUC
8.39
11.05
**
****
E
CHIKV Joint Swelling:
2 weeks post-antibiotics
Relative fold change
Day post-infection
AUC
5.27
8.16
****
F
CHIKV Joint Swelling:
4 weeks post-antibiotics
Relative fold change
Day post-infection
AUC
5.46
7.39
****
**
G
CHIKV Joint Swelling:
8 weeks post-antibiotics
Relative fold change
Day post-infection
AUC
6.22
7.75
*
H
MAYV Joint Swelling
Relative fold change
Day post infection
AUC
3.95
5.83
****
I
TNF-α ELISA
IFN-γ ELISA
pg/g tissue
0 dpi
4 dpi
ns
**
*
J
IL-4
IL-5
CCL11
IL-1β
IL-6
TNF
IFN-γ
GM-CSF
G-CSF
M-CSF
LIF
IL-15
IL-31
IL-33
CXCL1
CXCL9
CXCL10
CCL2
CCL3
CCL4
CCL5
pg/g tissue
0 dpi
4 dpi
ns
*
**
***
Water
AV

**Figure S1. Oral antibiotic treatment alters bacterial composition in cecal and colonic contents and has a sustained effect in exacerbating alphavirus-induced foot swelling.** (**A-C**) Cecal contents were collected from water- or AV-treated mice at 0 or 7 dpi. (**A**) Relative abundance of bacterial phyla detected (bacterial phyla represented in less than 1% per sample were removed), (**B**) number of bacterial taxa (richness), and (**C**) bacterial beta diversity (weighted UniFrac distance) of cecal contents (2 experiments, n = 8-9 mice per group). (**D-G**) Foot swelling in water- or AV-treated mice after CHIKV infection at (**D**) 1, (**E**) 2, (**F**) 4, or (**G**) 8 weeks post cessation of antibiotics (2 experiments, n = 7-8 mice per group). (**H**) Foot swelling in water- or AV-treated mice after MAYV infection (2 experiments, n = 9-10 per group). (**I**) TNF and IFN-$\gamma$ levels in joint homogenates from water- and AV-treated mice at 0 and 4 dpi, as measured by ELISA (n = 5-8 per group). (**J**) Cytokine and chemokine levels in joint homogenates from water and AV-treated mice at 0 and 4 dpi, as measured by BioPlex assay (n = 5-8 per group). Statistical analysis: **B**, Wilcoxon tests; **C**, permutational multivariate analysis of variance (ADONIS); **D-H**, mean ± SEM, two-way ANOVA with Šídák's post-test, or unpaired t-test for analysis of AUC; **I**, unpaired t-test; **J**, unpaired t-test with Bonferroni correction for multiple comparisons. **** $P < 0.0001$; *** $P < 0.001$; ** $P < 0.01$; * $P < 0.05$; ns, not significant.

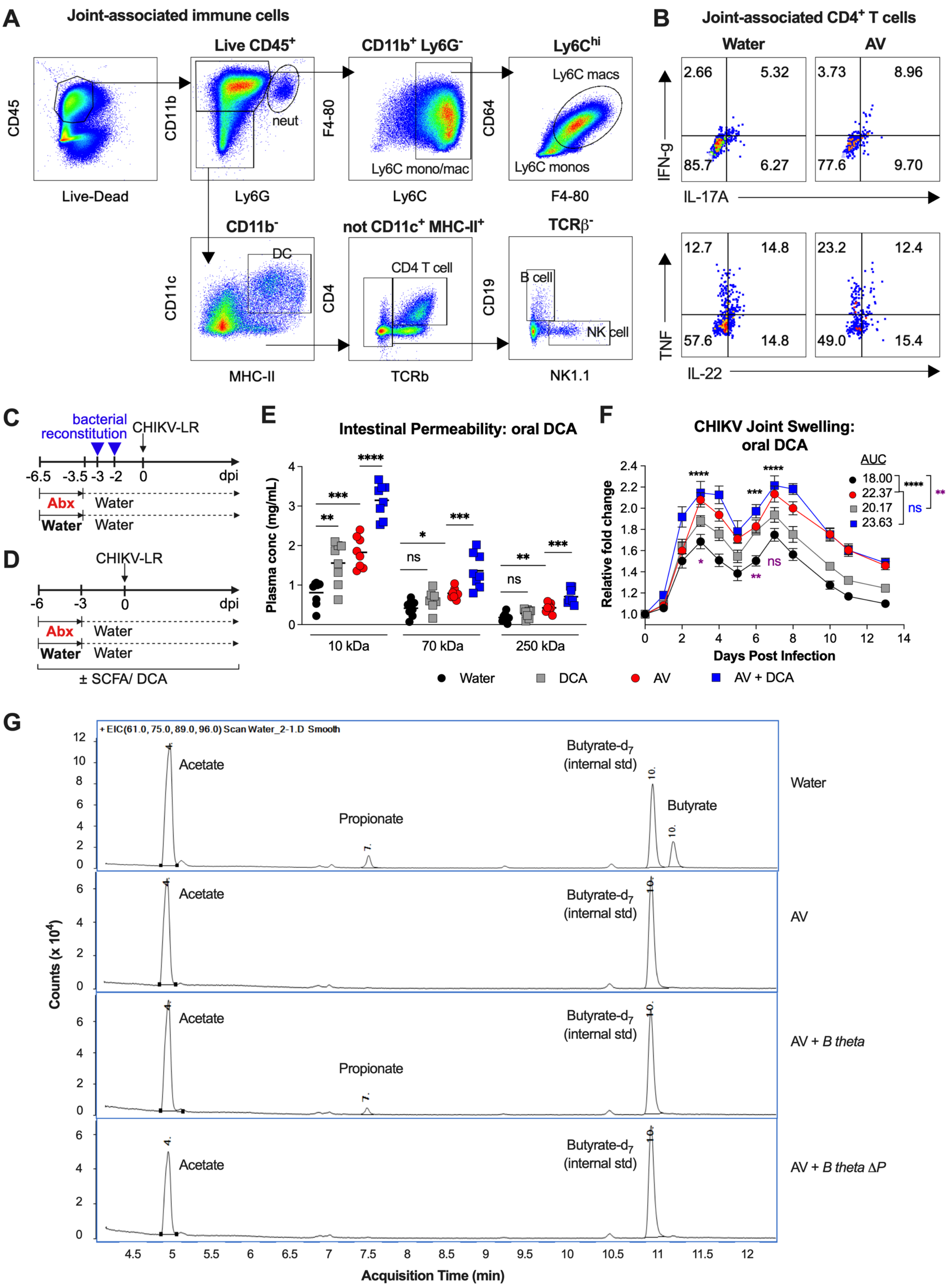

A
Joint-associated immune cells
CD45
Live-Dead
Live CD45+
CD11b
Ly6G
neut
CD11b+ Ly6G-
F4-80
Ly6C
Ly6C mono/mac
Ly6Chi
CD64
F4-80
Ly6C macs
Ly6C monos
CD11b-
CD11c
MHC-II
DC
not CD11c+ MHC-II+
CD4
TCRb
CD4 T cell
TCRβ-
CD19
NK1.1
B cell
NK cell
B
Joint-associated CD4+ T cells
Water
AV
IFN-g
IL-17A
2.66
5.32
85.7
6.27
3.73
8.96
77.6
9.70
TNF
IL-22
12.7
14.8
57.6
14.8
23.2
12.4
49.0
15.4
C
bacterial reconstitution
CHIKV-LR
-6.5 -3.5 -3 -2 0 dpi
Abx Water
Water Water
D
CHIKV-LR
-6 -3 0 dpi
Abx Water
Water Water
± SCFA/ DCA
E
Intestinal Permeability: oral DCA
Plasma conc (mg/mL)
10 kDa
70 kDa
250 kDa
F
CHIKV Joint Swelling: oral DCA
Relative fold change
Days Post Infection
AUC
18.00
22.37
20.17
23.63
ns
Water
DCA
AV
AV + DCA
G
+ EIC(61.0, 75.0, 89.0, 96.0) Scan Water_2-1.D Smooth
Acetate
Propionate
Butyrate-d7 (internal std)
Butyrate
Water
AV
AV + B theta
AV + B theta ΔP
Counts (x 10^4)
Acquisition Time (min)

**Figure S2. Gating scheme of immune cell subsets and oral metabolite supplementation.** (**A**) Gating scheme for immune cell subsets in joint-associated tissues after CHIKV infection. (**B**) Representative flow cytometry plots showing intracellular cytokine staining in $CD4^+$ T cells from joint-associated tissues of water or AV-treated mice at 4 dpi. (**C-D**) Schematics of treatments for mice that received (**C**) bacterial reconstitution or (**D**) exogenous supplementation with either individual SCFAs (butyrate, propionate, or acetate) or DCA in drinking water. (**E-F**) Mice were treated with water or AV, with or without DCA supplementation, and subsequently inoculated with CHIKV. (**E**) Intestinal permeability as measured by plasma concentrations of 10 kDa, 70 kDa, and 250 kDa dextran at 1.5 h after oral gavage (2 experiments, n = 8-9 per group). (**F**) Foot swelling after CHIKV infection in water or AV-treated mice receiving either water or SCFA supplementation with DCA (2 experiments, n = 8 per group). (**G**) Representative chromatograms showing intensity (counts) by retention time of processed cecal samples from control, AV-treated, and AV-treated mice colonized with either wild-type or mutant (*ΔP*) *B. thetaiotaomicron*. Statistical analysis: **E**, one-way ANOVA with Šídák's multiple comparisons test. **F**, mean ± SEM, two-way ANOVA with Šídák's post-test, or one-way ANOVA with Dunnett's multiple comparisons for analysis of AUC. **** $P < 0.0001$; *** $P < 0.001$; ** $P < 0.01$; * $P < 0.05$; ns, not significant.

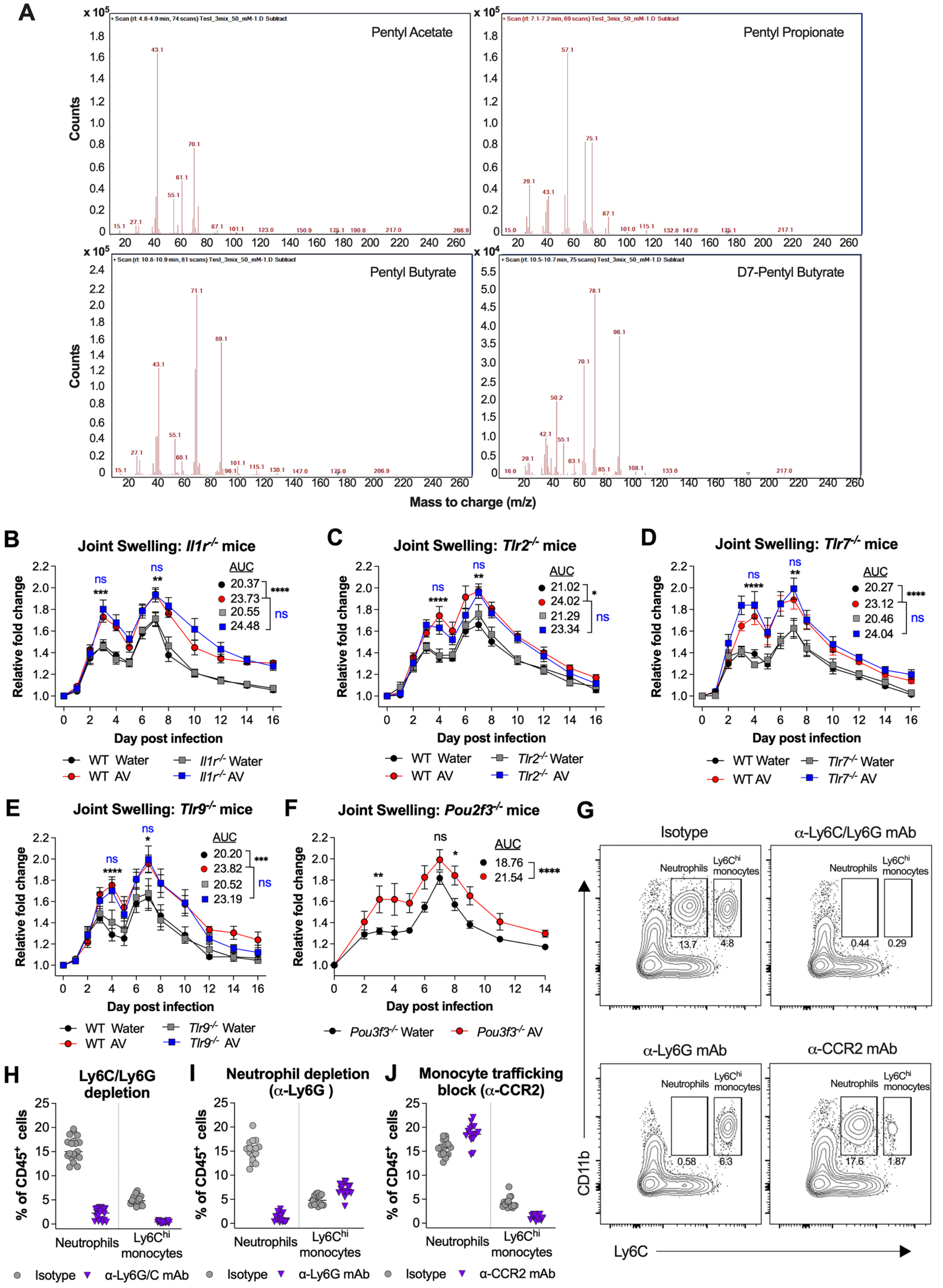
A
Pentyl Acetate
Pentyl Propionate
Pentyl Butyrate
D7-Pentyl Butyrate
Counts
Mass to charge (m/z)
B
Joint Swelling: Il1r-/- mice
C
Joint Swelling: Tlr2-/- mice
D
Joint Swelling: Tlr7-/- mice
E
Joint Swelling: Tlr9-/- mice
F
Joint Swelling: Pou2f3-/- mice
Relative fold change
Day post infection
AUC
20.37
23.73
20.55
24.48
21.02
24.02
21.29
23.34
20.27
23.12
20.46
24.04
20.20
23.82
20.52
23.19
18.76
21.54
WT Water
WT AV
Il1r-/- Water
Il1r-/- AV
Tlr2-/- Water
Tlr2-/- AV
Tlr7-/- Water
Tlr7-/- AV
Tlr9-/- Water
Tlr9-/- AV
Pou3f3-/- Water
Pou3f3-/- AV
G
Isotype
α-Ly6C/Ly6G mAb
α-Ly6G mAb
α-CCR2 mAb
Neutrophils
Ly6Chi monocytes
13.7
4.8
0.44
0.29
0.58
6.3
17.6
1.87
CD11b
Ly6C
H
Ly6C/Ly6G depletion
I
Neutrophil depletion (α-Ly6G )
J
Monocyte trafficking block (α-CCR2)
% of CD45+ cells
Neutrophils
Ly6Chi monocytes
Isotype
α-Ly6G/C mAb
α-Ly6G mAb
α-CCR2 mAb

**Figure S3. Increased CHIKV-induced joint swelling after antibiotics treatment is dependent on T cells but not affected by loss of IL1R, TLR2, TLR7, or TLR9 signaling or by an absence of Tuft cells.** (**A**) Mass spectra showing intensity by the mass to charge ratio for the indicated standards. (**B-E**) Foot swelling in water or AV-treated (**B**) $Il1r^{-/-}$, (**C**) $Tlr2^{-/-}$, (**D**) $Tlr7^{-/-}$, or (**E**) $Tlr9^{-/-}$ and WT littermate mice after CHIKV infection (2 to 5 experiments, n = 7-24 per group). (**F**) Foot swelling after CHIKV infection in water or AV-treated $Pou2f3^{-/-}$ mice (2 experiments, n = 6-9 per group). (**G**) Representative flow cytometry plots of monocytes and neutrophils from peripheral blood of mice treated with isotype control, anti-Ly6C/LyG, anti-Ly6G, or anti-CCR2 mAbs. (**H-J**) Frequency of neutrophils and $Ly6C^{hi}$ monocytes in peripheral blood following depletion with isotype control or mAbs against (**H**) Ly6G/Ly6G (GR-1), (**I**) ly6G, and (**J**) CCR2. Statistical analysis: **B-F,** mean ± SEM, two-way ANOVA with Dunnett's multiple comparisons (**B-E**) or Šídák's post-test (**F**); for analysis of AUC, one-way ANOVA with Dunnett's multiple comparisons (**B-E**) or unpaired t-test (**F**). **** $P < 0.0001$; *** $P < 0.001$; ** $P < 0.01$; * $P < 0.05$; ns, not significant.

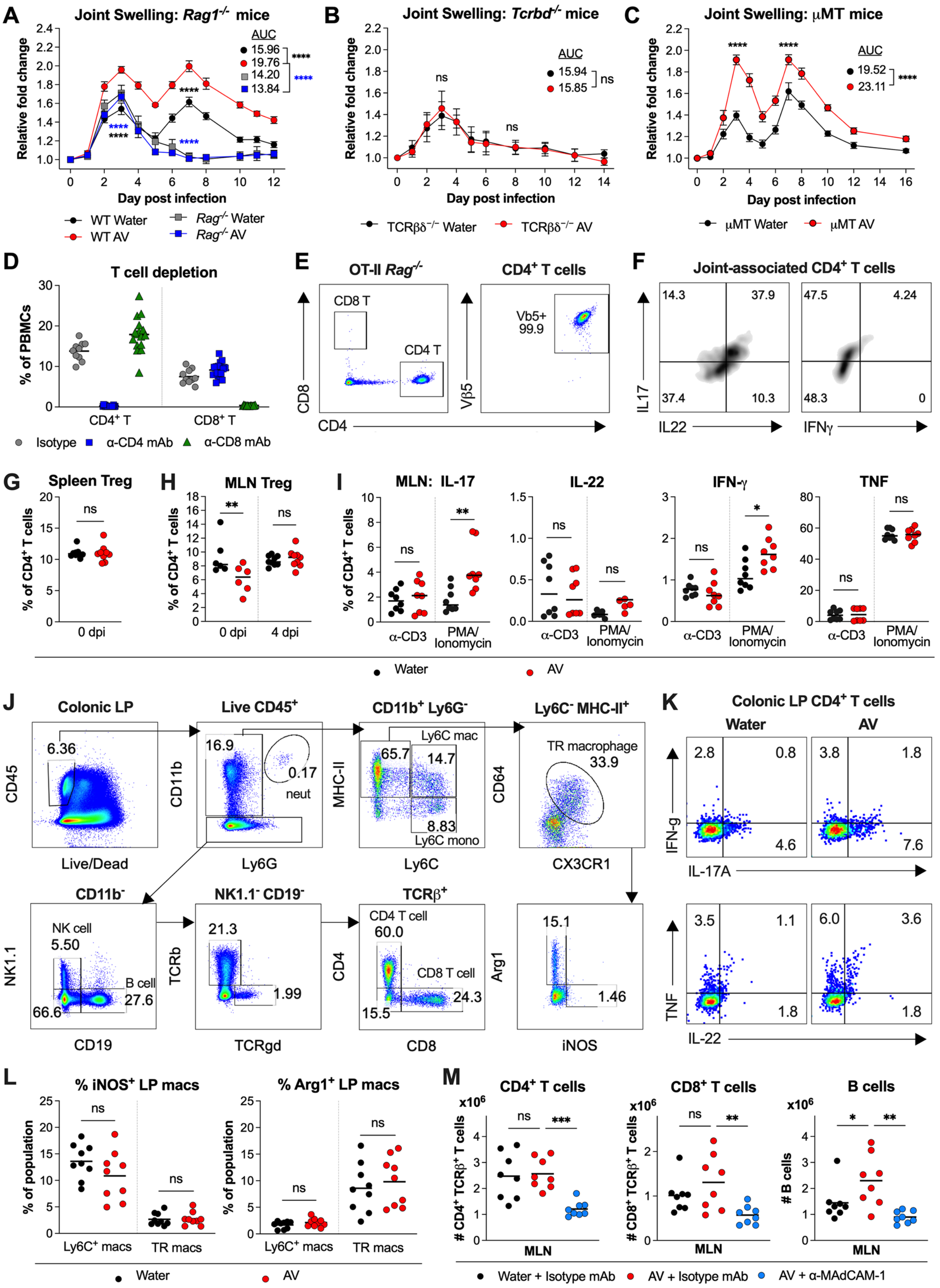

A
Joint Swelling: Rag1-/- mice
AUC
15.96
19.76
14.20
13.84
****
Relative fold change
Day post infection
WT Water
WT AV
Rag-/- Water
Rag-/- AV
B
Joint Swelling: Tcrbd-/- mice
AUC
15.94
15.85
ns
TCRβδ-/- Water
TCRβδ-/- AV
C
Joint Swelling: μMT mice
AUC
19.52
23.11
μMT Water
μMT AV
D
T cell depletion
% of PBMCs
CD4+ T
CD8+ T
Isotype
α-CD4 mAb
α-CD8 mAb
E
OT-II Rag-/-
CD4+ T cells
CD8 T
CD4 T
Vb5+
99.9
CD8
CD4
Vβ5
F
Joint-associated CD4+ T cells
14.3
37.9
37.4
10.3
47.5
4.24
48.3
0
IL17
IL22
IFNγ
G
Spleen Treg
% of CD4+ T cells
0 dpi
H
MLN Treg
4 dpi
I
MLN: IL-17
IL-22
IFN-γ
TNF
α-CD3
PMA/
Ionomycin
Water
AV
J
Colonic LP
Live CD45+
CD11b+ Ly6G-
Ly6C- MHC-II+
6.36
16.9
0.17
neut
Ly6C mac
65.7
14.7
8.83
Ly6C mono
TR macrophage
33.9
CD45
Live/Dead
CD11b
Ly6G
MHC-II
Ly6C
CD64
CX3CR1
CD11b-
NK1.1- CD19-
TCRβ+
NK cell
5.50
B cell
27.6
66.6
21.3
1.99
CD4 T cell
60.0
CD8 T cell
24.3
15.5
15.1
1.46
NK1.1
CD19
TCRb
TCRgd
CD4
CD8
Arg1
iNOS
K
Colonic LP CD4+ T cells
Water
AV
2.8
0.8
4.6
3.8
1.8
7.6
IFN-g
IL-17A
3.5
1.1
1.8
6.0
3.6
1.8
TNF
IL-22
L
% iNOS+ LP macs
% Arg1+ LP macs
% of population
Ly6C+ macs
TR macs
Water
AV
M
CD4+ T cells
CD8+ T cells
B cells
x10^6
### CD4+ TCRβ+ T cells
### CD8+ TCRβ+ T cells
### B cells
MLN
Water + Isotype mAb
AV + Isotype mAb
AV + α-MAdCAM-1

**Figure S4. Splenic, mesenteric lymph node, and lamina propria T cells in water- or antibiotic-treated mice.** (**A-C**) Foot swelling in water or AV-treated (**A**) *Rag*$^{-/-}$ (2 experiments, n = 9-10 per group), (**B**) *Tcrbd*$^{-/-}$ (2 experiments, n = 7-8 per group), or (**C**) μMT mice (2 experiments, n = 9 per group) after CHIKV infection. (**D)** Frequency of CD4$^{+}$ and CD8$^{+}$ T cells in the peripheral blood of mice following depletion with mAbs against CD4 and CD8 (2-3 experiments, n = 8-16 per group). (**E-F**) Representative flow cytometry plots showing (**E**) CD4$^{+}$ and CD8$^{+}$ T cells in the spleens of OT-II *Rag*$^{-/-}$ mice (virtually all CD4$^{+}$ T cells express the transgenic, MHC-II restricted, TCR [Vα2/Vβ5] against chicken ovalbumin) and (**F**) production of the indicated cytokines by CD4$^{+}$ T cells from the joint tissue of OT-II mice. (**G-H**) Percentages of CD25$^{+}$ FoxP3$^{+}$ Tregs of total CD4$^{+}$ T cells in the (**G**) spleen and (**H**) mesenteric lymph nodes (MLN) of water- or AV-treated mice at the indicated timepoints (2 experiments, n = 6-9 per group). (**I**) Percentages of CD4$^{+}$ T cells from the MLNs of water- or AV-treated mice that produce IL-17, IL-22, and IFN-γ in response to *ex vivo* stimulation with anti-CD3 mAb or PMA/ionomycin (2 experiments, n = 8 per group). (**J**) Gating scheme for immune cell subsets in the colonic lamina propria. (**K**) Representative flow cytometry plots showing intracellular cytokine staining in CD4$^{+}$ T cells from the colonic lamina propria of water- or AV-treated mice. (**L**) Percentages of iNOS- and Arg1-expressing macrophages in the colonic lamina propriat of water- or AV-treated mice. (**M**) Numbers of the indicated immune cells in the mesenteric lymph nodes of mice that received either isotype control or mAbs against MAdCAM-1 prior to treatment with water or AV and subsequent CHIKV infection (2 experiments, n = 8 per group). Statistical analysis: **A-C**, mean ± SEM, two-way ANOVA with Dunnett's multiple comparisons (**A**) or Šídák's post-test (**B-C**); for analysis of AUC, one-way ANOVA with Dunnett's multiple comparisons (**A**) or unpaired t-test (**B-C**). **G-I, L,** unpaired t-test. **M,** One-way ANOVA with Dunnett's multiple comparisons test. **** $P < 0.0001$; *** $P < 0.001$; ** $P < 0.01$; * $P < 0.05$; ns, not significant.

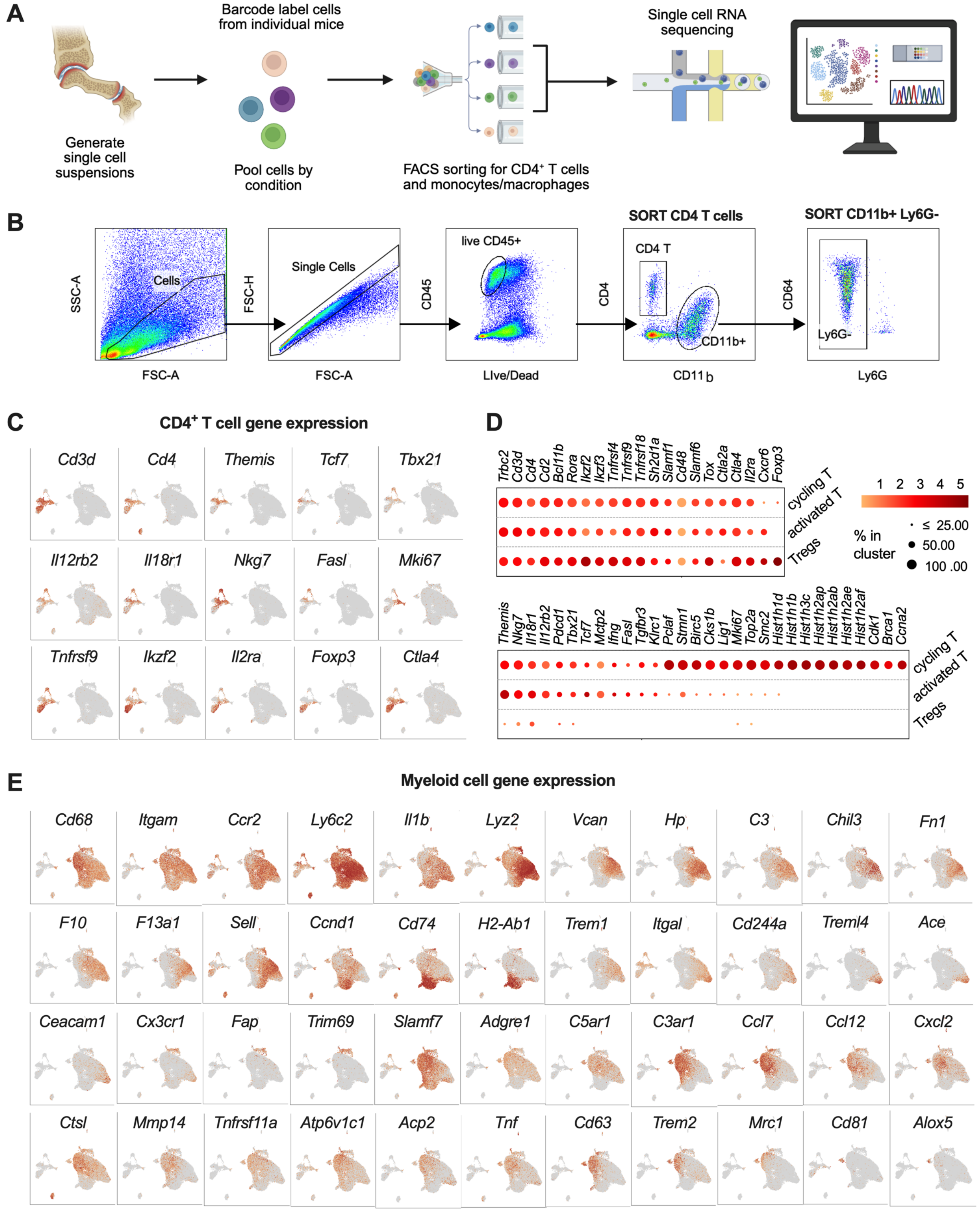


**Figure S5. Gene expression in joint-associated CD4⁺ T and myeloid cells.** (**A**) Schematic of experimental design for single cell transcriptomic analysis of joint-associated CD4⁺ T cells, monocytes, and macrophages. (**B**) Gating for flow cytometry sort enrichment of joint-associated CD4⁺ T cells, monocytes, and macrophages for single cell RNA sequencing. (**C**) UMAP visualization of enriched CD4$^+$ T and myeloid cells from single cell-RNA sequencing data, showing expression of the indicated genes in CD4$^+$ T cells. (**D**) Bubble plots showing patterns of gene expression amongst CD4$^+$ Tregs, activated T cells, and proliferating T cells. (**E**) UMAP visualization of enriched CD4$^+$ T and myeloid cells from single cell-RNA sequencing data showing expression of the indicated genes.
